# Disentangling cell-intrinsic and extrinsic factors underlying evolution

**DOI:** 10.1101/2024.05.06.592777

**Authors:** Alexander L. Starr, Toshiya Nishimura, Kyomi J. Igarashi, Chihiro Funamoto, Hiromitsu Nakauchi, Hunter B. Fraser

**Author notes:** Indicates corresponding author. These authors contributed equally to this work (Alexander L. Starr, Toshiya Nishimura).

## Abstract

A key goal of developmental biology is to determine the extent to which cells and organs develop autonomously, as opposed to requiring interactions with other cells or environmental factors. Chimeras have played a foundational role in this by enabling qualitative classification of cell-intrinsically vs. extrinsically driven processes. Here, we extend this framework to precisely decompose evolutionary divergence in any quantitative trait into cell-intrinsic, extrinsic, and intrinsic-extrinsic interaction components. Applying this framework to thousands of gene expression levels in reciprocal rat-mouse chimeras, we found that the majority of their divergence is attributable to cell-intrinsic factors, though extrinsic factors also play an integral role. For example, a rat-like extracellular environment extrinsically up-regulates the expression of a key transcriptional regulator of the endoplasmic reticulum (ER) stress response in some but not all cell types, which in turn strongly predicts extrinsic up-regulation of its target genes and of the ER stress response pathway as a whole. This effect is also seen at the protein level, suggesting propagation through multiple regulatory levels. Applying our framework to a cellular trait, neuronal differentiation, revealed a complex interaction of intrinsic and extrinsic factors. Finally, we show that imprinted genes are dramatically mis-expressed in species-mismatched environments, suggesting that mismatch between rapidly evolving intrinsic and extrinsic mechanisms controlling gene imprinting may contribute to barriers to interspecies chimerism. Overall, our conceptual framework opens new avenues to investigate the mechanistic basis of developmental processes and evolutionary divergence across myriad quantitative traits in any multicellular organism.

## Introduction

Soon after the founding of the field of experimental embryology in the 19th century, one of its pioneers emphasized that its central goal was distinguishing to what extent development proceeds via “selbstdifferenzierung” (self-differentiation, now known as cell-autonomous or cell-intrinsic), as opposed to “abhängige differenzierung” (dependent differentiation, now known as non-cell-autonomous or cell-extrinsic)^1,2^. Since then, many seminal discoveries have been made by studying this dichotomy^3–9^. Interspecies chimeras—formed by grafting donor cells from one species into the host embryo of another—have fueled many of these key insights for over a century^3–7^. In general, a process is said to be cell-intrinsic if the cells responsible for executing that process behave in a similar way regardless of the genetic background of surrounding cells. For example, quail donor cranial neural crest cells destined to form the beak follow their faster cell-intrinsic developmental timeline and execute their own spatial and temporal programs for gene expression when they are transplanted into a slower-developing duck host embryo^6,10,11^.

On the other hand, a process is cell-extrinsic when dependent on the genotype of other cells; duck host epidermal cells in the same quail-duck chimeras are extrinsically induced by the quail donor neural crest to alter their gene expression to make quail-like feathers and a conical egg tooth^6,10,11^. Although this categorization of developmental processes into extrinsically vs. intrinsically controlled has revealed central principles in developmental and evolutionary biology, there is currently no general framework to quantitatively deconvolve traits into cell-extrinsic and intrinsic components.

In contrast, the quantitative dissection of two fundamentally different molecular mechanisms to divergence in gene expression, a key driver of evolutionary adaptation and phenotypic novelty, has been extensively explored^12,13^. These mechanisms are known as *cis* and *trans*: *cis* refers to regulatory elements such as promoters and enhancers that affect nearby genes on the same chromosome, whereas *trans*-acting factors involve diffusible molecules such as transcription factors (TFs) or noncoding RNAs that can regulate genes throughout the genome^13^. A landmark study used *Drosophila* interspecies hybrids to disentangle the *cis*- and *trans*-regulatory contributions to gene expression divergence^13^. Since then, genome-wide *cis*- and *trans*- regulatory variation has been estimated for species across the tree of life, finding that the majority of variation within most species is caused by *trans*-acting mechanisms^14–17^. While *cis*- acting divergence is always cell-intrinsic, both intrinsic and extrinsic factors can lead to *trans*- acting divergence, yet their relative contributions to phenotypic evolution remain unknown.

Here, we unify these concepts and introduce a framework to disentangle the contributions of intrinsic factors, extrinsic factors, and their interactions to evolutionary divergence in gene expression and other quantitative traits.

## Results

### Cell-extrinsic and intrinsic divergence in interspecies chimeras: concepts

A chimera is an amalgamation of two sets of cells, donor and host. We define host cells as the cells derived from the injected blastocyst; when host cells make up a large majority, they are in species-matched environments similar to wildtype organisms (Fig. 1A). Donor cells are the progeny of the cells that were injected into the blastocyst; when these are a small minority, they are in species-mismatched environments determined by the host (Fig. 1B-C). Throughout, we use donor cells/cells in species-mismatched environments interchangeably and host cells/cells in species-matched environments interchangeably.

**Fig. 1:**
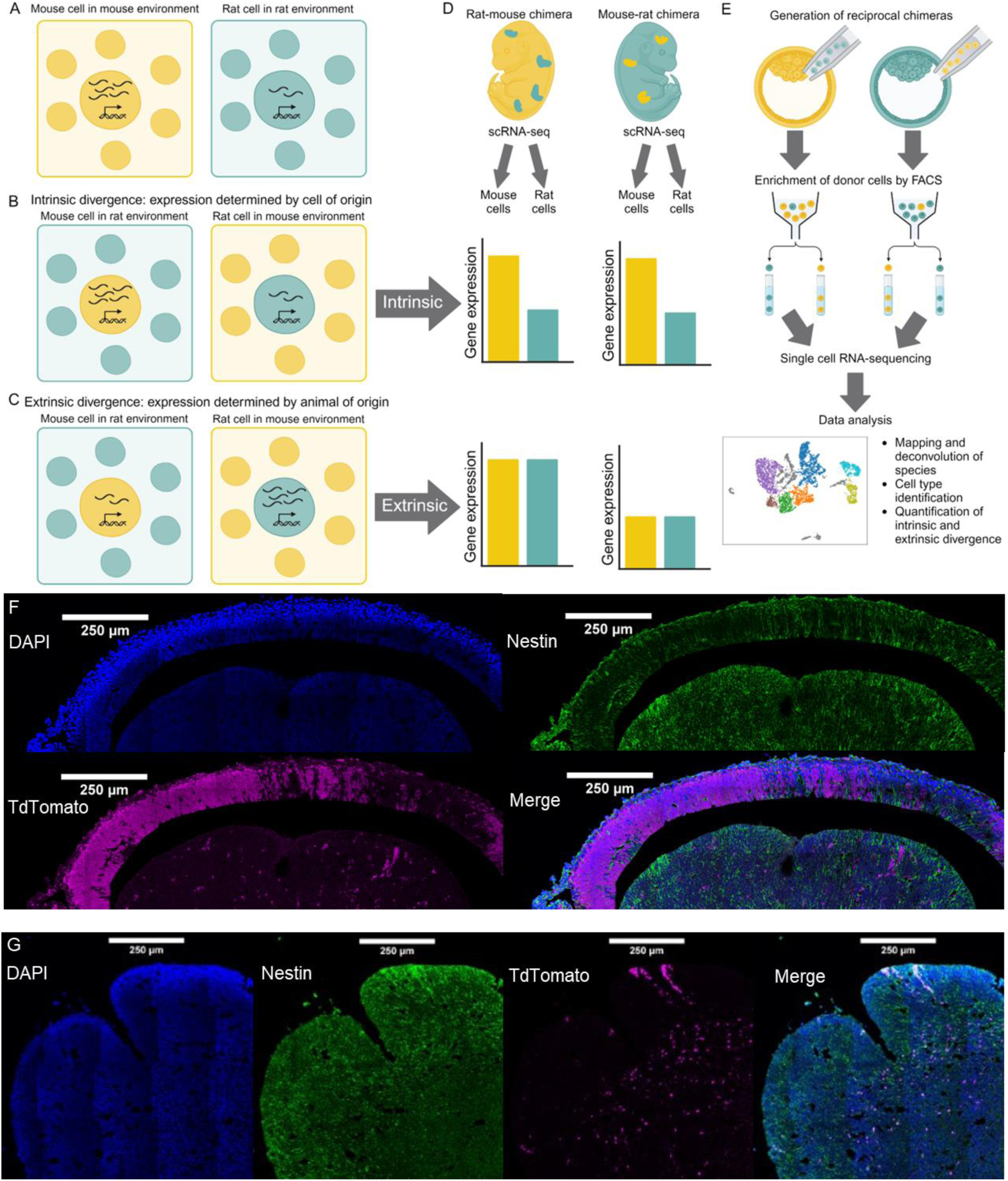
Conceptual model of estimating extrinsic and intrinsic divergence using reciprocal chimeras. **A)** Rat and mouse cells in species-matched environments. In wildtype animals, the divergence in expression of a single gene between species is determined by divergence intrinsic to mouse and rat cells, as well as divergence in the extracellular environment. In this example, expression is 2.5-fold higher in mouse cells compared to rat cells when both cells are in species-matched environments. However, it is unclear how much of this divergence is extrinsically driven or intrinsically driven. **B)** Example of cell-intrinsic divergence. In interspecies chimeras, we can measure gene expression in mouse cells in a rat-like environment and in rat cells in a mouse-like environment. If the expression of the gene remains unchanged compared to species-matched environments, then the divergence in gene expression must be due to intrinsic mechanisms. **C)** Example of cell-extrinsic divergence. If the expression of this gene in mouse cells in a rat-like environment matches the expression of the gene in rat cells in a rat-like environment and vice versa, then the divergence in gene expression must be due to extrinsic divergence in the extracellular environment. **D)** Outline of measuring intrinsic and extrinsic divergence genome-wide. To measure extrinsic and intrinsic divergence, we can generate mouse-like (a small proportion of rat cells and a large proportion of mouse cells) and rat-like (a small proportion of mouse cells and a large proportion of rat cells) reciprocal chimeras. We can then use scRNA-seq to measure gene expression in each cell type across the four different species-environment combinations. If the divergence in expression for a gene is similar in both rat-like and mouse-like environments, then it is driven by intrinsic divergence. On the other hand, if expression is similar when mouse and rat cells are in the same extracellular environment but different when comparing cells from two different environments, it is driven by extrinsic divergence. **E)** Outline of experimental procedure for this study. Reciprocal chimeras were generated via blastocyst injection. Cells from mouse-like chimeras were harvested at E13.5 and cells from rat-like chimeras were harvested at a matched developmental stage (E15.5). Donor cells in species-mismatched environments were enriched using fluorescence-activated cell sorting using TdTomato as a marker and single-cell RNA-sequencing was performed. Each cell was then computationally identified as mouse or rat based on RNA-seq reads, clustered, and used to estimate extrinsic and intrinsic divergence between species. **F)** Immunofluorescence image of the E13.5 forebrain of a mouse-like chimera. Nuclei are stained with DAPI (blue) and Nestin (a neural stem cell marker) is shown in green. TdTomato (magenta) marks donor rat cells in the mouse-like environment. **G)** Immunofluorescence image of the E15.5 ganglionic eminence of a rat-like chimera. In this case, TdTomato marks mouse cells in the rat-like environment.

To illustrate the concept of extrinsic and intrinsic divergence in gene expression, we can consider an orthologous gene that is more highly expressed in mice than in rats. If mouse cells express the gene at the same level regardless of their extracellular environment, and likewise for rat cells, then we infer that the expression difference is determined by the cell of origin rather than its extracellular environment, suggesting cell-intrinsic divergence (Fig. 1B). However, if donor and host cells express this gene at equal levels within each chimera, then we infer that this divergence is determined by the extracellular environment of the cells in that individual, suggesting extrinsic divergence in gene expression caused by divergence in the extracellular environment (Fig. 1C). In practice, we can measure the intrinsic and extrinsic divergence in gene expression using single-cell RNA-seq (scRNA-seq) of reciprocal interspecies chimeras (Fig. 1D).

### Cell-extrinsic and intrinsic divergence in interspecies chimeras: data

To explore this concept empirically, we first generated reciprocal rat-like and mouse-like chimeras via blastocyst injection (Fig. 1E, Methods). Importantly, we only used tissues with less than 10% donor contribution so that the extrinsic environment was primarily composed of host cells. As an example, we observed clear populations of host and donor cells (marked by TdTomato) in the forebrain of both mouse-like and rat-like chimeras (Fig. 1F-G). Consistent with previous work, different brain regions showed distinct patterns, as donor cells were rarer and more evenly distributed in the ganglionic eminence compared to the developing neocortex in mouse-like chimeras (Fig. 1G)^18^.

Next, we enriched for donor cells from stage-matched embryos (E13.5 in mouse, E15.5 in rat), then generated and analyzed scRNA-seq data from the forebrain and connective tissue/spinal cord of both mouse-like (i.e. made up primarily of mouse host cells) and rat-like chimeras (Fig. 1E, Figs. S1-S2, Methods). After removing low-quality cells, identifying cell types and filtering those with fewer than 10 cells in any sample type (host mouse, host rat, donor mouse, and donor rat), we retained 4,720 cells distributed across 11 cell types.

### Cell-extrinsic and intrinsic divergence in interspecies chimeras: a quantitative framework

With this dataset in hand, we developed a quantitative framework to estimate the extrinsic, intrinsic, and intrinsic-extrinsic interaction components of gene expression divergence for each gene (Fig. 2, Methods). In the absence of intrinsic-extrinsic interaction, the effect of the extracellular environment on mouse and rat cells is identical, enabling us to obtain two independent estimates of intrinsic divergence (Fig. 2A). On the other hand, when comparing cells from the same species in different extracellular environments, the intrinsic contribution of the cells of that species is identical, enabling us to obtain two estimates of extrinsic divergence (Fig. 2B). Any difference between these two estimates must be due to either measurement error or an interaction between extrinsic and intrinsic divergence (which is defined as extrinsic divergence in the host environments differing based on the genotype of the cell being measured, and/or intrinsic divergence between cells of different genotypes differing based on the genotype of the host; see below and Methods). Thus, we can use the four measurements obtained from reciprocal chimeras to decompose divergence in any quantitative trait into the intrinsic, extrinsic, and interaction components.

**Fig. 2:**
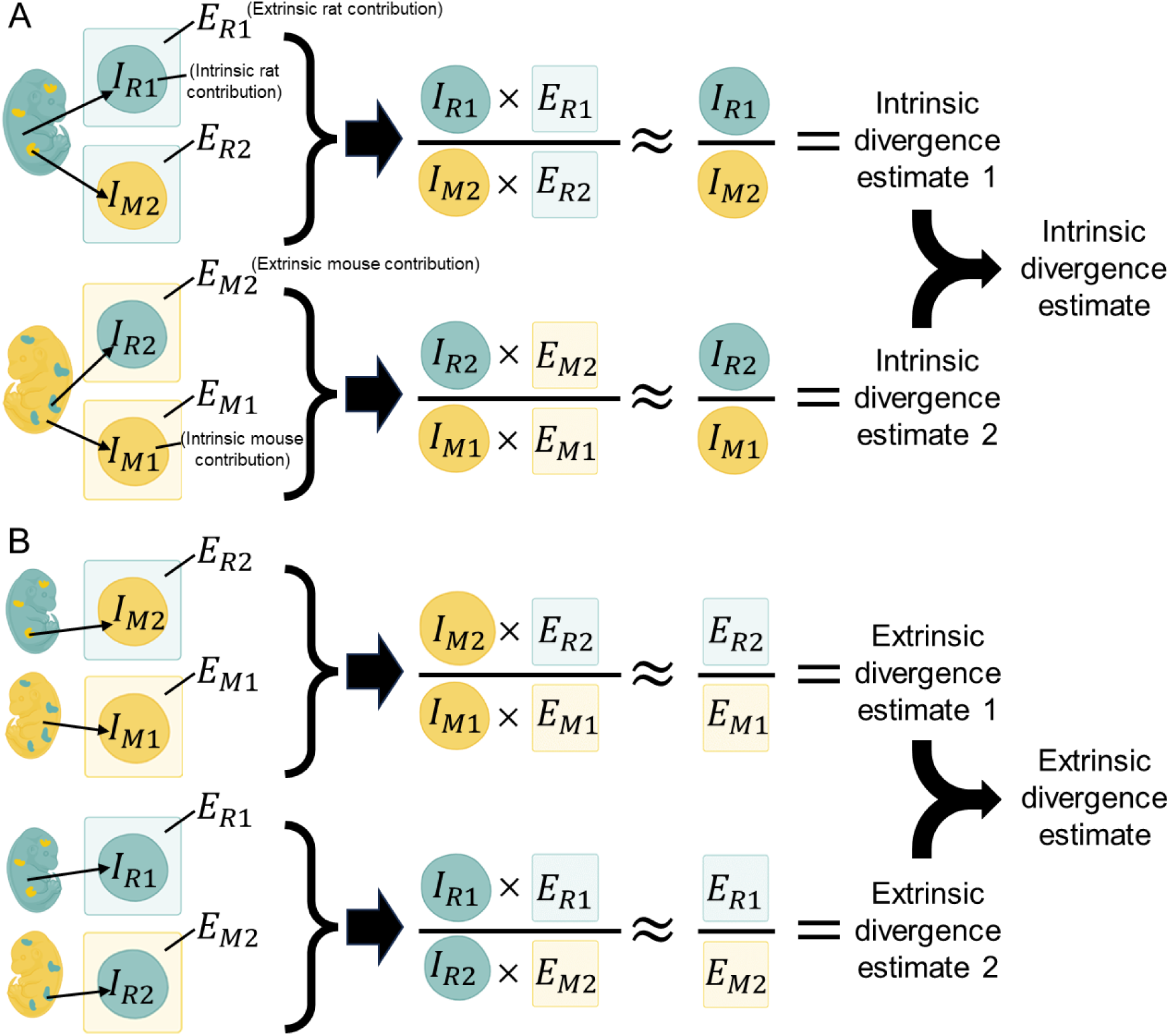
Quantitative framework for measuring intrinsic and extrinsic divergence. **A)** Quantitative framework for measuring intrinsic divergence. In the absence of noise and interaction between intrinsic and extrinsic components, the effect of the rat-like environment on mouse and rat cells is identical such that this extrinsic contribution cancels out, leaving only an estimate of the intrinsic divergence (top). Similarly, we can obtain another estimate of intrinsic divergence using the mouse-like chimeras, averaging the two estimates to obtain the final estimate of intrinsic divergence. Subscript numbers denote our two independent estimates of the same underlying quantity, which may differ from each other due to measurement error and/or intrinsic-extrinsic interactions. **B)** Quantitative framework for measuring extrinsic divergence. In the absence of noise and interaction between intrinsic and extrinsic components, the intrinsic contribution of mouse cells is identical in a mouse and rat environment, such that this intrinsic contribution cancels out leaving only an estimate of the intrinsic divergence (top). Similarly, we can obtain another estimate of extrinsic divergence using the other two measurements, averaging the two estimates to obtain the final estimate of intrinsic divergence. See methods for additional details and estimation of intrinsic-extrinsic interaction.

As an example, we can consider the expression of *Efnb3* in forebrain glutamatergic neurons (abbreviated as brain.glut.neu). In species-matched environments, we observed two-fold higher expression in mouse than in rat (Fig. 3A, left; the bars to the left of the dashed line are the same data as the bars for species-matched environments to the right of the dashed line and are included twice for clarity). However, in both chimeras, *Efnb3* expression in the donor cells mirrored that of the host species (Fig. 3A, middle/right; see Fig. S3A for the per-cell normalized count distribution). This suggests that the divergence in the expression of *Efnb3* in forebrain glutamatergic neurons was driven almost entirely by extrinsic factors, which is reflected in its very high proportion extrinsic divergence of 0.92. On the other hand, the expression of *Lsm6* was much higher in mouse cells regardless of their extracellular environment and has an intrinsic proportion divergence of 0.996, suggesting purely intrinsic divergence between species (Fig. 3B, Fig. S3B).

**Fig. 3:**
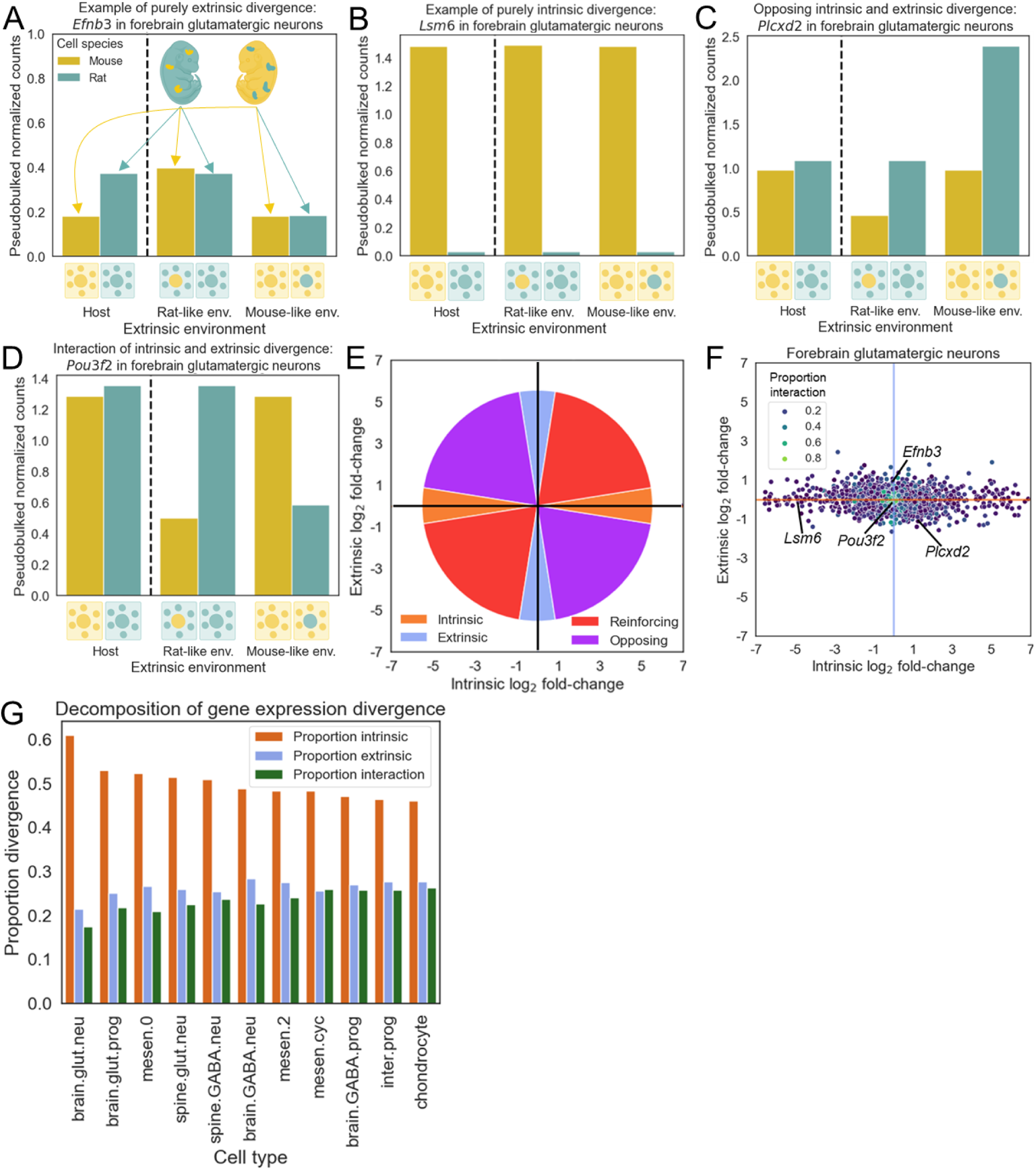
Decomposition of gene expression divergence into cell-intrinsic, cell-extrinsic, and interaction components. **A)** Expression of *Efnb3* in forebrain glutamatergic neurons as an example of purely extrinsic divergence. On the left of the dashed line, the expression of *Efnb3* is shown in species-matched environments. To the right of the dashed line, expression in mouse cells and rat cells is first shown in a rat-like environment and then in a mouse-like environment. The data used for the bars in species-matched environments to the left of the dashed line is from the same host cells as the data in species-matched environments to the right of the dashed line and is included for clarity. **B)** Expression of *Lsm6* in forebrain glutamatergic neurons as an example of intrinsic divergence. **C)** Expression of *Plcxd2* as an example of opposing extrinsic and intrinsic divergence. **D)** Expression of *Pou3f2* in forebrain glutamatergic neurons as an example of intrinsic-extrinsic interaction. **E)** Schematic outlining how each gene’s location in panel F indicates intrinsic and extrinsic contributions to its expression divergence. Purely intrinsic divergence lies along the x-axis, purely extrinsic divergence along the y-axis, and a combination of both (which can be reinforcing or opposing in direction) occurs closer to the four diagonals. **F)** Plot of intrinsic and extrinsic divergence in forebrain glutamatergic neurons. Each point represents a single gene and the example genes shown in panels A-D are indicated on the graph. Points are colored by the proportion of gene expression divergence explained by interaction. **G)** Barplot showing the mean proportion of gene expression divergence explained by intrinsic divergence, extrinsic divergence, and their interaction across all genes that met filtering criteria in each cell type.

However, most genes do not fit neatly into either category. For example, in both mouse and rat cells, *Plcxd2* had higher expression in a mouse-like environment than in a rat-like environment, suggesting extrinsic effects (Fig. 3C, Fig. S3C). However, it also had higher expression in rat cells than in mouse cells in both chimeras, consistent with intrinsically higher expression in rat cells. These two opposing differences cancel out, resulting in nearly identical expression in host cells and approximately equal intrinsic and extrinsic proportions (0.46 and 0.53, Fig. 3C) (see Figs. S4-9 and Supplementary Text 1 for discussion of how developmentally dynamic gene expression and differences in the spatial distribution of donor cells between cell types might affect our results). While most genes primarily exhibit some combination of extrinsic and intrinsic divergence, some genes have evidence for an interaction between the two forms of divergence. For example, *Pou3f2* is down-regulated in species-mismatched environments in both chimeras, which cannot be explained by any simple combination of intrinsic and extrinsic divergence, suggesting an interaction causing *Pou3f2* in mouse cells to respond differently to the extracellular environment than it does in rat cells (Fig. 3D, Fig. S3D). This is reflected in *Pou3f2* having an interaction proportion of 0.85.

### Cell-intrinsic divergence drives gene expression evolution

To summarize these results genome-wide, we visualize these values on a scatter plot so that genes with purely intrinsic divergence are on the x-axis and those with purely extrinsic divergence are on the y-axis (Fig. 3F-G, Figs. S10-11, Supplementary Table 1). Across all cell types, we consistently observed that the genome-wide intrinsic component of gene expression divergence is larger than the extrinsic component, which is in turn larger than the interaction component (Fig. 3G and Figs. S10-11). We estimate that 21-28% of gene expression divergence is cell-extrinsic across cell types, suggesting that a substantial fraction of the *trans*- regulatory divergence between species is due to differences in the extracellular environment.

We found that intrinsic divergence is substantially more correlated across cell types than extrinsic divergence (Fig. S12A), consistent with the idea that *cis*-regulatory variation is also less tissue-specific than *trans*^19–21^. Interestingly, neural and mesenchymal cell types clustered separately in terms of intrinsic divergence, but for extrinsic divergence they instead clustered by location within the embryo, resulting in spinal neurons clustering with other cells from more caudal parts of the embryo such as mesenchyme (Fig. S12B). This suggests that when specifically comparing extrinsically-driven divergence, the similar extracellular environment shared between both neuronal and non-neuronal caudal cell types could be the dominant factor, overriding the tendency for similar cell types such as neurons to cluster together. In addition, we evaluated whether several gene-level variables were correlated with intrinsic vs extrinsic divergence proportions. The best predictor for individual genes was each gene’s expression difference between mouse and rat: all genes with >4-fold difference had primarily intrinsic divergence (Fig. S12C), consistent with the finding that *cis*-regulatory divergence (which is cell-intrinsic) contributes more to large changes in gene expression^22^. Other factors like expression level and tissue-specificity had little to no predictive power (Fig. S12D).

### Cell-extrinsic and intrinsic divergence can propagate through transcriptional networks

To investigate whether extrinsic divergence converges on particular pathways, we tested whether extrinsically-driven genes were enriched for different gene sets, identifying dozens of enrichments (Supplementary Table 2). For example, we found that genes involved in the ER stress response, such as *Hsp90b1*, had >2-fold enrichment for up-regulation in a rat-like environment in 10 out of 11 cell types, Fig. 4A-B, Fig. S13A-B)^23^. Conversely, we found that genes encoding histone methyltransferases, such as *Setbp1*, were highly enriched for up-regulation in a mouse-like environment in neural but not connective tissue (Fig. S14A-D)^24^. Finally, many pathways were strongly enriched for extrinsic divergence in only one cell type, such as positive regulation of programmed cell death which was specifically up-regulated by forebrain GABAergic neurons in a rat-like environment (Fig. S15A-D).

**Fig. 4:**
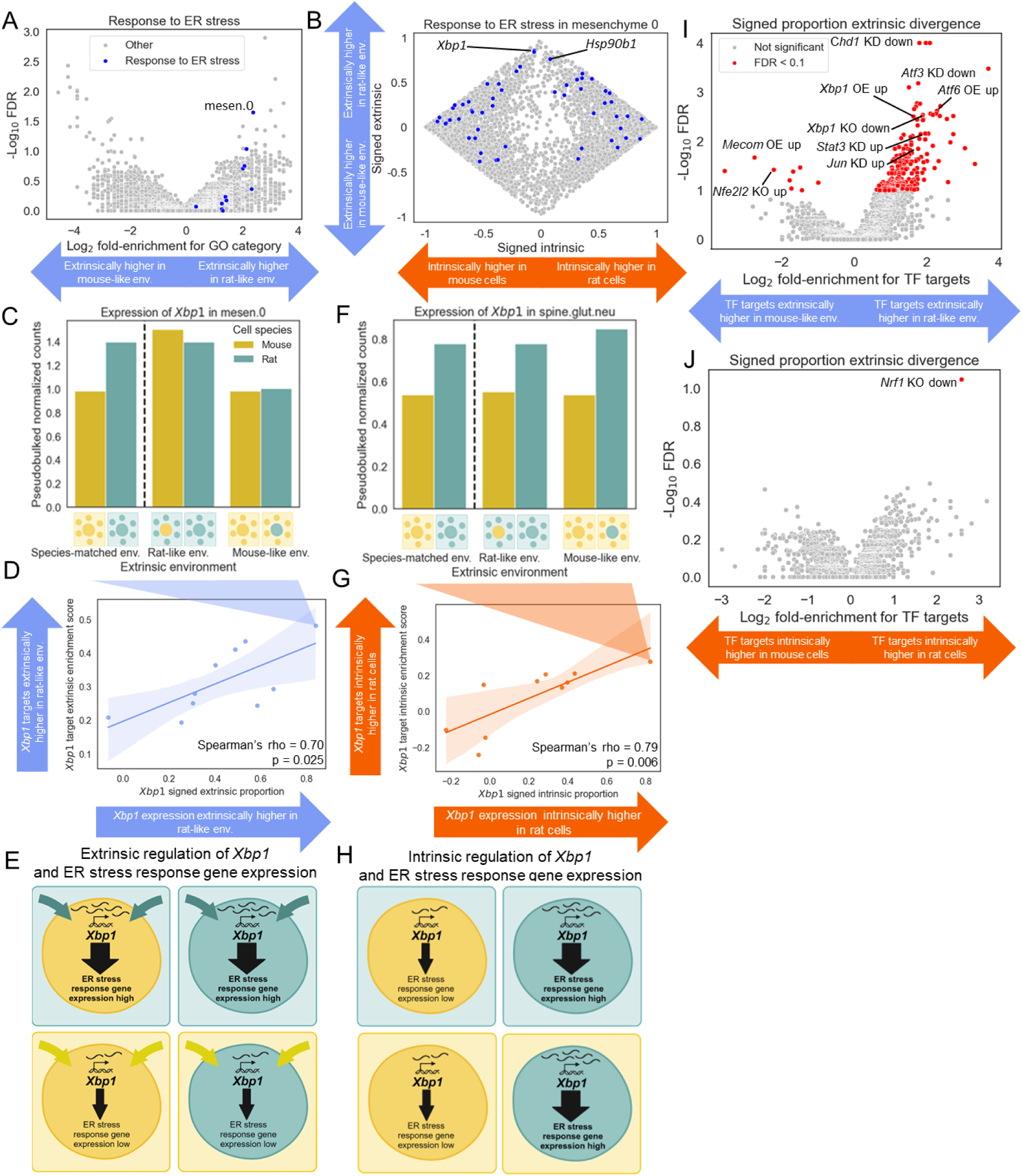
Extrinsic and intrinsic divergence in the expression of *Xbp1* and its target genes. **A)** Enrichment of endoplasmic reticulum stress response genes for positive extrinsic divergence across cell types. Each point is a GO biological process category in a cell type and the points corresponding to the “Response to ER stress” GO category are colored blue. The x-axis shows the log_2_ fold-enrichment and the y-axis shows the −log_10_ false discovery rate. **B)** Scatterplot showing signed proportion intrinsic divergence (x-axis) and signed proportion extrinsic divergence (y-axis) for all genes passing our filtering criteria for mesenchymal cluster 0. ER stress response genes are shown in blue and all other genes are shown in grey. **C)** Expression of *Xbp1* in mesenchymal cluster 0. **D)** Correlation between signed extrinsic proportion divergence for *Xbp1* (x-axis) and the enrichment of its target genes for signed extrinsic proportion divergence (y-axis). Each point is a cell type and cell types with extrinsically driven increased expression of *Xbp1* in a rat-like environment have larger values on the x-axis. Cell types with extrinsically driven increased expression of *Xbp1* target genes in a rat-like environment have larger values on the y-axis. The line and shaded region represent the best fit and 95% confidence interval of a linear model fit to the data. **E)** Conceptual model for *Xbp1*-driven increased expression of ER stress response genes in a rat-like environment occurring in some cell types. Due to extrinsic factor(s), *Xbp1* expression is increased in a rat-like environment compared to a mouse-like environment. This then up-regulates *Xbp1* target genes, many of which are ER stress response genes, in a rat-like environment. **F)** Expression of *Xbp1* in spinal glutamatergic neurons. **G)** Correlation between signed intrinsic proportion divergence for *Xbp1* (x-axis) and the enrichment of its target genes for signed intrinsic proportion divergence (y-axis). Each point is a cell type and cell types with intrinsically driven increased expression of *Xbp1* in rat cells have larger values on the x-axis. Cell types with intrinsically driven increased expression of *Xbp1* target genes in rat cells have larger values on the y-axis. The line and shaded region represent the best fit and 95% confidence interval of a linear model fit to the data. **H)** Conceptual model for *Xbp1*-driven increased expression of ER stress response genes in rat cells occurring in some cell types. Due to intrinsic factor(s), *Xbp1* expression is increased in rat cells compared to mouse cells. This then up-regulates *Xbp1* target genes, many of which are ER stress response genes, in rat cells compared to mouse cells. **I)** Volcano plot of enrichment of TF target genes for signed extrinsic proportion divergence. The y-axis shows the −log_10_(FDR). The x-axis shows the log_2_ fold-enrichment with target sets enriched for higher expression in rat-like environments on the right and mouse-like environments on the left. KO stands for knockout, KD stands for knock down, OE stands for overexpression, up implies increased expression after the experimental manipulation, and down implies decreased expression after the experimental manipulation. **J)** Volcano plot of enrichment of TF target genes for signed intrinsic proportion divergence. The y-axis shows the −log_10_(FDR). The x-axis shows the log_2_ fold-enrichment with target sets enriched for higher expression in rat cells on the right and in mouse cells on the left.

We next hypothesized that intrinsic and extrinsic regulation may propagate through transcriptional regulatory networks; for example, extrinsic regulation of a TF may result in extrinsic regulation of its target genes. In our analysis of ER stress response genes (Fig. 4A) we noticed that divergence in the expression of *Xbp1*, a key transcriptional regulator of ER stress response genes^25^, was almost entirely extrinsic in some cell types such as mesenchyme (Fig. 4D, Fig. S16A). To determine whether extrinsically driven changes in *Xbp1* expression might drive the extrinsic divergence of its target genes, we compared the degree of extrinsic up-regulation in the rat-like environment for *Xbp1* target genes (defined as genes up-regulated by *Xbp1* overexpression)^26^ with that of *Xbp1* itself in each cell type^27^. Remarkably, we observed a strong correlation between these two quantities, where cell types with extrinsic up-regulation of *Xbp1* in the rat-like environment showed a similar pattern of extrinsic up-regulation for its target genes (Spearman’s rho = 0.70, p = 0.025, Fig. 4E). Moreover, we observed a similarly strong correlation between *Xbp1* extrinsic divergence and the extrinsic divergence of ER stress response-associated genes (Spearman’s rho = 0.73, p = 0.014, Fig. S17A). This suggests that extrinsic regulation of *Xbp1* may lead to extrinsic regulation of its target genes (Fig. 4F), though we cannot rule out a potential role for other unobserved factors such as post-transcriptional regulation^25^.

Interestingly, although *Xbp1* had a similar magnitude of up-regulation in both rat host spinal glutamatergic neurons and rat host mesenchyme compared to their mouse host counterparts (Fig. 4D, G), this was almost entirely due to intrinsic divergence in the spinal neurons (Fig. 4G, Fig. S16B). Analogous to the pattern for extrinsic divergence (Fig. 4E), we found that cell types with intrinsic up-regulation of *Xbp1* in rat cells showed a similar pattern of intrinsic up-regulation of its target genes (Spearman’s rho = 0.79, p = 0.006, Fig. 4H), although in this case *Xbp1* intrinsic regulation was not as strongly predictive of intrinsic up-regulation of ER stress response genes more broadly (Spearman’s rho = 0.37, p = 0.29, Fig. S17B). This suggests that intrinsically driven divergence in *Xbp1* expression may propagate to its target genes (Fig. 4I) in much the same way as extrinsic divergence.

More generally, we expanded our analysis to test whether the known targets of many different TFs were enriched for unidirectional extrinsic divergence. Notably, the targets of several well-known regulators of ER stress including *Atf3*, *Atf6*, and *Jun* were also enriched for extrinsic up-regulation in a rat-like environment, similar to *Xbp1* (Fig. 4J)^28–30^. The extrinsic regulation of these TFs was not correlated with that of *Xbp1* (Fig. S18), suggesting divergence in distinct extrinsic factors that converge on TFs regulating the ER stress response. Overall, we observed 155 enriched TFs at a 10% FDR, compared to just one enriched TF when performing an identical procedure with intrinsic regulation (Fig. 4K, Supplementary Table 3).

### Complex patterns of intrinsic and extrinsic divergence underlie coordinated cell type-specific expression of protein complex subunits

The single TF with targets enriched for intrinsic divergence was *Nfe2l1* (which encodes the protein Nrf1), a well-established regulator of proteasomal genes (Fig. 4K)^31^. For example, *Pomp*, a target of Nrf1 involved in the formation and activation of the proteasome, was consistently more highly expressed in rat cells than mouse cells regardless of the extracellular environment (Fig. S19A-F)^31^. In contrast, the targets of other proteasome TFs such as *Nfe2l2* (encoding the protein Nrf2) and *Stat3* were enriched for extrinsic up-regulation in mouse-like and rat-like environments, respectively (Fig. 4J)^32,33^. This heterogeneity in the mode (intrinsic vs extrinsic) and direction (rat vs. mouse up-regulation) of divergence in proteasomal TFs contrasted with the four ER stress response TFs mentioned above, whose targets were all enriched for extrinsic up-regulation in a rat-like environment. Interestingly, the conflicting modes and directions of divergence of these proteasomal TFs did not correlate strongly with the divergence of their target genes (Fig. S20A-C), suggesting that heterogeneity in the regulatory divergence of multiple TFs targeting the same genes might lead to more complex patterns than the relatively simple propagation seen for *Xbp1* and its targets.

To investigate the consequences of having multiple proteasomal TFs with differing modes and directions of expression divergence, we explored how proteasomal subunits were regulated across cell types. For example, in rat cells, the proteasomal subunit *Psmb1* had higher expression in a rat-like environment in forebrain GABAergic neurons, but higher expression in a mouse-like environment in spinal glutamatergic neurons and chondrocytes (Fig. 5A-C, Fig. S21A-C). Interestingly, many proteasomal genes showed a similar pattern (Fig. 5D-F). In forebrain GABAergic neurons, expression was up-regulated both intrinsically in rat cells and extrinsically by the rat-like environment. However, in spinal glutamatergic neurons, the intrinsic effect was preserved but the extrinsic rat-like environment had the opposite effect, down-regulating expression. Lastly, for chondrocytes, both effects were flipped compared to forebrain GABAergic neurons: both the intrinsic and extrinsic effects of rat were repressive. Based on this finding, we looked for other gene sets that showed similarly variable enrichments across cell types and identified context-specific patterns of intrinsic and extrinsic components for spliceosomal genes as well as nuclear-encoded subunits of mitochondrial respiratory complex I (Figs. S22-23). This suggests that highly coordinated context-specificity, integrating both extrinsic and intrinsic signals, has evolved to regulate multiple protein complexes, perhaps to maintain proper stoichiometry while modulating their abundances in response to both extrinsic and intrinsic cues.

**Fig. 5:**
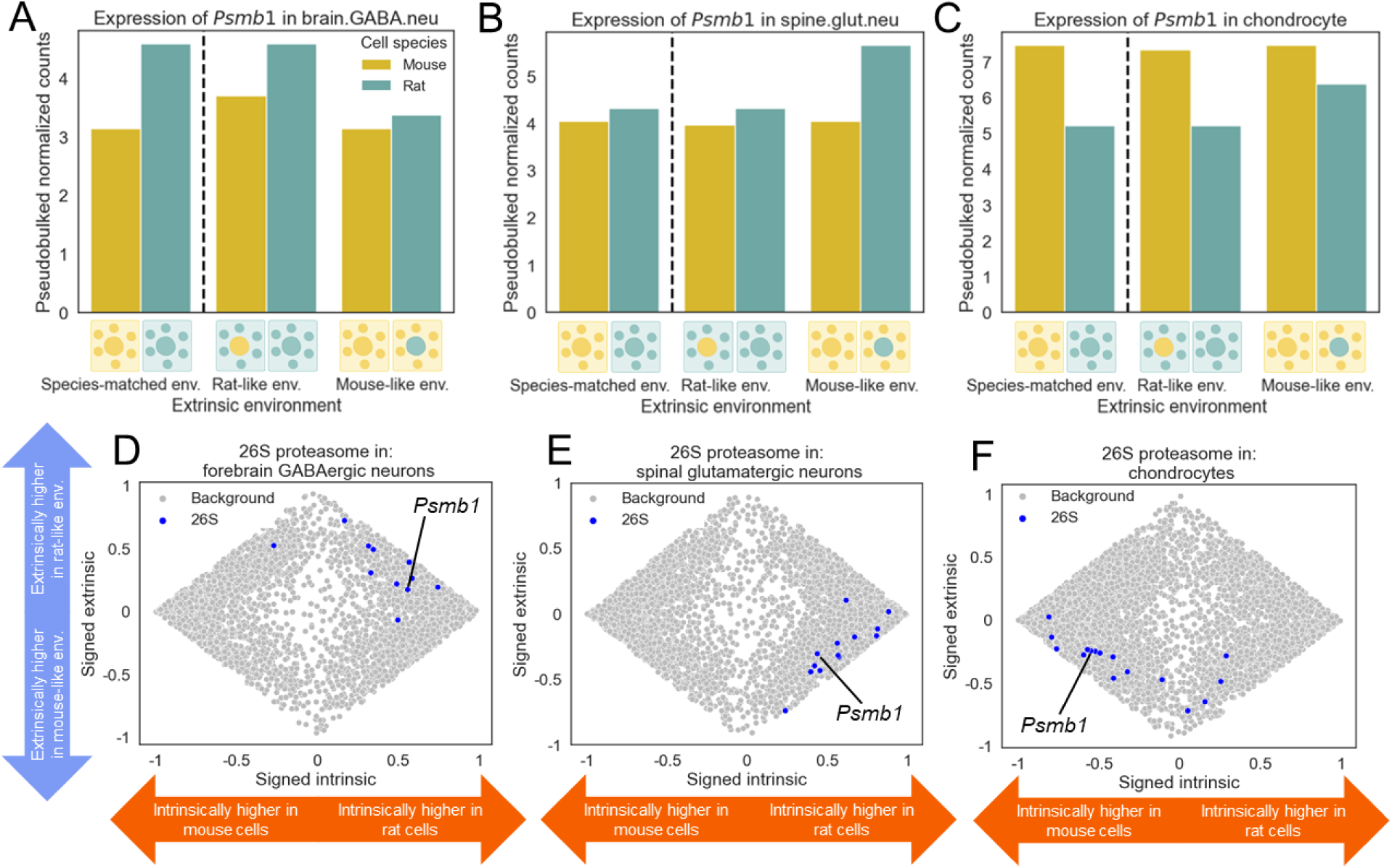
Cell-type specific intrinsic and extrinsic divergence of the expression of genes encoding proteasomal subunits across cell types. **A)** Expression of *Psmb1* in forebrain GABAergic neurons. **B)** Same as in (A) but for spinal glutamatergic neurons. **C)** Same as in (A) but for chondrocytes. **D)** Scatterplot showing signed proportion intrinsic divergence (x-axis) and signed proportion extrinsic divergence (y-axis) for all genes passing our filtering criteria for forebrain GABAergic neurons. Genes coding for 26S proteasomal subunits are shown in blue and all other genes are shown in grey. **E)** Same as in (D) but for spinal glutamatergic neurons. **F)** Same as in (D) but for chondrocytes.

### Cell-extrinsic divergence in the protein expression of two ER stress response genes

To test whether the extrinsically-driven gene expression divergence we observed translates to the protein level, we used immunofluorescence (IF) to quantify extrinsic, intrinsic, and interaction components of divergence in protein levels for two proteins: *Jun* and *Hspa5*^34^. We prioritized these genes because they are involved in the ER stress response, show strong extrinsic divergence across cell types (especially in GABAergic forebrain neurons and progenitors, Fig. 6A-B, Fig. S24A-D), and are sufficiently highly expressed to be detected with IF. In the ganglionic eminences (which contain both GABAergic progenitors and recently born GABAergic neurons) of two mouse-like and two rat-like chimeras, we observed extrinsically-driven up-regulation in rat-like environments for both proteins, closely matching the estimates from our scRNA-seq data (Fig. 6C-E). This suggests that extrinsic divergence in mRNA abundances can translate to the protein level, and also that our estimates of extrinsic divergence for these genes are unlikely to be inflated by experimental variability.

**Fig. 6:**
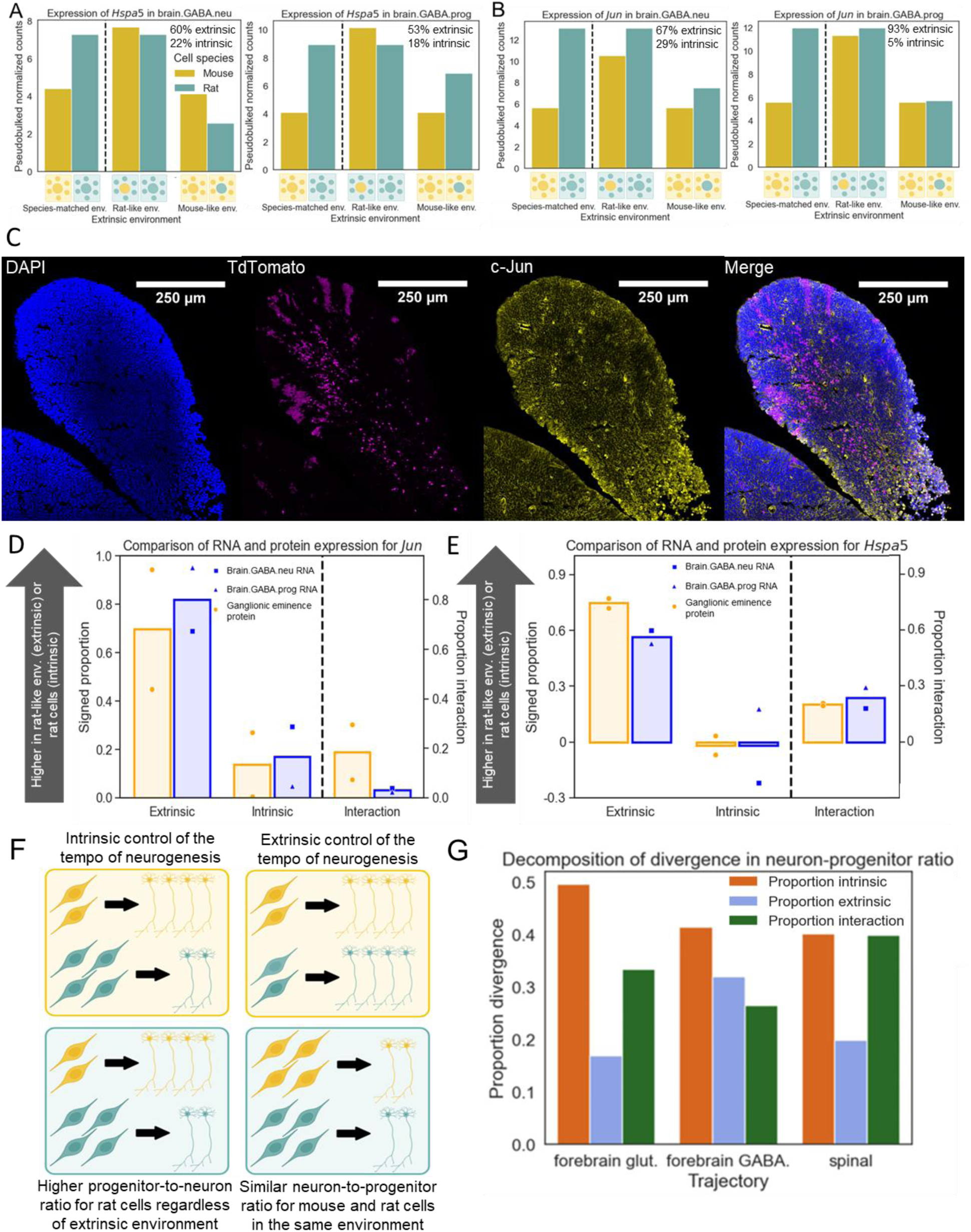
Extrinsic divergence of *Jun* and *Hspa5* at the RNA and protein levels. **A)** Expression of *Hspa5* in forebrain GABAergic neurons (left) and progenitors (right). The percentage of gene expression divergence explained by extrinsic and intrinsic factors is shown in the upper right of each plot. **B)** The same as in (A) but showing the expression of *Jun*, which codes for the protein c-Jun. **C)** Immunofluorescent staining for c-Jun in the medial ganglionic eminence of a rat-like chimera. Donor mouse cells are marked with TdTomato and c-Jun staining is shown in yellow. **D)** Comparison of intrinsic, extrinsic, and interaction divergence at the RNA and protein level for *Jun*. To the left of the dashed line, the signed proportion divergence is along the y-axis, with larger values indicating a greater proportion of either intrinsic or extrinsic divergence leading to higher expression in rat. The extrinsic and intrinsic divergence for protein levels across two chimera pairs is shown in blue and the divergence in RNA expression in orange for GABAergic neurons (square marker) and neurons (triangle marker). The bars indicate the average across replicates for protein and the two cell types for RNA. To the right of the dashed line, the proportion of interaction divergence (which is non-negative by definition) is shown with its own y-axis. **E)** Similar to (D) but showing *Hspa5* divergence. **F)** Conceptual model for extrinsic or intrinsic divergence in the pace of neurogenesis. The left outlines intrinsic divergence as mouse cells include more neurons than progenitors regardless of whether they are in a rat-like or mouse-like environment. The right outlines extrinsic divergence in which the ratio of neurons to progenitors is similar for mouse and rat cells in the same extracellular environment, but differs between environments. **G)** Proportion intrinsic, extrinsic, and interaction divergence for neuron-progenitor ratio in forebrain glutamatergic neurogenesis, forebrain GABAergic neurogenesis, and spinal GABAergic and glutamatergic neurogenesis together.

### Extrinsic and intrinsic components of neural progenitor differentiation kinetics

As discussed above, our framework is applicable to most quantitative traits. To demonstrate this, we explored the application of this framework to estimate extrinsic, intrinsic, and interaction components of a cellular-level quantitative trait: the ratio of post-mitotic neurons to neural progenitors, which reflects the timing of neurogenesis. If the timing of neurogenesis is intrinsically controlled, we would expect a higher neuron to progenitor ratio for mouse cells, regardless of their extracellular environment (Fig. 6F, top). On the other hand, if this trait is extrinsically controlled, we would expect similar ratios for both mouse and rat cells in the same extracellular environment (Fig. 6F, bottom). Strikingly, across three different branches of neurogenesis (forebrain GABAergic, forebrain glutamatergic, and spinal), we consistently found that the ratio of neurons to progenitors was much higher in mouse host cells than in rat donor cells, consistent with an intrinsic component for the faster development of mouse neurons and faster global maturation rate of mouse embryos (Fig. S25A-C). However, we also found this ratio to be similar for mouse donor cells and rat host cells, consistent with previous work showing an effect of the rat maternal environment on the timing of neurogenesis (Fig. S25A-C)^9^. Overall, we estimated an average contribution of 44% intrinsic, 23% extrinsic, and 33% interaction for divergence in the timing of mouse and rat neuronal differentiation (Fig. 6G).

### Imprinted genes are strongly mis-expressed in species-mismatched environments

During our analysis, we noticed that several imprinted genes had strikingly large interaction components. For example, while the imprinted gene *Grb10* was highly expressed in both mouse and rat chondrocytes when in species-matched environments, its expression was nearly zero in both mouse and rat chondrocytes in species-mismatched environments (Fig. 7A, Fig. S26A)^35^. In GABAergic progenitors *Grb10* expression in mouse cells was unaffected by the extracellular environment, but was undetectable in rat cells in a mouse-like environment (Fig. 7B, Fig. S26B). In contrast, *Grb10* expression was unaltered in species-mismatched environments in forebrain GABAergic neurons (Fig. 7C, Fig. S26C). This stark divergence was not restricted to *Grb10* as, for example, *Igf2* is more highly expressed in rat chondrocytes than mouse chondrocytes in species-matched environments, but drops close to zero when rat chondrocytes are in a mouse-like environment (Fig. S27A, D)^36^.

**Fig. 7:**
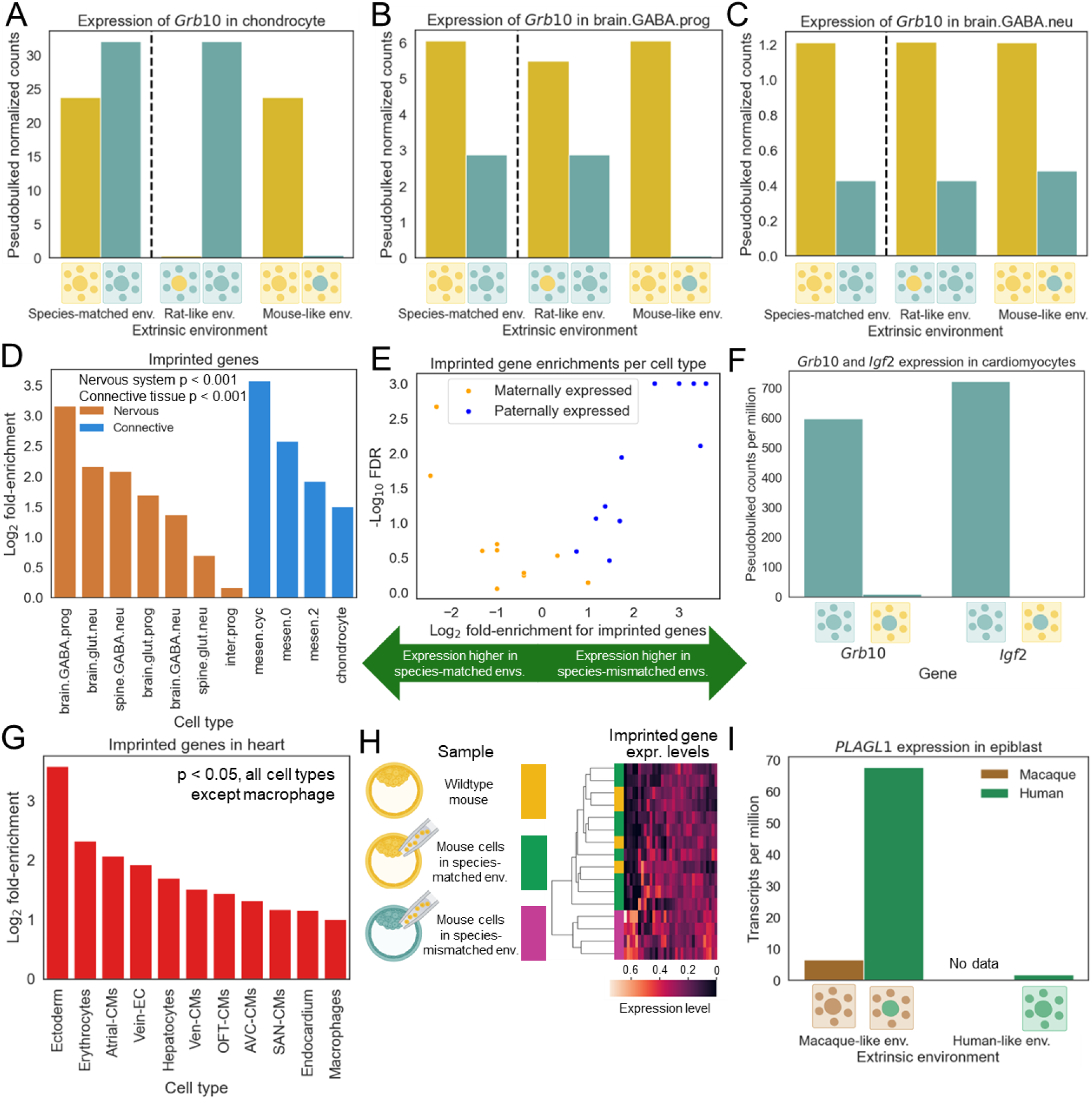
Misexpression of imprinted genes in species-mismatched environments. **A)** Expression of the imprinted gene *Grb10* in chondrocytes. **B)** Expression of *Grb10* in forebrain GABAergic progenitors. **C)** Expression of *Grb10* in forebrain GABAergic neurons. **D)** Enrichment of imprinted genes for high absolute interaction divergence across cell types. The bars represent the log_2_ fold-enrichment. **E)** Enrichment of maternally expressed and paternally expressed genes for signed interaction divergence. Each of the eleven cell types analyzed in this study is represented by one blue point and one orange point. The y-axis shows the −log_10_(FDR) and the x-axis shows the log_2_ fold-enrichment. A negative enrichment score indicates higher expression in species-matched environments, whereas a positive enrichment score indicates higher expression species-mismatched environments. **F)** Expression of imprinted genes *Grb10* and *Igf2* in ventricular cardiomyocytes. **G)** The same as in (D), but for cell types in the heart dataset and using the absolute log fold-change between expression in a mouse-like environment and expression in a rat-like environment to rank genes when computing enrichments. **H)** Heatmap of imprinted gene expression across samples for adult parathyroid data. Samples were hierarchically clustered using the Euclidean distance metric. **I)** Expression of the imprinted gene *PLAGL1* in macaque and human epiblast. No data are shown for macaque cells in a human-like environment since these have not been reported.

To explore whether this pattern holds for imprinted genes more generally, we tested whether previously reported imprinted genes were enriched for interactions between extrinsic and intrinsic divergence. Remarkably, we observed that imprinted genes were strongly enriched for large interaction components (p < 0.001 in both connective and nervous tissue, Fig. 7D). Given that genes expressed from the maternal allele (maternally-expressed) and expressed from the paternal allele (paternally-expressed) often have opposing effects on growth, we split imprinted genes based on the expressed allele^37^. We found that paternally-expressed genes were strongly enriched for higher expression in species-mismatched environments (p < 0.01 in both brain and connective tissue), whereas maternally-expressed genes were enriched for higher expression in species-matched environments only in connective tissue (p < 0.01 in connective tissue, p > 0.5 in brain tissue), suggesting that maternally-expressed and paternally-expressed genes are often affected by species-mismatched environments in opposite ways (Fig. 7E). The pattern for paternally expressed genes is exemplified by *Zdbf2*, which is expressed approximately 2-fold higher in species-mismatched environments for both mouse cells and rat cells across connective and nervous cell types (Fig. S27B-C, E-F)^38^.

To determine if this pattern of disrupted imprinting generalized to other types of cells and chimeras, we sought to test for this effect in other chimera scRNA-seq data sets. Because these studies do not have reciprocal chimeras, we could not disentangle the extrinsic and interaction components, but we could test the simpler prediction that imprinted genes should be highly sensitive to the extracellular environment. In one previous study, rat-mouse chimeras were generated in which the hearts were almost entirely derived from rat donor cells, and their gene expression was compared to wildtype rat heart cells from the same embryonic stage^39^. Despite the data being independently generated using different host and donor strains, tissues, and developmental stages, we observe that that *Grb10* and *Igf2* were highly expressed in wildtype rat but were nearly undetectable in rat cells present in a mouse-like environment across all 11 heart data cell types tested (Fig. 7F, Fig. S28A-B). More generally, imprinted genes were strongly enriched for differential expression between rat cells in a rat-like environment vs. rat cells in a mouse-like environment (p < 0.05 in all cell types except macrophages where p = 0.11, Fig. 7G).

One possible explanation for disrupted imprinting in donor cells is that imprinting could already be disrupted in the ESCs from which these cells are derived. To test this, we examined a published bulk RNA-seq dataset of adult parathyroid gland cells where the same mouse ESCs were injected into both rat and mouse blastocysts and compared to wildtype mice with no blastocyst injection^40^. If imprinting disruption is due to the species-mismatched environment rather than any technical factor related to cell lines or chimera generation, then we would expect imprinted genes to be mis-expressed in rat-mouse chimeras but not mouse-mouse chimeras.

To evaluate this, we hierarchically clustered all samples based on the expression of all expressed imprinted genes. We found that samples from species-mismatched environments clearly clustered together, whereas samples from wildtype mice and donor mouse cells in a species-matched environment were intermixed in a separate cluster (Fig. 7G). For example, *Grb10* and *Igf2* were strongly mis-expressed in a species-mismatched environment (Fig. S29A). Overall, these results suggest that a species-mismatched environment, not blastocyst injection of ESCs more generally, underlies mis-expression of imprinted genes and that mis-expression can persist into adulthood.

Finally, we investigated whether imprinting may also be disrupted in *ex vivo* human-macaque chimeric embryos^41,42^. We found that many imprinted genes, such as *PLAGL1,* showed strong mis-regulation in human cells growing in a macaque-like embryo (Fig. 7I). Imprinted genes were enriched for differential expression between human cells vs. macaque cells in the same macaque-like environment, as well as between human cells in a human-like environment vs. in a macaque-like environment (p < 0.01 in each). These results suggest that imprinting disruption in chimeras does not depend on the maternal environment (which is not present for *ex vivo* chimeras) and occurs in both rodents and primates. Overall, our results suggest that the interaction between extrinsic and intrinsic divergence results in widespread mis-expression of imprinted genes in interspecies chimeras.

## Discussion

Despite widespread interest in the concept of developmental autonomy for over a century^1,2^, and numerous fundamental discoveries^3–9^, its investigation has remained a qualitative and low-throughput endeavor. Building on this foundational body of work, we have developed a broadly applicable framework to decompose divergence in any quantitative trait into cell-extrinsic and intrinsic components (Fig. 2). This enables not only mathematical precision, but also the potential for high-throughput measurement of intrinsic and extrinsic contributions to thousands of traits in a single experiment.

Application of this framework to gene expression divergence between mouse and rat revealed a preponderance of intrinsic factors in the evolution of gene expression (Fig. 3). However, extrinsic factors also contribute to many genes, and even dominate the divergence of some critical pathways such as the ER stress response (Fig. 4). In other cases, such as the proteasome (Fig. 5), histone methyltransferases (Fig. S14), and regulators of programmed cell death (Fig. S15), the contributions of extrinsic factors are highly cell type-specific. This cell type-specificity enabled us to track the propagation of extrinsically dominated divergence through transcriptional networks (Fig. 4) as well as to the protein level (Fig. 6).

In addition, we identified widespread mis-regulation of imprinted genes as a result of interactions between extrinsic and intrinsic divergence (Fig. 7), suggesting that imprinting may be controlled by a complex interplay between rapidly evolving intrinsic and extrinsic factors. As many imprinted genes play key roles in growth and cell survival, our results also suggest that modulating the expression of imprinted genes could be a promising avenue to improve the growth and survival of cells in species-mismatched environments^46^. Although it is outside the scope of this work, it will be interesting to explore the effects of xenotransplantation on the expression of imprinted genes as well.

Our framework can also provide insight into the genetic basis of interspecies chimeric incompatibility^43^. For example, we identified strong extrinsic divergence in ER stress response genes such as *Hspa5,* which is vital for neuronal survival during development^44^, as well as extrinsic divergence in histone methyltransferases, which play a key role in controlling the tempo of neuronal development^45^. Consistent with this, we found that the tempo of neurogenesis—which is faster in mouse than in rat—has evolved via a complex combination of both intrinsic and extrinsically-driven divergence, as well as interactions between these.

Interestingly, although the intrinsic:extrinsic ratio was similar for neuronal differentiation (Fig. 6G) and global gene expression (Fig. 3G)—about 2:1—neuronal differentiation showed a substantially greater contribution of intrinsic-extrinsic interaction. Further work will be required to confirm this finding across different species and developmental stages, and to disentangle effects such as maternal environment^9^ from the host embryo’s genotype. Nevertheless, it is tempting to speculate that this difference may be related to the greater complexity of cellular-level traits compared to individual gene expression levels, and that increasingly complex traits may show a general trend towards greater intrinsic-extrinsic interaction.

A recent study reported rat-mouse chimeras where rat ESCs were injected into host mice blastocysts that were unable to generate a forebrain (*Hesx1*^-/-^)^18^. This study is challenging to directly compare to ours since donor cells composed ∼60% of the forebrain at the time of scRNA-seq profiling, suggesting that the extracellular environment was neither mouse-like nor rat-like. In addition, the study did not have reciprocal chimeras or a quantitative analysis framework. Perhaps in part because of these and other technical factors, our results differ substantially from theirs. Specifically, they found the pace of neurodevelopment (at a gross anatomical level) in rat donor cells to be extrinsically controlled, while the overall transcriptome was intrinsically determined. In contrast, using reciprocal chimeras and our quantitative framework we found both neurodevelopment (at a single-cell level) and the transcriptome to be subject to about twice as much intrinsic as extrinsic divergence, along with a substantial effect of interaction between these, especially for neurodevelopment.

Our study opens the door to myriad avenues for further research. While our initial study focused on gene expression, the same framework can be applied to any quantitative trait. For example, we can now quantify the cell-intrinsic vs extrinsic contributions to the evolution of numerous components of gene regulation such as chromatin accessibility, DNA methylation, histone post-translational modifications, 3D chromosomal contacts, splicing, translation, etc. In addition, questions about the cell-autonomy of cellular morphology, electrophysiology, differentiation dynamics, cell fate decisions, somatic mutations, cell death, and protein subcellular localization can begin to be quantitatively addressed.

Other exciting future directions include sampling chimeras across a time course of development, which will allow us to disentangle the role of developmental time from intrinsic and extrinsic divergence to better understand the evolution of developmentally dynamic gene expression. In addition, our framework could be applied in conjunction with spatial transcriptomic measurements to further decompose extrinsic divergence into local and organ-wide components. Finally, our approach can be applied to any pair of strains or species compatible with chimerism. Overall, the application of our framework to a wide range of quantitative traits will enable new insights into the mechanistic basis of developmental processes and phenotypic evolution across the tree of multicellular life.

## Methods

### Animals

Seven-week-old Wistar female rats and 10-week-old male rats were purchased from Charles River Laboratory (Wilmington, MA; CRL: 003). Seven-week-old CD1 female mice and 10-week-old male mice (CRL: 022) were purchased from Charles River Laboratory. Seven-week-old C57BL/6 female mice and 10-week-old male mice were purchased from Jackson Laboratories (Bar Harbor, ME: 000664). Littermates of the same sex were randomly assigned to experimental groups. All rats and mice were housed in pathogen-free conditions with free access to food and water. All animal protocols were approved by the Administrative Panel on Laboratory Animal Care at Stanford University.

### ESC culture

Undifferentiated tdTomato-labelled or GFP-labeled mouse ESCs (SUN106.2 and SGE2) were maintained on mitomycin-c treated mouse embryonic fibroblasts (MEFs) in N2B27 medium^47^ containing 1000 U/ml LIF (Peprotech, Cranbury, NJ; 300-05), 1 µM MEK inhibitor PD0325901 (Tocris, Barton Ln, Abingdon, United Kingdom; 4192) and 3 μM GSK3 inhibitor CHIR99021 (Tocris: 4423). Undifferentiated tdTomato-labelled rat ESCs were maintained in N2B27 medium containing 1 μM MEK inhibitor PD0325901, 3 μM CHIR99021, and 1000 U/ml of rat LIF as described^48^. Their pluripotency was confirmed by chimera generation assay. All PSC lines used in this study were male lines.

### Embryo culture and manipulation

Wild-type mouse embryos were prepared according to published protocols^49^. In brief, morula-stage embryos were obtained by uterus perfusion from superovulated CD1 mice at 2.5 days postcoitum (dpc). Morula-stage embryos were cultured in KSOM-AA medium (CytoSpring, Mountain View, CA; K0101) for 1 day and developed to blastocyst-stage embryos. Wild-type rat blastocysts were obtained by uterus perfusion from female rats at 4.5 dpc and cultured in Rat KSOM medium. For micromanipulation, mouse or rat ESCs were trypsinized and suspended in mouse or rat ESC culture medium. A piezo-driven micromanipulator (Prime Tech, Tsuchiura, Japan) was used to pierce the zona pellucida and trophectoderm under microscopy and 5–7 mouse or rat ESCs were introduced into blastocyst cavities near the inner cell mass. After blastocyst injection, embryos were cultured for 1–2 hours. Mouse blastocysts were then transferred into uteri of pseudopregnant recipient CD1 female mice at 2.5 dpc. Rat blastocysts were then transferred into uteri of pseudopregnant recipient Wistar female rats at 3.5 dpc. Table S1 shows results of the cell injection.

### Chimera dissection and cell preparation

Embryonic day (E) 15.25 mouse-rat and E13.5 rat-mouse chimeras were dissected to harvest forebrain and connective tissue. In brief, chimeric fetuses were collected from either E15.25 rat or E13.5 mouse pregnant mothers and forebrain and connective tissues were harvested from those chimeric fetuses. Both forebrains and connective tissues were dissociated in the solution containing 2mg/ml Collagenase/Dispase (Roche, Basel, Switzerland; 10269638001) in Hanks’ Balanced Salt solution. After 30-60 min incubation at 37 degrees, 10% fetal bovine serum (FBS) in PBS solution was added into the tissue solution to inactivate the enzymes. All the dissociated cells were then filtered and used for the subsequent experiment.

### Flow cytometry

Dissociated cells derived from forebrain and connective tissue were stained with APC-anti-mouse CD45 antibody (Biolegend, San Diego, CA; 103112), PE-Cy7-anti rat CD45 (Biolegend: 202214). Donor chimerism was analyzed by detecting CD45 negative, tdTomato or GFP-expressing cells (detailed in Table S2). Both donor and host cells in the forebrain and connective tissue of chimeric fetuses were sorted using a FACS Aria II (BD, Franklin Lakes, NJ).

### Single cell RNA-sequencing

The chimeric fetuses from the CD1-Wistar chimeras that presented less than 10% donor contribution to either forebrain or connective tissue were used for single cell RNA-sequencing. The connective tissue and forebrain of three rat-like chimeras, three mouse-like chimeras, one wildtype mouse, and one wildtype rat were used for scRNA-seq using 10x 3’ v3.1 chemistry. The connective tissue and forebrain were from different chimeras but pooled together before the cells entered the 10x controller. Of the resulting eight libraries, the rat-like chimera in library MR1 had high quality data for forebrain and connective tissue, the mouse-like chimera in library RM1 had sufficiently high quality data for the neuronal lineage, the mouse-like chimera in library RM2 had sufficiently high quality data for connective tissue, and both the wildtype mouse and rat libraries had sufficiently high quality data for both tissue types.

### Histological analysis

Tissues were fixed with 4% paraformaldehyde and embedded in Optimal Cutting Temperature (O.C.T.) compound (Sakura Finetek, Tokyo, Japan). OCT-embedded sections were stained with antibodies for fluorescence microscopic analysis. Each section was incubated with the primary antibodies (Abs) for 24 hours at 4 degrees and with the secondary Abs for 1 hour at room temperature (detailed in Table S3). Following a wash step, sections were mounted with Vibrance Antifade Mounting Medium with DAPI (Vector LABORATORIES, Newark, CA; H-1800), and observed under confocal laser scanning microscopy (Carl Zeiss, Oberkochen, Baden-Württemberg, Germany).

### Alignment, species deconvolution, and clustering

All fastq files were aligned to the mouse (GRCm39, from ENSEMBL v109), rat (mRatBN7.2, from ENSEMBL v109), and a concatenated mouse and rat genome using CellRanger v7.1.0^50^. For mapping, we retained protein-coding genes, lncRNA, antisense transcripts, snoRNA, snRNA, miRNA, and scaRNA. The resulting count matrices were read into scanpy v1.9.5 and the proportion of aligned reads to each genome in the concatenated genome was used to determine the species of origin of each cell^51^. A cell was assigned to mouse if greater than or equal to 70% of aligned reads aligned to the mouse genome, assigned to rat if greater than or equal to 70% of aligned reads aligned to the rat genome, and discarded as a doublet otherwise. For all subsequent steps, we only used the mouse-aligned-only counts for mouse cells and the rat-aligned-only counts for rat cells. We then removed all genes on sex chromosomes, restricted to mouse-rat one-to-one orthologs (defined using data from ENSEMBL), and performed standard pre-processing steps to remove low quality cells^50^. For the latter, we performed standard filtering and removed cells with greater than 15% of counts coming from mitochondrial reads and cells with n_genes_by_counts (a commonly used quality control metric) greater than 7500. Next, we converted to the log normalized counts, identified highly variable genes, computed principal components, used the Python implementation of harmony v0.0.9 to integrate the mouse and rat cells, and then found nearest neighbors^52^. We then clustered the cells using the Leiden algorithm v0.10.2 with resolution equal to 0.1^53^. This first round of clustering split cells into neuronal, connective tissue, and hematopoietic lineages. The neuronal and connective tissue lineage cells were then retained for further analysis (see below).

Several libraries did not contain a sufficient number of cells to be useful for our analysis. Therefore, we used the rat-like chimera data from library MR1 for both neural and connective tissue analyses, the mouse-like chimera data from library RM1 for neural analyses, and the mouse-like chimera data from library RM2 for all subsequent analyses after the pseudobulking step (see below).

Our main goal in clustering the cells was to identify cell types and remove cell subtypes that had fewer than 10 cells in any of donor mouse cells, donor rat cells, host mouse cells, or host rat cells in MR1 and RM1 (for neuronal lineages) or MR1 and RM2 (for connective tissue). We repeated the harmony and Leiden-based procedure (with resolution equal to 0.5) on the neural lineages and identified 18 subclusters. Due to a low number of cells in other cell types, we restricted further analysis to forebrain glutamatergic progenitors (*Gli3* high, *Eomes*-, *Mki67*+), forebrain GABAergic progenitors (*Gli3* low, *Eomes*-, *Mki67*+), intermediate progenitors (*Eomes*+, *Mki67+*), forebrain glutamatergic neurons (*Slc17a6*+, *Tcf7l2*-, *Reln*-, *Hoxb8*-, *Mki67*-), forebrain GABAergic neurons (*Gad1*+, *Hoxb8*-), spinal glutamatergic neurons (*Slc17a6*+, *Hoxb8*+), and spinal GABAergic neurons (*Gad1*+, *Hoxb8*+)^54–56^. As some forebrain GABAergic subtypes had very few rat donor cells, we subclustered (Leiden resolution equal to 0.35) that cell type further, removed the subtypes with very few rat donor cells, and then merged the remaining 3 cell subtypes which had similar cell type proportions across the four species-environment combinations.

For the connective tissue, we reclustered the cells (Leiden resolution equal to 0.15) and identified six clusters. We focused subsequent analysis on mesenchymal cells (*Postn* high, *Prrx1* high) and chondrocytes (*Acan*+)^57^. We subclustered both cell types (Leiden resolution equal to 0.25 for mesenchymal cells and chondrocytes) and removed subtypes according to the above criteria (i.e. at least 10 cells in each of the 4 species-environment combinations). After this procedure, we retained one chondrocyte subcluster, two mesenchymal subclusters, and cycling mesenchymal cells (*Mki67*+) for further analysis.

### Gene filtering, pseudobulking, and count normalization

To avoid overly noisy estimates of intrinsic and extrinsic divergence, we removed genes that were not expressed in greater than or equal to 20% of cells in at least one of the four species-environment combinations. For each gene, we summed all counts across cells within a cell type in each library for mouse and rat cells separately and removed genes that had fewer than ten counts in all four species-environment combinations.

To normalize counts, we divided by total expression and multiplied by 10,000 for each pseudobulked sample in each cell type separately. Rather than normalize the counts directly, we first randomly sampled from the gene-count distribution within each sample in each cell type so that the total number of counts was identical across all species-environment combinations. We performed this procedure 100 times and took the mean normalized counts across all 100 samplings. This mean value was highly correlated with the actual normalized counts (Spearman’s rho > 0.999, p < 10^-300^). This procedure was used instead of simply normalizing by total counts because this can lead to biased estimation of log fold-changes when there are few counts in one of the four species-environment combinations. For example, consider a scenario in which donor mouse, host mouse, and host rat all have 1,000,000 total counts whereas donor rat only has 100,000 counts. If a gene has 99 (100 after adding a pseudocount) counts in donor mouse and host mouse and 0 (1 after adding a pseudocount) counts for donor rat and host rat and we normalize to total counts and multiply by 10,000, we will have 1 normalized count in donor mouse and host mouse, 0.01 normalized counts in host rat, and 0.1 normalized counts in donor rat. If we then compute fold-changes, donor mouse divided by host rat would be 100 but donor mouse divided by donor rat would be 10. This difference would propagate and (see below) cause overestimation of the extrinsic or interaction component of gene expression and underestimation of the intrinsic component.

From here, we will refer to normalized counts for mouse cells in a rat-like environment as DM (donor mouse), counts for rat cells in a rat-like environment as HR (host rat), counts for mouse cells in a mouse-like environment as HM (host mouse), and counts for rat cells in a mouse-like environment as DR (donor rat).

### A framework to decompose gene expression divergence into extrinsic, intrinsic, and interaction components

Throughout, we use the log_2_ fold-change to measure gene expression divergence. We formulate the total expression of a single gene in one of the species-environment combinations as the product of an extrinsic component and an intrinsic component. For example, the expression of a gene in a donor rat cell in a mouse-like environment (*D_R_*) is equal to the intrinsic rat expression (*I_R_*) multiplied by the expression induced by the mouse-like environment (*E_M_*). Throughout, the subscript numbers reflect noise and intrinsic-extrinsic interactions make the values derived from different chimeras and cells unequal. In this framework, the normalized counts are:

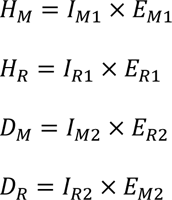

Where *H_M_* is the gene expression for a host mouse cell in a mouse-like environment, *H_R_* is the gene expression for a host rat cell in a rat-like environment, *D_M_* is the gene expression for a donor mouse cell in a rat-like environment, and *D_R_* is the gene expression for a donor rat cell in a mouse-like environment. Using these four values with three degrees of freedom, we can obtain two independent estimates of the intrinsic divergence between mouse and rat (*I*_1_ and *I*_2_):

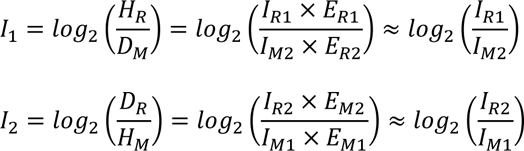

Similarly, we obtain two independent estimates of the extrinsic divergence between mouse and rat (*E*_1_ and *E*_2_):

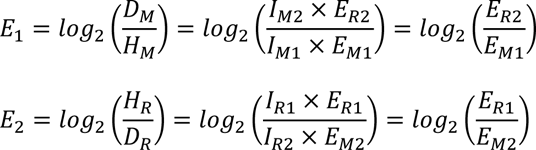

However, for some genes the effects of a rat-like environment on mouse cells will likely differ from the effects of a rat-like environment on rat cells (and vice versa) due to interactions between extrinsic and intrinsic divergence. In the absence of such an interaction, the two estimates of intrinsic divergence should be equal and the two estimates of extrinsic divergence should be equal, so the difference between these pairs of estimates is itself an estimate of the interaction between extrinsic and intrinsic divergence. Therefore, to quantify this interaction, we can compute:

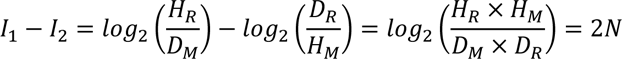

This value is identical to what we obtain starting with extrinsic divergence:

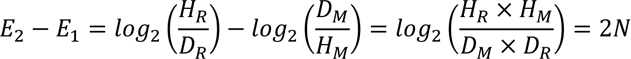

Based on this framework, we obtain our final estimates of intrinsic, extrinsic, and interaction divergence by averaging the two estimates:

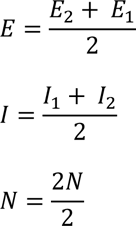

To compute the proportion extrinsic, intrinsic, and interaction divergence, we take the magnitude of each quantity and divide it by the sum of the magnitudes of all three quantities:

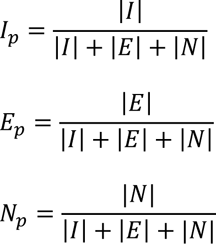

Finally, the signed proportion extrinsic, intrinsic, and interaction divergence is then computed as the sign of the divergence multiplied by the proportion divergence for each quantity. We only applied this framework to genes that had some evidence for divergence in expression level (i.e. an absolute log_2_ fold-change greater than 0.5 in at least one of the four comparisons).

### Analysis of bulk RNA-seq data and developmentally dynamic gene expression

To assess how developmentally dynamic gene expression might influence our results, we focused on the E11.5, E13.5, and E15.5 (or their closest approximation e.g. E15 for rats) timepoints collected from brain tissue in Cardoso-Moreira et al^58^. For mice and rats separately, we computed the Pearson correlation between expression level and developmental stage^59^. We then classified genes as having similar, opposite, or a shift in developmentally dynamic gene expression that goes against the global difference in maturation rate between mouse and rat (referred to as “temporal shift”) according to the following criteria. We considered a gene as having a similar change over development if the product of the correlation coefficients for mice and rats was greater than 0.7 (increasing if both correlation coefficients were positive, decreasing if both correlation coefficients were negative). Similarly, we classified genes as opposite if the product of the correlation coefficients was less than −0.25. These different cutoffs were chosen such that there were a sufficient number of genes for the analysis. As there were many fewer genes with opposite expression trajectories over development, a less strict cutoff (−0.25) was used. To classify genes as having a temporal shift, we restricted to the set of genes with similar changes in expression over time and required that the absolute log fold-change between mouse and rat at both E13.5 and E15.5 be less than 0.25, consistent with the logic outlined in the supplementary text.

To further subdivide genes, we considered a gene as having higher expression in mouse if the log_2_ fold-change of mouse expression divided by rat expression was greater than 0.5 and considered a gene as having higher expression in rat if the log_2_ fold-change was less than −0.5. Restricting to only the genes with similar gene expression trajectories in mouse and rat (i.e. the product of the mouse and rat correlation coefficients was greater than 0.7), we classified genes as increasing in expression over time if a gene had a positive correlation coefficient in mice and rats and as decreasing over time if a gene had a negative correlation coefficient in both mice and rats (for both decreasing and increasing genes, the product would be positive.

In general, to investigate the role of developmentally dynamic gene expression in our results (see Supplementary Text 1 and Supplementary Figs. 4-9), we defined some gene set that we would expect to be enriched for a target divergence (opposing extrinsic and intrinsic divergence, reinforcing extrinsic and intrinsic divergence, mostly extrinsic divergence, or mostly interaction divergence) under a model of purely intrinsic divergence combined with some form of developmentally dynamic gene expression. We then used GSEAPY v1.0.6 preranked to test for enrichment of that gene set at the top or bottom of the gene list when ranking by the target (signed) divergence^27^. Throughout the next section, we describe each gene set in terms of what we would expect under the purely intrinsic model outlined above. We also always use GSEAPY preranked with the following parameters: threads=4, permutation_num=1000, format=’png’, seed=6, min_size = 10, max_size = 30000^27^.

To define the first gene set in which we expect enrichment of both opposing and reinforcing genes, we took the union of genes with similar expression trajectories in mice and rats and genes that were classified as either higher in mouse or higher in rat. We then ranked genes by the product of proportion intrinsic and proportion extrinsic and used GSEAPY preranked to test for enrichment^27^. To further investigate genes with opposing or reinforcing extrinsic and intrinsic divergence, we ranked genes by the product of the signed extrinsic divergence and signed intrinsic divergence so that genes with a large positive value for this metric are reinforcing and genes with a large negative value are opposing. We then defined the gene set in which we expect enrichment of opposing genes as the union of genes that either (1) were increasing over time and had higher expression in mouse or (2) were decreasing over time and had higher expression in rat. Similarly, for the gene set in which we expect enrichment of reinforcing genes we took the union of genes that either (1) were decreasing over time and had higher expression in mouse or (2) were increasing over time and had higher expression in rat. We then ranked genes by the product of the signed intrinsic proportion and signed extrinsic proportion (so that genes with a very low value are opposing and genes with a very high value are reinforcing) and used GSEAPY preranked^27^.

To investigate purely extrinsic genes, we defined the gene set in which we expect enrichment of extrinsic genes as the temporal shift genes (defined above), sorted by extrinsic proportion, and used GSEAPY preranked^27^. Next, we split temporal shift genes into decreasing (expected to be enriched for negative extrinsic divergence) and increasing (expected to be enriched for positive extrinsic divergence) and input both gene sets, sorted by signed extrinsic proportion, and used GSEAPY preranked^27^.

Finally, to investigate interaction divergence genes we defined the gene set we expect to be enriched for interaction divergence as the opposing genes (defined above), sorted by proportion interaction, and used GSEAPY preranked^27^. Next, we defined genes expected to have higher expression in species-matched environments as genes with increasing expression in rat and decreasing expression in mouse and genes expected to have higher expression in species-mismatched environments as genes with increasing expression in mouse and decreasing expression in rat. We then sorted the input gene list by the signed interaction proportion and tested for enrichment using GSEAPY preranked^27^.

### Data processing and analysis for predictors of intrinsic and extrinsic divergence

We tested six variables for their correlation with [intrinsic proportion - extrinsic proportion]. In general, if high quality data were not available for both rats and mice, we used data from the human genome to avoid biasing our analysis. To measure gene expression divergence and total expression in mice and rats, we used the log_2_ fold-change between and average expression of mouse cells in a mouse environment and rat cells in a rat environment. To measure evolutionary constraint, we used the metric for constraint on each gene published in Zeng et al^60^. As a proxy for regulatory complexity, we assigned candidate *cis*-regulatory elements (CREs) from the ENCODE project to the nearest TSS of a protein-coding gene, removed CREs greater than 100,000 bases from the nearest TSS of a protein coding gene, and computed the total number of regulatory elements assigned to each gene^61^. For cell type specificity, we used total expression (as described above) for each of the eleven cell types in our analysis as input to compute Tau, a well-established measure of tissue/cell type-specificity. Tau is equal to one if a gene is completely specific to a single cell type or tissue, and zero if it is equally expressed across all cell types or tissues^62^. To compute tissue specificity in mice and rats at E13.5 and E15.5 respectively, we averaged normalized counts across replicates for six of the seven tissues from Cardoso-Moreira et al. (we excluded cerebellum/hindbrain due to its transcriptional similarity to the rest of the brain) and computed Tau^58^. Finally, we computed the log_2_ fold-change between mouse and rat within each tissue (again using E13.5 and E15.5 respectively and excluding cerebellum/hindbrain) and used the variance of the log_2_ fold-change across tissues as a measure of the tissue-specificity of gene expression divergence.

To merge these six variables into one matrix we converted mouse gene names to human gene names using the orthologs from the Ensembl database and joined all the above metrics together, removing any genes for which one or more of these metrics could not be computed^50^. We then calculated the Spearman correlation between [intrinsic proportion - extrinsic proportion] and each of these variables^59^.

### Enrichment analysis

For the enrichment analysis, we either performed analysis on each cell type separately or averaged expression in connective tissue, the neural lineage (i.e. including spinal neurons with the forebrain cells), or across all eleven cell types. In the first two cases, we required that a gene have passed our filtering criteria (see above) in at least three cell types and in the latter case we required a gene to have passed the filtering criteria in at least five cell types.

Throughout, we always use GSEAPY preranked with parameters: threads=4, permutation_num=1000, format=’png’, seed=6, min_size = 10, max_size = 300^27^. To identify enriched gene sets in extrinsically driven genes, we sorted the input gene list by the signed extrinsic proportion, converted mouse gene symbols to human gene symbols using orthologs defined by Ensembl, and used GSEAPY preranked^27,50^. We tested gene sets from the gene ontology (GO) biological process (2023 version), GO cellular component (2023 version), GO molecular function (2023 version), MGI Level 4 Mammalian Phenotypes (2021 version), and the CORUM protein complex database^63–65^. In all cases, we used the versions directly supported by GSEAPY. After running on all cell types, we retained all terms across the five ontologies with FDR < 0.25.

To identify shared and cell type-specific enrichments, we counted the number of times each term occurred in this final merged file. To further investigate shared terms, we visualized the signed extrinsic and signed intrinsic proportion as shown in Fig. 4G-H. The log_2_ fold-enrichment shown in many subFigs., is defined as the log_2_ fold-change for the number of genes in the gene set with signed proportion extrinsic above the rank cutoff determined by GSEAPY preranked and the number of genes below an identical cutoff (or vice versa if the enrichment was for negative proportion extrinsic divergence). For example, consider an enrichment for positive extrinsic proportion with 500 total genes above the GSEAPY cutoff, 15 of which are imprinted genes. We would then count all imprinted genes in the 500 genes with the most negative signed extrinsic proportion. If, continuing the example, 0 genes of those 500 were in the gene set, the final log_2_ fold-change would be:

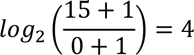

### Transcription factor target enrichment analysis

For the transcription factor (TF) target enrichment analysis, we focused on the averaged signed extrinsic and intrinsic proportion across all cell types to reduce noise, but also performed the analysis on each cell type separately. The analysis was performed as described above, but using the TF Perturbations Followed By Expression database in GSEAPY/MSigDB^27^. We then computed the Spearman correlation between the extrinsic TF target enrichment score for genes with increased expression following *Xbp1* overexpression (from GSEAPY as described above) and the signed proportion extrinsic for *Xbp1.* With regard to the significant enrichment for targets *Nfe2l1* (which codes for the protein Nrf1) for positive intrinsic divergence, the gene is often erroneously referred to as *NRF1* in the database of TF targets and so the significant enrichment for *NRF1* we observe actually refers to *Nfe2l1* (and is referred to as *Nfe2l1*) throughout the text. We repeated this analysis using the TF target enrichment score and signed proportion extrinsic proportion for *Nfe2l2* and *Stat3*. We computed a similar correlation using *Xbp1* and *Nrf1* intrinsic signed proportion and the intrinsic TF target enrichment score for *Xbp1* and *Nrf1* target genes respectively. For *Xbp1*, we also computed the correlation between the signed extrinsic or intrinsic proportion and the enrichment score for the Response to Endoplasmic Reticulum Stress (Response to ER stress) Gene Ontology category^65^. Importantly, a strong correlation between signed extrinsic divergence in *Xbp1* expression and enrichment for it targets or ER stress genes does not necessarily imply a strong correlation between signed intrinsic divergence in *Xbp1* expression and the corresponding enrichments. For example, if *cis*- acting mutations altered the expression of many *Xbp1* target genes, this might lead to weak to no correlation for signed intrinsic divergence even if there were a strong correlation for signed extrinsic divergence.

### Quantification and analysis of immunofluorescence (IF) images

All quantification of fluorescence intensity was performed in Python and all IF images shown in this work were made in ImageJ^66^. We paired chimeras that were generated, fixed, processed, stained, and imaged in the same batch with the same reagents. For all regions of interest, selection, tuning, and parameter selection was performed using only the DAPI and TdTomato channels before measuring fluorescence intensity for c-Jun or Hspa5, ensuring that we were effectively blind to the outcome of the analysis and that the results were not biased.

To quantify fluorescence intensity, we read in the images and drew a rectangular region of interest containing only the medial and lateral ganglionic eminences for chimera pair 1, and only the medial ganglionic eminence for chimera 3 (as the lateral ganglionic eminence had no mouse cells in the rat-like chimera). We then used a Gaussian filter to blur the DAPI and TdTomato channels and masked the images into DAPI positive, DAPI negative, TdTomato positive, and TdTomato negative. We tuned the Gaussian blue sigma parameter and cutoff for the masking by hand until the DAPI mask included all of the area in the region of interest except clearly visible larger regions without DAPI+ nuclei in the region of interest. For the TdTomato signal, we ensured that the mask closely matched the original TdTomato signal in the region of interest. The sigma and masking cutoff values selected in this process were used for all results presented in the text.

We then considered all DAPI+ TdTomato+ pixels to be from donor cells (species-mismatched environment) and all DAPI+ TdT- pixels to be from host cells (species-matched environment), restricted to pixels with non-zero c-Jun or Hspa5 intensity values, and computed the average fluorescent intensity in each set of pixels. For within chimera comparisons, we computed the log_2_ fold-change between the average donor signal and the average host signal as was done for the RNA-seq measurements. We also subtracted the average background signal from a rectangular region with no nuclei from the values computed above before computing log_2_ fold-changes between chimeras. As the sigma parameter and cutoff for the masking were chosen by hand, we tested all possible combinations of three different masking cutoffs (the originally selected cutoff and two other similar values) for DAPI and TdTomato for the rat-like and mouse-like chimeras separately (81 combinations in total) and obtained highly similar log fold-changes (the largest standard error for any log fold-change was 0.0046). These log_2_ fold-changes were then used to compute the extrinsic, intrinsic, and interaction proportions for protein expression divergence as described above for the gene expression divergence measured through scRNA-seq.

To produce the images in the publication the images were first read into ImageJ. Next, channels were assigned colors and the brightness and contrast tuned to visualize the fluorescent signal. Scale bars were then added by synchronizing the windows, drawing horizontal line, and adding a scale bar to each image with the Scale Bar tool. These images were then exported as .png objects.

### Analysis of neuron-progenitor ratios

For the forebrain glutamatergic cells, forebrain GABAergic cells, and the combined spinal cord glutamatergic and GABAergic cells (combined due to the low number of progenitors we detect), we computed the ratio of post-mitotic cells to progenitors (excluding intermediate progenitors) in all four species-environment combinations. We then used these values to compute log_2_ fold-changes and the intrinsic, extrinsic, and interaction proportions in the same way we did for the normalized counts from the scRNA-seq data.

### Analysis of imprinted gene expression

The list of imprinted genes for mice and humans was downloaded from https://www.geneimprint.com/site/home through the “gene by species” tab. To test whether imprinted genes were enriched for high absolute interaction divergence, we created an imprinted gene set consisting of all “confident” and “predicted” imprinted genes in mice from the above resource. Only 2 genes were predicted and not confident, so our results are unaltered by restricting only to confident imprinted genes. We next performed enrichment analysis as described above, including averaging across cell types within nervous and connective tissue, with the following modification. Initially, we simply ranked genes by the absolute interaction divergence and observed very strong enrichment. However, due to the GSEA running sum statistic being initially developed for signed metrics (e.g. log fold-change) we reasoned that this procedure could lead to spuriously strong enrichments^27^. To address this, we rank transformed the list such that the genes with very large absolute interaction divergence have large positive values and genes with absolute interaction divergence near zero had negative values with equally large magnitude. For example, if there were 5,000 input genes, the gene with the highest absolute interaction divergence would have a value of 2,500, the next highest 2,499, etc. The gene with the lowest absolute interaction divergence would have a value of −2,500, the next lowest −2,499, etc. Using this strategy, we still observe strong enrichment (we report the p-values obtained using this strategy in the text and Fig. 7D). For the signed enrichment analysis, we ranked genes by the signed interaction divergence and used GSEAPY preranked with the imprinted gene set.

To analyze the data from chimeric rodent hearts, we downloaded the counts matrices and associated metadata (including the species of origin for each cell) from GEO: https://www.ncbi.nlm.nih.gov/geo/query/acc.cgi?acc=GSE236400^39^. The cell type and other metadata were shared by the original authors of the study.

We then pseudobulked counts by summing across all cells within a cell type for chimera 1, chimera 2, and the wildtype rat sample separately, removed genes with fewer than twenty counts in all samples, and computed counts per million (CPM). We then took the mean of the two chimera replicates and computed the log_2_ fold-change of rat-like environment CPM divided by mouse-like environment CPM. This value was then used in place of interaction divergence for the enrichment analyses described above.

To analyze the data from adult parathyroid glands, we downloaded files containing TPM estimates for each mouse transcript from GEO: https://www.ncbi.nlm.nih.gov/geo/query/acc.cgi?acc=GSE232600. We then collapsed TPM estimates to the gene level by summing the TPM for each transcript assigned to individual genes. As each sample was generated from a very small number of cells (approximately 20), expression estimates for many genes were noisy so we used the median TPM across replicates per condition as our estimate of the expression level of that gene. We removed genes with median TPM less than 5 in all conditions (species-mismatched environment, species-matched environment, and wildtype mice). We restricted to imprinted genes as described above, normalized the expression of each gene, and applied hierarchical clustering with the default parameters in seaborn clustermap. The log fold-change was computed by comparing the median TPMs. All imprinted genes with absolute log_2_ fold-change between expression for species-mismatched and species-matched environment samples greater than 1 were plotted in Fig. S29A.

To analyze the data from human and macaque epiblast, we downloaded the transcripts per million (TPM) matrices for human-macaque chimeric embryos and wildtype human embryos from GEO: https://www.ncbi.nlm.nih.gov/geo/query/acc.cgi?acc=GSE155381 and https://www.ncbi.nlm.nih.gov/geo/query/acc.cgi?acc=GSE109555 respectively. We restricted to cells annotated as epiblast for the human-macaque chimeric data and used Ensembl to identify one-to-one orthologs between crab-eating macaques and humans. For the wildtype human embryos, the raw count matrices and cell type annotations were unavailable. Therefore, we restricted to cells with greater than one TPM for *NANOG*, *SOX2*, or *POU5F1* (also known as *OCT4*) as putative epiblast cells. We then averaged TPM across all cells in each species-environment category, filtered genes that were expressed below 1 TPM in all categories, and computed the log_2_ fold-change of human in a macaque-like environment divided by macaque in a macaque-like environment and human in a macaque-like environment divided by human in a human-like environment after adding one pseudo-TPM to all values. We then performed the enrichment analysis using absolute log_2_ fold-change as described for mouse and rat heart data. Although the comparison of human data across studies undoubtedly introduces batch effects, it is very unlikely that those batch effects would specifically increase the absolute log_2_ fold-change of imprinted genes as a whole.

## Acknowledgements

We thank Alina Xiao, Daniel Pederick, Tom Hindmarsh-Sten, Colleen McLaughlin, Liqun Luo, and other Luo lab members for helpful feedback on the manuscript and immunofluorescence analysis. We also thank Leslie Magtanong, Gabriella Cale, and other members of the Fraser Lab for helpful discussions and feedback on the manuscript. We thank the Stanford Functional Genomics core facility for preparing the scRNA-seq libraries. We thank Xabier López Aranguren and Asier Ullate Agote for sharing their cell type annotations and other metadata for the heart embryonic dataset. We thank Jonathan Pritchard, Richard Schneider, and Vanessa Barrone for helpful feedback on the manuscript. Biorender was used in parts of the main and supplemental figures.

## Authors contributions

TN led all wet-lab work including cell culture, chimera generation and assessment of donor contribution, FACS, preparation of cells for input for scRNA-seq, and immunofluorescence. KJI helped with FACS and sampling of chimeric embryos. CF helped with immunofluorescent staining and optimization. ALS performed all bioinformatic analysis, visualization, validation, and writing of software with guidance from HBF. ALS wrote the main and supplementary text and created Figs. with input from HBF, TN, and HN. ALS and TN wrote the methods section with input from HBF and HN. HBF and HN provided funding and conceived the study.

## Data availability

Sequencing data are available through the gene expression omnibus with accession GSE266218. All code needed to reproduce the analyses described in this study is available at https://github.com/astarr97/Chimera_Pilot.

## Materials and correspondence

Correspondence should be addressed to HBF at. Requests for materials will be fulfilled by the corresponding author.

## List of supplementary materials

Figs. S1-S29

Tables S1-S3

Table S1 contains the estimates of expression levels and intrinsic, extrinsic, and interaction components of each gene passing our filtering criteria in each cell type.

Table S2 contains the full set of signed extrinsic enrichment results.

Table S3 contains the full set of signed extrinsic and intrinsic transcription factor target enrichment results.

Supplementary Text 1

## References and Notes

1. Roux Beiträge zur Entwickelungsmechanik des Embryos. I. Zur Orientirung über einige Probleme der organischen Entwickelung. Z Biol. 21, 411–524.

2. Schlosser, G. (2023). From “self-differentiation” to organoids—the quest for the units of development. Dev. Genes Evol. 10.1007/s00427-023-00711-z.

3. Spemann, H., and Mangold, H. (2001). Induction of embryonic primordia by implantation of organizers from a different species. 1923. Int. J. Dev. Biol. 45, 13–38.

4. Le Douarin, N.M. (1980). The ontogeny of the neural crest in avian embryo chimaeras. Nature 286, 663–669. 10.1038/286663a0.

5. Twitty, Victor C. (1936). CORRELATED GENETIC AND EMBRYOLOGICAL EXPERIMENTS ON TRITURUS. I and II. The Journal of Experimental Zoology 74.

6. Schneider, R.A. (2018). Neural crest and the origin of species-specific pattern. genesis *56*, e23219. 10.1002/dvg.23219.

7. Le Douarin N, McLaren A. (1984). Chimeras in developmental biology (Academic Press).

8. Suchy, F., and Nakauchi, H. (2018). Interspecies chimeras. Curr. Opin. Genet. Dev. 52, 36–41. 10.1016/j.gde.2018.05.007.

9. Stepien, B.K., Naumann, R., Holtz, A., Helppi, J., Huttner, W.B., and Vaid, S. (2020). Lengthening Neurogenic Period during Neocortical Development Causes a Hallmark of Neocortex Expansion. Curr. Biol. 30, 4227–4237.e5. 10.1016/j.cub.2020.08.046.

10. Eames, B.F., and Schneider, R.A. (2005). Quail-duck chimeras reveal spatiotemporal plasticity in molecular and histogenic programs of cranial feather development. Dev. Camb. Engl. 132, 1499–1509. 10.1242/dev.01719.

11. Schneider, R.A., and Helms, J.A. (2003). The Cellular and Molecular Origins of Beak Morphology. Science 299, 565–568. 10.1126/science.1077827.

12. Wittkopp, P.J., and Kalay, G. (2012). Cis-regulatory elements: molecular mechanisms and evolutionary processes underlying divergence. Nat. Rev. Genet. 13, 59–69. 10.1038/nrg3095.

13. Wittkopp, P.J., Haerum, B.K., and Clark, A.G. (2004). Evolutionary changes in cis and trans gene regulation. Nature 430, 85–88. 10.1038/nature02698.

14. Coolon, J.D., McManus, C.J., Stevenson, K.R., Graveley, B.R., and Wittkopp, P.J. (2014). Tempo and mode of regulatory evolution in *Drosophila*. Genome Res. 24, 797–808. 10.1101/gr.163014.113.

15. Liu, X., Li, Y.I., and Pritchard, J.K. (2019). Trans Effects on Gene Expression Can Drive Omnigenic Inheritance. Cell 177, 1022–1034.e6. 10.1016/j.cell.2019.04.014.

16. Albert, F.W., Bloom, J.S., Siegel, J., Day, L., and Kruglyak, L. (2018). Genetics of trans-regulatory variation in gene expression. eLife 7, e35471. 10.7554/eLife.35471.

17. Wang, Q., Jia, Y., Wang, Y., Jiang, Z., Zhou, X., Zhang, Z., Nie, C., Li, J., Yang, N., and Qu, L. (2019). Evolution of cis- and trans-regulatory divergence in the chicken genome between two contrasting breeds analyzed using three tissue types at one-day-old. BMC Genomics 20, 933. 10.1186/s12864-019-6342-5.

18. Huang, J., He, B., Yang, X., Long, X., Wei, Y., Gao, Y., Fang, Y., Ying, W., Wang, Z., Li, C., et al. (2023). Interspecies blastocyst complementation generates functional rat cell-derived forebrain tissues in mice (Developmental Biology) 10.1101/2023.04.13.536774.

19. Kim-Hellmuth, S., Aguet, F., Oliva, M., Muñoz-Aguirre, M., Kasela, S., Wucher, V., Castel, S.E., Hamel, A.R., Viñuela, A., Roberts, A.L., et al. (2020). Cell type–specific genetic regulation of gene expression across human tissues. Science 369, eaaz8528. 10.1126/science.aaz8528.

20. The GTEx Consortium, Aguet, F., Anand, S., Ardlie, K.G., Gabriel, S., Getz, G.A., Graubert, A., Hadley, K., Handsaker, R.E., Huang, K.H., et al. (2020). The GTEx Consortium atlas of genetic regulatory effects across human tissues. Science 369, 1318–1330. 10.1126/science.aaz1776.

21. GTEx Consortium (2017). Genetic effects on gene expression across human tissues. Nature 550, 204–213. 10.1038/nature24277.

22. Metzger, B.P.H., Duveau, F., Yuan, D.C., Tryban, S., Yang, B., and Wittkopp, P.J. (2016). Contrasting Frequencies and Effects of *cis* - and *trans* -Regulatory Mutations Affecting Gene Expression. Mol. Biol. Evol. 33, 1131–1146. 10.1093/molbev/msw011.

23. Hoter, A., El-Sabban, M., and Naim, H. (2018). The HSP90 Family: Structure, Regulation, Function, and Implications in Health and Disease. Int. J. Mol. Sci. 19, 2560. 10.3390/ijms19092560.

24. Zhu, Y., Sun, D., Jakovcevski, M., and Jiang, Y. (2020). Epigenetic mechanism of SETDB1 in brain: implications for neuropsychiatric disorders. Transl. Psychiatry 10, 115. 10.1038/s41398-020-0797-7.

25. Park, S.-M., Kang, T.-I., and So, J.-S. (2021). Roles of XBP1s in Transcriptional Regulation of Target Genes. Biomedicines 9, 791. 10.3390/biomedicines9070791.

26. Gregor, M.F., Misch, E.S., Yang, L., Hummasti, S., Inouye, K.E., Lee, A.-H., Bierie, B., and Hotamisligil, G.S. (2013). The role of adipocyte XBP1 in metabolic regulation during lactation. Cell Rep. 3, 1430–1439. 10.1016/j.celrep.2013.03.042.

27. Subramanian, A., Tamayo, P., Mootha, V.K., Mukherjee, S., Ebert, B.L., Gillette, M.A., Paulovich, A., Pomeroy, S.L., Golub, T.R., Lander, E.S., et al. (2005). Gene set enrichment analysis: A knowledge-based approach for interpreting genome-wide expression profiles. Proc. Natl. Acad. Sci. 102, 15545–15550. 10.1073/pnas.0506580102.

28. Ku, H.-C., and Cheng, C.-F. (2020). Master Regulator Activating Transcription Factor 3 (ATF3) in Metabolic Homeostasis and Cancer. Front. Endocrinol. 11, 556. 10.3389/fendo.2020.00556.

29. Adachi, Y., Yamamoto, K., Okada, T., Yoshida, H., Harada, A., and Mori, K. (2008). ATF6 is a transcription factor specializing in the regulation of quality control proteins in the endoplasmic reticulum. Cell Struct. Funct. 33, 75–89. 10.1247/csf.07044.

30. Yu, W., Wang, B., Zhou, L., and Xu, G. (2021). Endoplasmic Reticulum Stress-Mediated p62 Downregulation Inhibits Apoptosis via c-Jun Upregulation. Biomol. Ther. 29, 195–204. 10.4062/biomolther.2020.089.

31. Sha, Z., and Goldberg, A.L. (2014). Proteasome-Mediated Processing of Nrf1 Is Essential for Coordinate Induction of All Proteasome Subunits and p97. Curr. Biol. 24, 1573–1583. 10.1016/j.cub.2014.06.004.

32. Vangala, J.R., Dudem, S., Jain, N., and Kalivendi, S.V. (2014). Regulation of PSMB5 Protein and β Subunits of Mammalian Proteasome by Constitutively Activated Signal Transducer and Activator of Transcription 3 (STAT3). J. Biol. Chem. 289, 12612–12622. 10.1074/jbc.M113.542829.

33. He, F., Ru, X., and Wen, T. (2020). NRF2, a Transcription Factor for Stress Response and Beyond. Int. J. Mol. Sci. 21, 4777. 10.3390/ijms21134777.

34. Wang, J., Lee, J., Liem, D., and Ping, P. (2017). HSPA5 Gene encoding Hsp70 chaperone BiP in the endoplasmic reticulum. Gene 618, 14–23. 10.1016/j.gene.2017.03.005.

35. Juan, A.M., Foong, Y.H., Thorvaldsen, J.L., Lan, Y., Leu, N.A., Rurik, J.G., Li, L., Krapp, C., Rosier, C.L., Epstein, J.A., et al. (2022). Tissue-specific Grb10/Ddc insulator drives allelic architecture for cardiac development. Mol. Cell 82, 3613–3631.e7. 10.1016/j.molcel.2022.08.021.

36. Giannoukakis, N., Deal, C., Paquette, J., Goodyer, C.G., and Polychronakos, C. (1993). Parental genomic imprinting of the human IGF2 gene. Nat. Genet. 4, 98–101. 10.1038/ng0593-98.

37. Moore, G.E., Ishida, M., Demetriou, C., Al-Olabi, L., Leon, L.J., Thomas, A.C., Abu-Amero, S., Frost, J.M., Stafford, J.L., Chaoqun, Y., et al. (2015). The role and interaction of imprinted genes in human fetal growth. Philos. Trans. R. Soc. B Biol. Sci. 370, 20140074. 10.1098/rstb.2014.0074.

38. Glaser, J., Iranzo, J., Borensztein, M., Marinucci, M., Gualtieri, A., Jouhanneau, C., Teissandier, A., Gaston-Massuet, C., and Bourc’his, D. (2022). The imprinted Zdbf2 gene finely tunes control of feeding and growth in neonates. eLife 11, e65641. 10.7554/eLife.65641.

39. Coppiello, G., Barlabé, P., Moya-Jódar, M., Abizanda, G., Pogontke, C., Barreda, C., Iglesias, E., Linares, J., Arellano-Viera, E., Larequi, E., et al. (2023). Generation of heart and vascular system in rodents by blastocyst complementation. Dev. Cell 58, 2881–2895.e7. 10.1016/j.devcel.2023.10.008.

40. Kano, M., Mizuno, N., Sato, H., Kimura, T., Hirochika, R., Iwasaki, Y., Inoshita, N., Nagano, H., Kasai, M., Yamamoto, H., et al. (2023). Functional calcium-responsive parathyroid glands generated using single-step blastocyst complementation. Proc. Natl. Acad. Sci. 120, e2216564120. 10.1073/pnas.2216564120.

41. Zhou, F., Wang, R., Yuan, P., Ren, Y., Mao, Y., Li, R., Lian, Y., Li, J., Wen, L., Yan, L., et al. (2019). Reconstituting the transcriptome and DNA methylome landscapes of human implantation. Nature 572, 660–664. 10.1038/s41586-019-1500-0.

42. Tan, T., Wu, J., Si, C., Dai, S., Zhang, Y., Sun, N., Zhang, E., Shao, H., Si, W., Yang, P., et al. (2021). Chimeric contribution of human extended pluripotent stem cells to monkey embryos ex vivo. Cell 184, 2020–2032.e14. 10.1016/j.cell.2021.03.020.

43. Masaki, H., and Nakauchi, H. (2017). Interspecies chimeras for human stem cell research. Development 144, 2544–2547. 10.1242/dev.151183.

44. Ramani, B., Rose, I.V.L., Pan, A., Tian, R., Ma, K., Palop, J.J., and Kampmann, M. (2023). Scalable, cell type-selective, AAV-based in vivo CRISPR screening in the mouse brain (Genomics) 10.1101/2023.06.13.544831.

45. Ciceri, G., Baggiolini, A., Cho, H.S., Kshirsagar, M., Benito-Kwiecinski, S., Walsh, R.M., Aromolaran, K.A., Gonzalez-Hernandez, A.J., Munguba, H., Koo, S.Y., et al. (2024). An epigenetic barrier sets the timing of human neuronal maturation. Nature 626, 881–890. 10.1038/s41586-023-06984-8.

46. Piedrahita, J.A. (2011). The role of imprinted genes in fetal growth abnormalities. Birt. Defects Res. A. Clin. Mol. Teratol. 91, 682–692. 10.1002/bdra.20795.

47. Ying, Q.-L., Stavridis, M., Griffiths, D., Li, M., and Smith, A. (2003). Conversion of embryonic stem cells into neuroectodermal precursors in adherent monoculture. Nat. Biotechnol. 21, 183–186. 10.1038/nbt780.

48. Yamaguchi, T., Hamanaka, S., and Nakauchi, H. (2014). The generation and maintenance of rat induced pluripotent stem cells. Methods Mol. Biol. Clifton NJ 1210, 143–150. 10.1007/978-1-4939-1435-7_11.

49. Brownstein, D.G. (2003). Manipulating the Mouse Embryo: A Laboratory Manual . Third Edition. *By* Andras Nagy, Marina Gertsenstein, Kristina Vintersten, and Richard Behringer. *Cold Spring Harbor (New York)*: Cold Spring Harbor Laboratory Press . $195.00 (hardcover); $115.00 (paper). x + 764 p; ill.; index. ISBN: 0-87969-574-9 (hc); 0-87969-591-9 (pb). 2003. Q. Rev. Biol. *78*, 365–365. 10.1086/380032.

50. Yates, A.D., Allen, J., Amode, R.M., Azov, A.G., Barba, M., Becerra, A., Bhai, J., Campbell, L.I., Carbajo Martinez, M., Chakiachvili, M., et al. (2022). Ensembl Genomes 2022: an expanding genome resource for non-vertebrates. Nucleic Acids Res. 50, D996–D1003. 10.1093/nar/gkab1007.

51. Wolf, F.A., Angerer, P., and Theis, F.J. (2018). SCANPY: large-scale single-cell gene expression data analysis. Genome Biol. 19, 15. 10.1186/s13059-017-1382-0.

52. Korsunsky, I., Millard, N., Fan, J., Slowikowski, K., Zhang, F., Wei, K., Baglaenko, Y., Brenner, M., Loh, P., and Raychaudhuri, S. (2019). Fast, sensitive and accurate integration of single-cell data with Harmony. Nat. Methods 16, 1289–1296. 10.1038/s41592-019-0619-0.

53. Traag, V.A., Waltman, L., and Van Eck, N.J. (2019). From Louvain to Leiden: guaranteeing well-connected communities. Sci. Rep. 9, 5233. 10.1038/s41598-019-41695-z.

54. Di Bella, D.J., Habibi, E., Stickels, R.R., Scalia, G., Brown, J., Yadollahpour, P., Yang, S.M., Abbate, C., Biancalani, T., Macosko, E.Z., et al. (2021). Molecular logic of cellular diversification in the mouse cerebral cortex. Nature 595, 554–559. 10.1038/s41586-021-03670-5.

55. Lipiec, M.A., Bem, J., Koziński, K., Chakraborty, C., Urban-Ciećko, J., Zajkowski, T., Dąbrowski, M., Szewczyk, Ł.M., Toval, A., Ferran, J.L., et al. (2020). TCF7L2 regulates postmitotic differentiation programs and excitability patterns in the thalamus. Development, dev.190181. 10.1242/dev.190181.

56. Witschi, R., Johansson, T., Morscher, G., Scheurer, L., Deschamps, J., and Zeilhofer, H.U. (2010). *Hoxb8-Cre* mice: A tool for brain-sparing conditional gene deletion. genesis 48, 596–602. 10.1002/dvg.20656.

57. Lee, J., Rabbani, C.C., Gao, H., Steinhart, M.R., Woodruff, B.M., Pflum, Z.E., Kim, A., Heller, S., Liu, Y., Shipchandler, T.Z., et al. (2020). Hair-bearing human skin generated entirely from pluripotent stem cells. Nature 582, 399–404. 10.1038/s41586-020-2352-3.

58. Cardoso-Moreira, M., Halbert, J., Valloton, D., Velten, B., Chen, C., Shao, Y., Liechti, A., Ascenção, K., Rummel, C., Ovchinnikova, S., et al. (2019). Gene expression across mammalian organ development. Nature 571, 505–509. 10.1038/s41586-019-1338-5.

59. Virtanen, P., Gommers, R., Oliphant, T.E., Haberland, M., Reddy, T., Cournapeau, D., Burovski, E., Peterson, P., Weckesser, W., Bright, J., et al. (2020). SciPy 1.0: fundamental algorithms for scientific computing in Python. Nat. Methods 17, 261–272. 10.1038/s41592-019-0686-2.

60. Zeng, T., Spence, J.P., Mostafavi, H., and Pritchard, J.K. (2023). Bayesian estimation of gene constraint from an evolutionary model with gene features (Genetics) 10.1101/2023.05.19.541520.

61. The ENCODE Project Consortium, Abascal, F., Acosta, R., Addleman, N.J., Adrian, J., Afzal, V., Ai, R., Aken, B., Akiyama, J.A., Jammal, O.A., et al. (2020). Expanded encyclopaedias of DNA elements in the human and mouse genomes. Nature 583, 699–710. 10.1038/s41586-020-2493-4.

62. Yanai, I., Benjamin, H., Shmoish, M., Chalifa-Caspi, V., Shklar, M., Ophir, R., Bar-Even, A., Horn-Saban, S., Safran, M., Domany, E., et al. (2005). Genome-wide midrange transcription profiles reveal expression level relationships in human tissue specification. Bioinforma. Oxf. Engl. 21, 650–659. 10.1093/bioinformatics/bti042.

63. Giurgiu, M., Reinhard, J., Brauner, B., Dunger-Kaltenbach, I., Fobo, G., Frishman, G., Montrone, C., and Ruepp, A. (2019). CORUM: the comprehensive resource of mammalian protein complexes—2019. Nucleic Acids Res. 47, D559–D563. 10.1093/nar/gky973.

64. Rouillard, A.D., Gundersen, G.W., Fernandez, N.F., Wang, Z., Monteiro, C.D., McDermott, M.G., and Ma’ayan, A. (2016). The harmonizome: a collection of processed datasets gathered to serve and mine knowledge about genes and proteins. Database 2016, baw100. 10.1093/database/baw100.

65. Ashburner, M., Ball, C.A., Blake, J.A., Botstein, D., Butler, H., Cherry, J.M., Davis, A.P., Dolinski, K., Dwight, S.S., Eppig, J.T., et al. (2000). Gene Ontology: tool for the unification of biology. Nat. Genet. 25, 25–29. 10.1038/75556.

66. Schindelin, J., Arganda-Carreras, I., Frise, E., Kaynig, V., Longair, M., Pietzsch, T., Preibisch, S., Rueden, C., Saalfeld, S., Schmid, B., et al. (2012). Fiji: an open-source platform for biological-image analysis. Nat. Methods 9, 676–682. 10.1038/nmeth.2019.

