## Supplemental Text 1 for "Disentangling cell-intrinsic and extrinsic factors underlying evolution"

***The effects of developmentally dynamic gene expression on estimates of extrinsic and intrinsic gene expression divergence***

An important aspect of gene expression we have not explored in this study is changes in gene expression over developmental time (developmentally dynamic expression).  In addition, the trajectory of gene expression can diverge between species.  To explore how developmentally dynamic expression influences the results presented here, we analyzed bulk RNA-seq data from a time course of mouse and rat development, focusing on E11.5, E13.5, E15.5 in mouse and E11, E13, and E15 in rat.  We analyze how a combination of intrinsic divergence and developmentally dynamic gene expression can appear as having purely interaction, purely extrinsic, or reinforcing/opposing extrinsic and intrinsic divergence in our study.

***Intrinsic temporal shifts in gene expression could appear as purely extrinsic divergence***

First, a “temporal shift” (i.e. a shift against the global shift in gene expression associated with slower development of rats compared to mice) in gene expression could appear as extrinsic divergence in our study.  As mentioned above, rat development proceeds at a slower pace than mouse development.  Therefore, if there is no divergence in the expression trajectory of a gene beyond this global change, we would expect rat expression to differ from mouse expression at identical (but not stage-matched) timepoints for developmentally dynamic genes.  However, the expression trajectories of some genes might not shift with this global change.  This would result in the same expression trajectory (i.e. increasing or decreasing) in both species, but with similar expression levels at identical time points (Supplemental Figure 3A-B).  If this occurs through purely intrinsic mechanisms, we would observe purely extrinsic divergence in our study.  To test whether this occurs frequently, we identified genes with similar expression trajectories in mice and rats but small differences in expression at both E13.5 and E15.5 (absolute log fold-change less than 0.25).  Importantly, when adding the restriction that the log fold-change in expression between E13.5 and E15.5 within species be greater than 0.5, less than 1% of genes fulfill these criteria suggesting the kind of temporal shift in expression hypothesized here is very rare.  Therefore, we proceeded only with the restriction of low divergence between species at identical timepoints and similar expression trajectories between species.  Overall, we find weak or no enrichment for genes with high proportion extrinsic divergence in this set of genes with evidence for a temporal shift (no cell types with enrichments in the expected direction with p < 0.1, Supplemental Figure 4A).  In addition, we would expect that temporal shift genes that decrease over time would be enriched for negative extrinsic divergence and vice versa (Supplemental Figure 3A-B).  Here again we find only weak enrichments (Supplemental Figure 4B).

***Intrinsically opposite gene expression trajectories in mouse and rat could appear as interactions between intrinsic and extrinsic divergence***

Another form of temporal divergence in gene expression is when the expression of a gene increases over time in one species but decreases over time in another species.  For example, if a gene is intrinsically decreasing over time in mouse and increasing over time in rat (or vice versa), this could appear as purely an interaction between extrinsic and intrinsic divergence in our study (Supplemental Figure 5A-B).  If this appreciably contributes to the inflation of the interaction component in our study, we would expect that genes with high interaction proportions would be enriched for opposing trajectories in the time course data.  However, across all brain cell types in our study, we find little to no evidence for this enrichment (p < 0.1 in one cell type, Supplemental Figure 6A).  In addition, we can make a stronger prediction about the sign of interaction divergence if this confounder plays a major role.  If gene expression is decreasing in mouse and increasing in rat, then this would lead to higher expression in species-matched environments (i.e. negative interaction divergence) due to the slower development of rat cells (Supplemental Figure 5A).  As a result, this category of genes should be enriched for negative interaction divergence.  Similarly, genes that increase in expression over time in mouse but decrease over time in rat would lead to higher expression in species-mismatched environments (i.e. positive interaction divergence, Supplemental Figure 5B).  This category should then be enriched for positive interaction divergence.  Again, we find weak to no enrichment in the expected direction (p < 0.1 for one cell type for increasing in mouse, no cell types for decreasing in mouse Supplemental Figure 6B).

***Conserved expression trajectories and intrinsic divergence could appear as opposing or reinforcing extrinsic and intrinsic divergence***

Finally, intrinsic divergence coupled with a conserved expression trajectory can lead to the appearance of opposing or reinforcing extrinsic or intrinsic divergence. For example, if a gene is increasing in expression over time in both species and is intrinsically more highly expressed in mouse cells, this can appear as opposing extrinsic and intrinsic divergence in our study (Supplemental Figure 7A).  In general, we would expect that both opposing and reinforcing genes would be enriched for genes with conserved expression trajectories in mouse and rat development (Supplemental Figure 7A-D).  In addition, we can make the more specific hypothesis that genes that are intrinsically higher in mouse cells and increasing over time as well as genes that are intrinsically higher in rat and decreasing over time should be enriched for opposing extrinsic and intrinsic divergence (Supplemental Figure 7A-B).  On the other hand, genes that are intrinsically higher in mouse cells and decreasing over time as well as genes that are intrinsically higher in rat and increasing over time should be enriched for reinforcing genes (Supplemental Figure 7C-D).  However, we find limited evidence for either hypothesis (p < 0.1 in one cell type for each enrichment respectively Supplemental Figure 8A-B**).**  Although this suggests that intrinsic divergence coupled with similar changes in expression over time is not the primary contributor to opposing or reinforcing extrinsic and intrinsic divergence, there are many genes with conserved expression trajectories suggesting that this area in particular should be explored further in future studies.

Overall, these results suggest that a combination of intrinsic divergence and developmentally dynamic expression do not overly inflate our estimates of extrinsic and interaction divergence.  However, we have only discussed intrinsic divergence in conjunction with developmentally dynamic gene expression.  Various other complex combinations of divergence can instead inflate estimates of intrinsic divergence.  For example, extrinsic divergence itself can be partially responsible for global shifts in the trajectory of gene expression further adding to complexity.  Data from developmental time courses in reciprocal chimeras will undoubtedly provide valuable insight into how intrinsic, extrinsic, and temporal divergence interact and be vital in developing a more complete understanding of the molecular mechanisms underlying gene expression divergence.

***The spatial distribution of donor cells could lead to differences in estimates of intrinsic and extrinsic divergence between cell types***

In our study, different cell types and tissues have different percentages of donor cells and different spatial distributions of donor cells.  For example, rat donor cells make up a much larger portion of the developing cortex than the ganglionic eminences (Figure 1F).  For rat cells surrounded by other rat cells in the developing cortex, the local environment is likely more rat-like than for rat cells surrounded by mouse cells.  This might result in underestimation of the extrinsic component of gene expression divergence and overestimation of the intrinsic component for forebrain glutamatergic progenitors and forebrain glutamatergic neurons compared to forebrain GABAergic cells.  While we do observe a larger intrinsic component for forebrain glutamatergic neurons compared to their GABAergic counterparts, the intrinsic and extrinsic estimates for glutamatergic and GABAergic progenitors are highly similar (Figure 2G).  With the scRNA-seq analysis in this study, the spatial distribution of the donor cells in each tissue is lost so we are unable to directly test the extent to which the species of the cells adjacent to each sequences cell affects estimates of extrinsic and intrinsic divergence.  In the future, using spatial transcriptomics will enable more accurate estimation of extrinsic and intrinsic divergence as well as further division of the extrinsic component of gene expression divergence into global and local effects.
