## Supplemental Figures for "Disentangling cell-intrinsic and extrinsic factors underlying evolution"

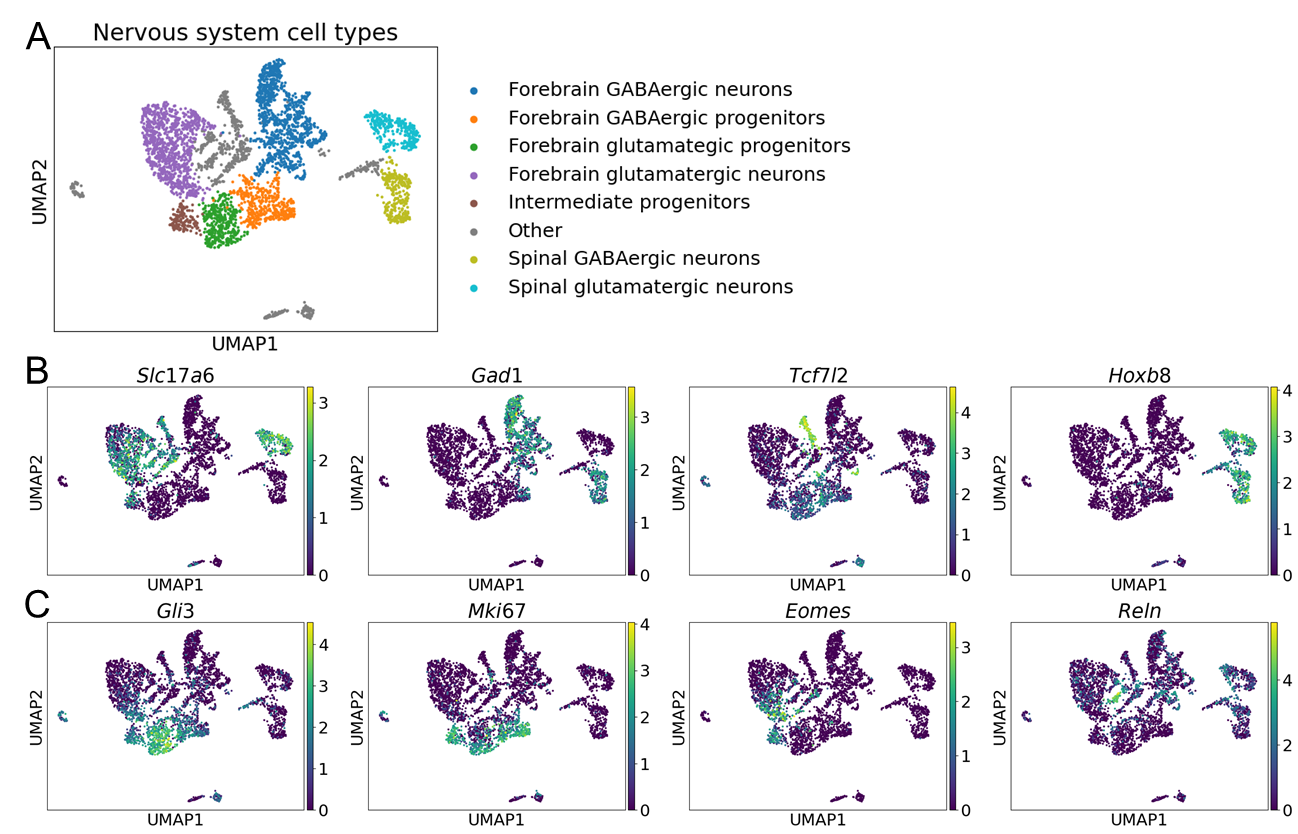

**Fig. S1: Nervous system cell type annotations.** **A)** Uniform manifold approximation (UMAP) of all nervous system cell types. Cell types analyzed in this study are labeled and all other cell types are categorized as “Other”. **B)** Marker genes used to classify nervous system cell types. From left to right: expression of *Slc17a6*, a marker of glutamatergic cells, expression of *Gad1,* a marker of GABAergic cells, expression of *Tcf7l2*, a marker for midbrain neurons (which were not analyzed), expression of *Hoxb8*, a marker for spinal neurons. Each point is colored by the log normalized expression in the cell represented by the point. **C)** Additional marker genes used to classify cell types. From left to right: expression of *Gli3*, which is highly expressed in forebrain glutamatergic progenitors, expression of *Mki67*, a marker of cycling cells, expression of *Eomes*, a marker for intermediate progenitors, expression of *Reln*, a marker of Cajal-Retzius cells which were excluded from further analysis.

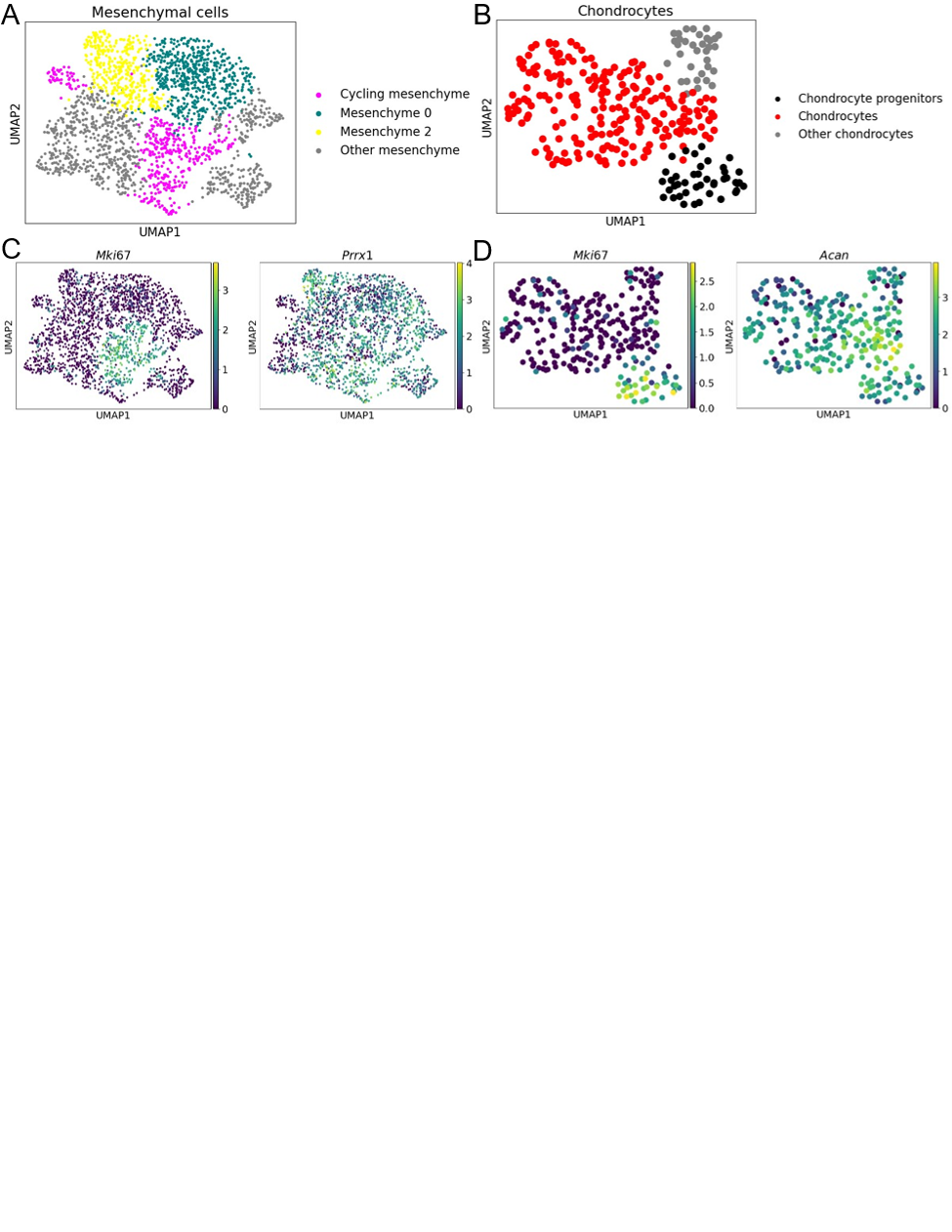

**Fig. S2: Connective tissue cell type annotations. A)** UMAP of mesenchymal cell subtypes. UMAP is used purely for visualization and the proximity of different clusters in UMAP space cannot be used as a proxy for the true similarity between groups of cells. The small group of cycling mesenchymal cells with lower *Mki67* expression in the upper left is most similar in gene expression to the large group of cycling mesenchymal cells with higher *Mki67* expression as determined by leiden clustering and so are included with that cluster. Some cell types excluded from further analysis are labeled as Other. **B)** UMAP of chondrocyte subtypes. Cycling chondrocytes and other chondrocytes were excluded from further analysis. **C)** Expression of *Mki67*, a marker of cycling cells, and *Prrx1*, a marker of mesenchymal cells, in mesenchymal cells. Each point is colored by the log normalized expression in the cell represented by the point. **D)** Expression of *Mki67* and *Acan*, a marker of chondrocytes, in chondrocytes.

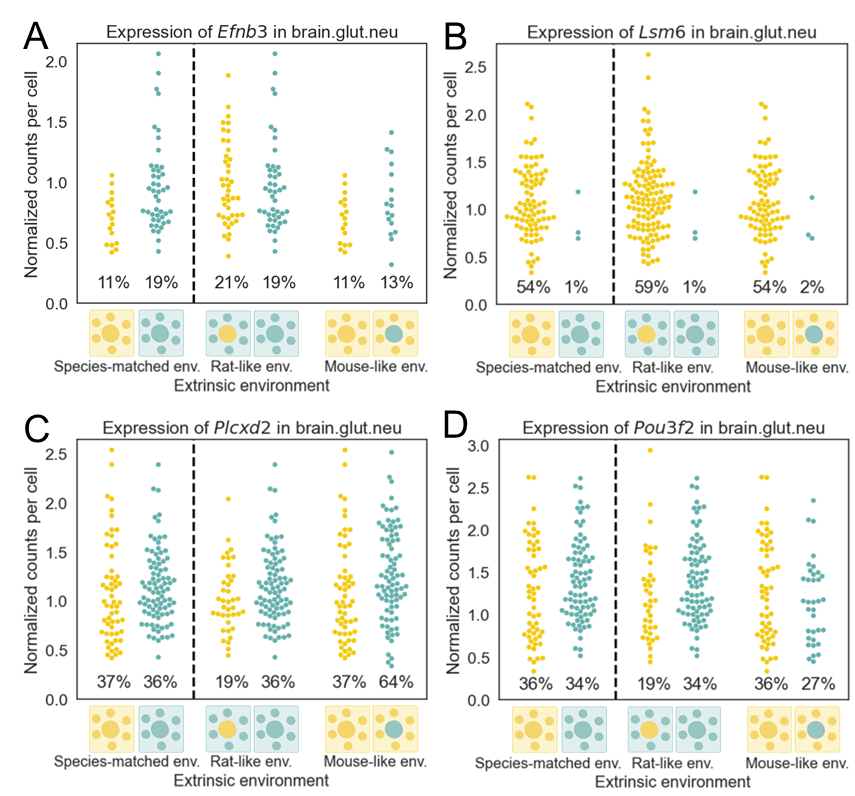

**Fig. S3: Per-cell expression distribution for example genes.** Each swarm of points shows the normalized counts for a gene in each forebrain glutamatergic neuronal cells with non-zero counts for that gene. The percentage near the bottom of the plot indicates the percentage of cells with non-zero counts for that gene. A) Per-cell expression for *Efnb3*. B) Per-cell expression for *Lsm6*. C) Per-cell expression for *Plcxd2*. D) Per-cell expression for *Pou3f2*.

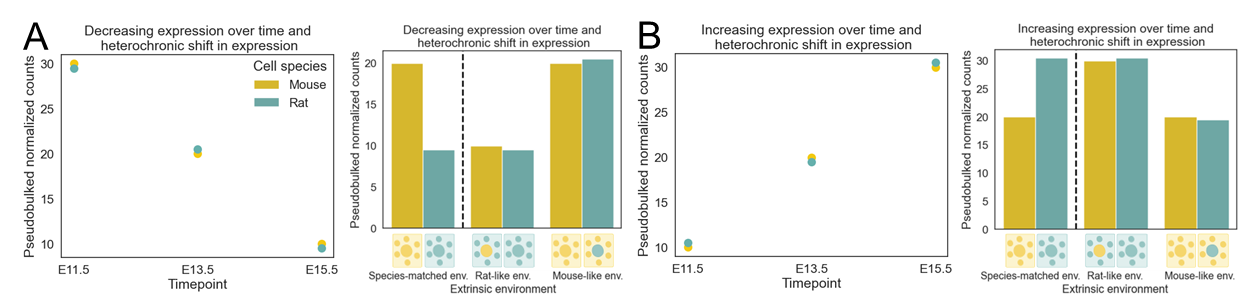

**Fig. S4: Conceptual outline of the interplay of intrinsic divergence and heterochronic shifts in gene expression. A)** Expression of a hypothetical gene across development. The gene decreases in expression over development but has very similar expression in mice and rats at the same embryonic timepoints, going against the global difference in developmental rate between species. In the absence of extrinsic or interaction divergence, this gene would appear to have purely extrinsic divergence and higher expression in a mouse-like environment. **B)** The same as in (A) but showing a gene with increasing expression over time and a heterochronic shift. In the absence of extrinsic or interaction divergence, this gene would appear to have purely extrinsic divergence and higher expression in a rat-like environment.

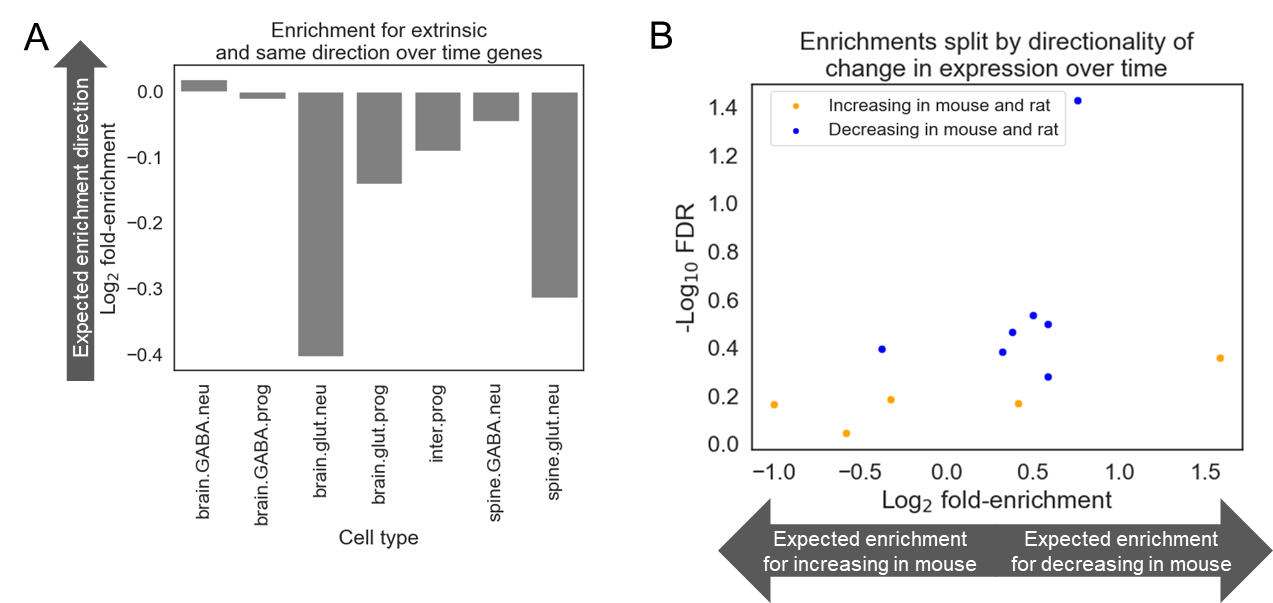

**Fig. S5: Enrichment analysis for heterochronic shift genes.** See Methods for how heterochronic shift genes were defined. **A)** Enrichment analysis for genes with similar expression trajectories during embryonic brain development and similar expression levels at E13.5 and E15.5 in mice and rats and proportion extrinsic divergence. The y-axis shows the log_2_ fold-enrichment and the x-axis corresponds to cell type. The arrow shows the expected enrichment if intrinsic heterochronic shift genes were inflating estimates of extrinsic divergence. **B)** Enrichment analysis for genes with similar expression trajectories during embryonic brain development and similar expression levels at E13.5 and E15.5 in mice and rats and signed proportion extrinsic divergence. The x-axis is the log_2_ fold-enrichment and the y-axis is the -log­_10_(FDR). The arrow shows the expected enrichment if intrinsic heterochronic shift genes were inflating estimates of extrinsic divergence. Each point corresponds to the enrichment in a central nervous system cell type and each cell type is represented twice, once as an blue dot for genes that are decreasing in mouse and rat (and have similar expression levels at E13.5 and E15.5 in mouse and rat) and once as an orange dot for genes that are increasing in mouse and rat (and have similar expression levels at E13.5 and E15.5 in mouse and rat).

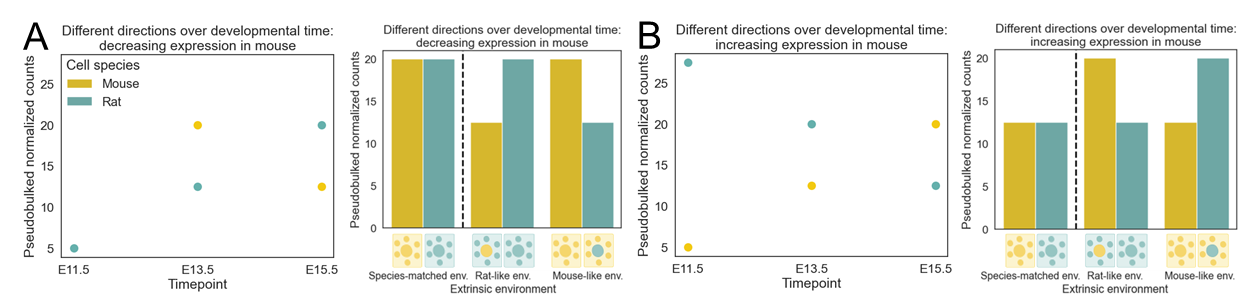

**Fig. S6: Conceptual outline of the interplay of intrinsic divergence and switches in the trajectory of gene expression during development between species. A)** Expression of a hypothetical gene across development. The gene increases in expression over time in rats, but decreases in expression over time in mice. In the absence of extrinsic or interaction divergence, this gene would appear to have purely interaction divergence and higher expression in species-matched environments. **B)** The same as in (A) but showing a gene with increasing expression over time in mice and decreasing expression over time in rats. In the absence of extrinsic or interaction divergence, this gene would appear to have purely interaction divergence and higher expression in species-mismatched environments.

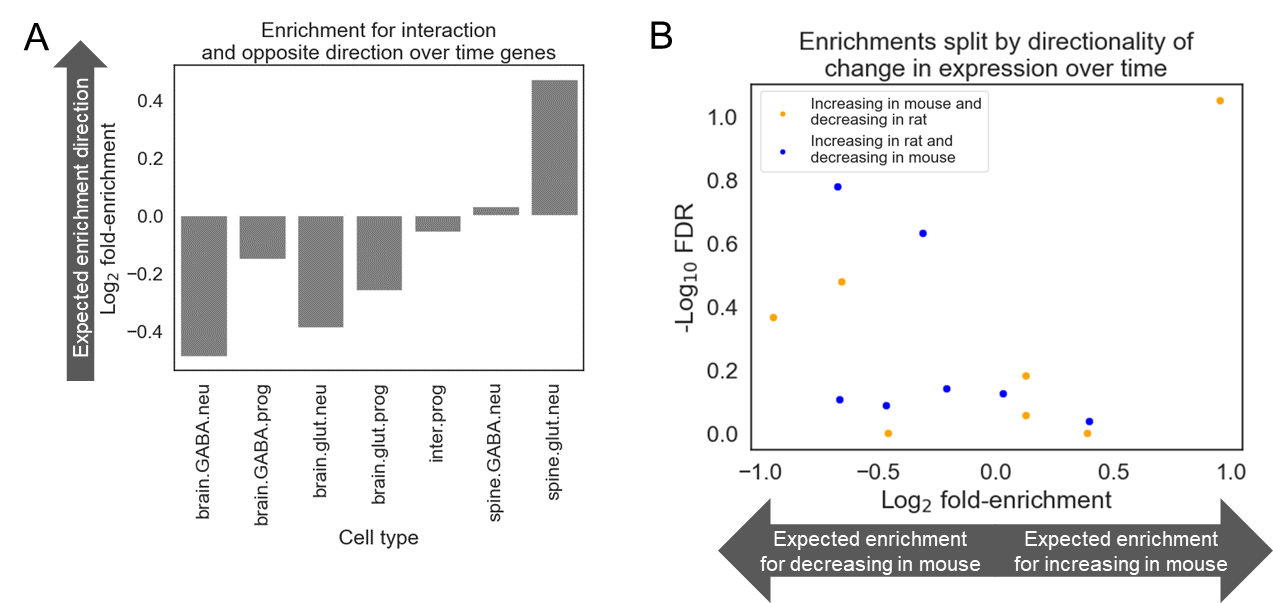

**Fig. S7: Enrichment analysis for genes with switched trajectories of gene expression during development in mice and rats.** See Methods for how genes with switches in gene expression trajectory between species were defined. **A)** Enrichment analysis for genes with switches in gene expression during embryonic brain development and proportion interaction divergence. The y-axis shows the log_2_ fold-enrichment and the x-axis corresponds to cell type. The arrow shows the expected enrichment if genes with the opposite gene expression trajectory between mice and rats were inflating estimates of interaction divergence. **B)** Enrichment analysis for genes with switches in gene expression during embryonic brain development between species and signed proportion interaction divergence. The x-axis is log_2_ fold-enrichment and the y-axis is the -log_10_(FDR). The arrow shows the expected enrichment if genes with the opposite gene expression trajectory between mice and rats were inflating estimates of interaction divergence. Each point corresponds to the enrichment in a central nervous system cell type and each cell type is represented twice, once as a blue dot for genes that are increasing over time in rat and decreasing over time in mouse, and once as an orange dot for genes that are decreasing over time in rat and increasing over time in mouse.

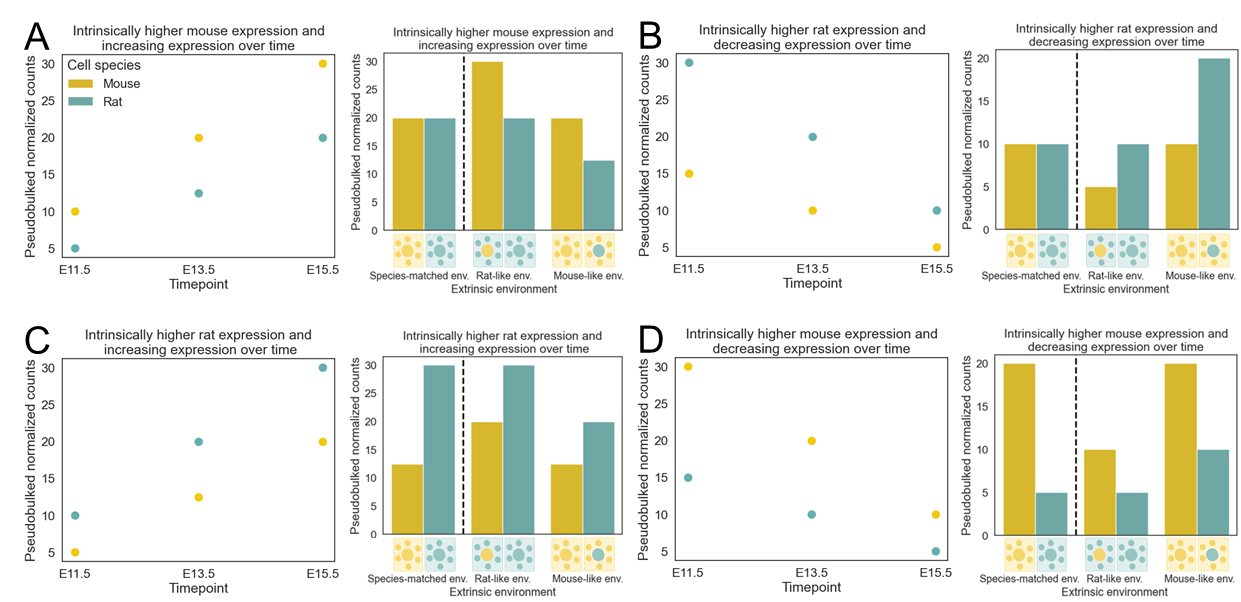

**Fig. S8: Conceptual outline of the interplay of intrinsic divergence and conserved gene expression trajectories. A)** Expression of a hypothetical gene across development. The gene increases in expression over development in both mice and rats, but is intrinsically more highly expressed in mice. In the absence of extrinsic or interaction divergence, this gene would appear to have opposing extrinsic and intrinsic divergence. **B)** The same as in (A) but showing a gene with decreasing expression over time in both species and intrinsically higher expression in rat cells. In the absence of extrinsic or interaction divergence, this gene would appear to have opposing extrinsic and intrinsic divergence. **C)** The same as in (A) but showing a gene with increasing expression over time in both species and intrinsically higher expression in rat cells. In the absence of extrinsic or interaction divergence, this gene would appear to have reinforcing extrinsic and intrinsic divergence. **D)** The same as in (A) but showing a gene with decreasing expression over time in both species and intrinsically higher expression in mouse cells. In the absence of extrinsic or interaction divergence, this gene would appear to have reinforcing extrinsic and intrinsic divergence.

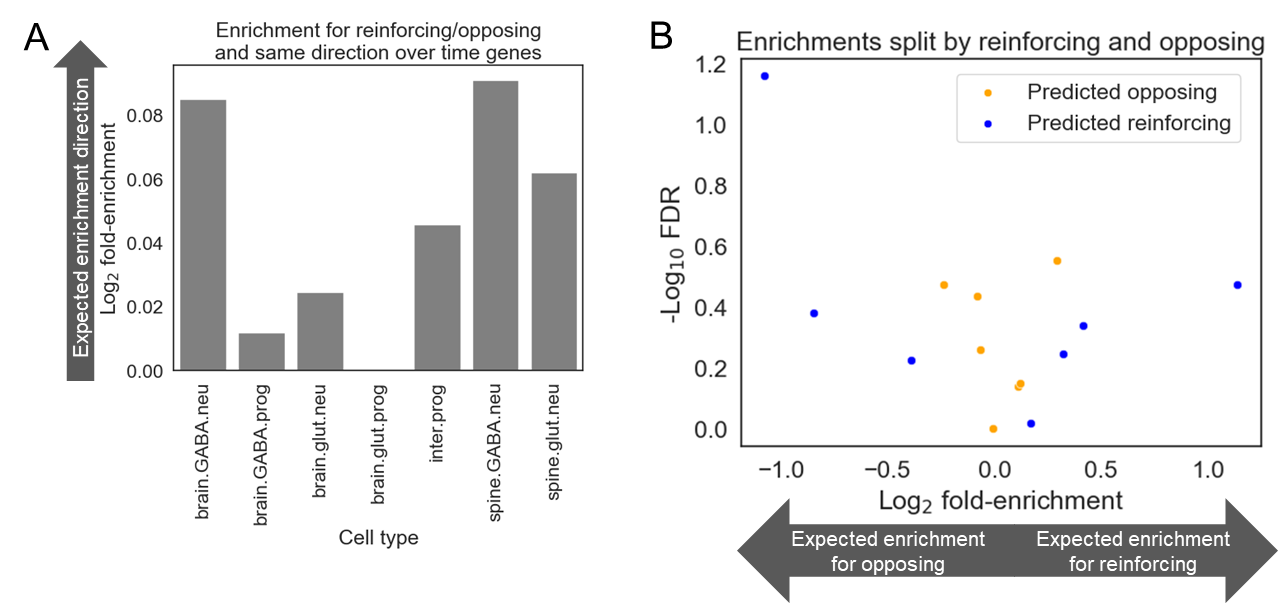

**Fig. S9: Enrichment analysis for genes with conserved gene expression trajectories and reinforcing/opposing intrinsic and extrinsic divergence.** See Methods for how genes with conserved expression trajectories between species were defined. **A)** Enrichment analysis for genes with conserved expression trajectories between species and the product of intrinsic proportion divergence and extrinsic proportion divergence. For this metric, genes with high values have opposing or reinforcing expression and genes with low values do not. The y-axis shows the log_2_ fold-enrichment and the y-axis corresponds to cell type. The arrow shows the expected enrichment if genes with conserved expression trajectories between mice and rats were inflating estimates of reinforcing/opposing divergence. **B)** Enrichment analysis for genes with conserved expression trajectories between species and the product of signed intrinsic proportion divergence and signed extrinsic proportion divergence. For this metric, larger positive values indicate reinforcing intrinsic and extrinsic divergence, large negative values indicate opposing intrinsic and extrinsic divergence, and values near zero indicate neither reinforcing nor opposing intrinsic and extrinsic divergence. The x-axis is the log_2_ fold-enrichment and the y-axis is the -log_10_(FDR). The arrow shows the expected enrichment if genes with the conserved gene expression trajectories between mice and rats were inflating estimates of reinforcing/opposing intrinsic and extrinsic divergence. Each point corresponds to the enrichment in a central nervous system cell type and each cell type is represented twice, once as a blue dot for genes that are increasing over time in mouse and rat so would be predicted to appear as having reinforcing extrinsic and intrinsic divergence in our study, and once as an orange dot for genes that are decreasing over time in mouse and rat so would be predicted to appear as having opposing extrinsic and intrinsic divergence in our study.

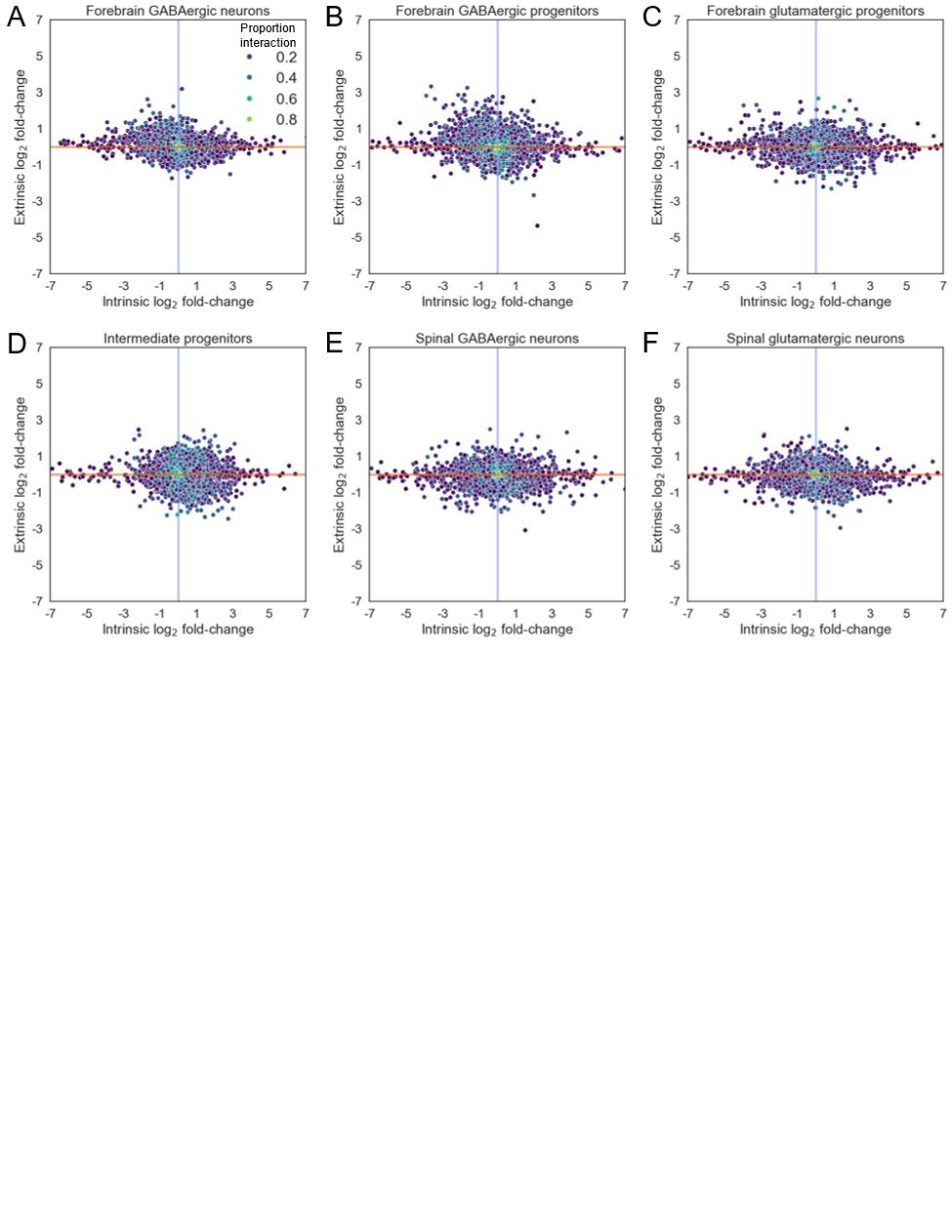

**Fig. S10: Intrinsic and extrinsic divergence of nervous system cell types.** Each point is a gene. Intrinsic divergence is on the y-axis and extrinsic divergence is on the x-axis. Genes are colored by their proportion interaction. Forebrain glutamatergic neurons are shown in Fig. 2F. **A)** Plot for forebrain GABAergic neurons. **B)** Plot for forebrain GABAergic progenitors. **C)** Plot for forebrain glutamatergic progenitors. **D)** Plot for intermediate progenitors. **E)** Plot for spinal GABAergic neurons. **F)** Plot for spinal glutamatergic neurons.

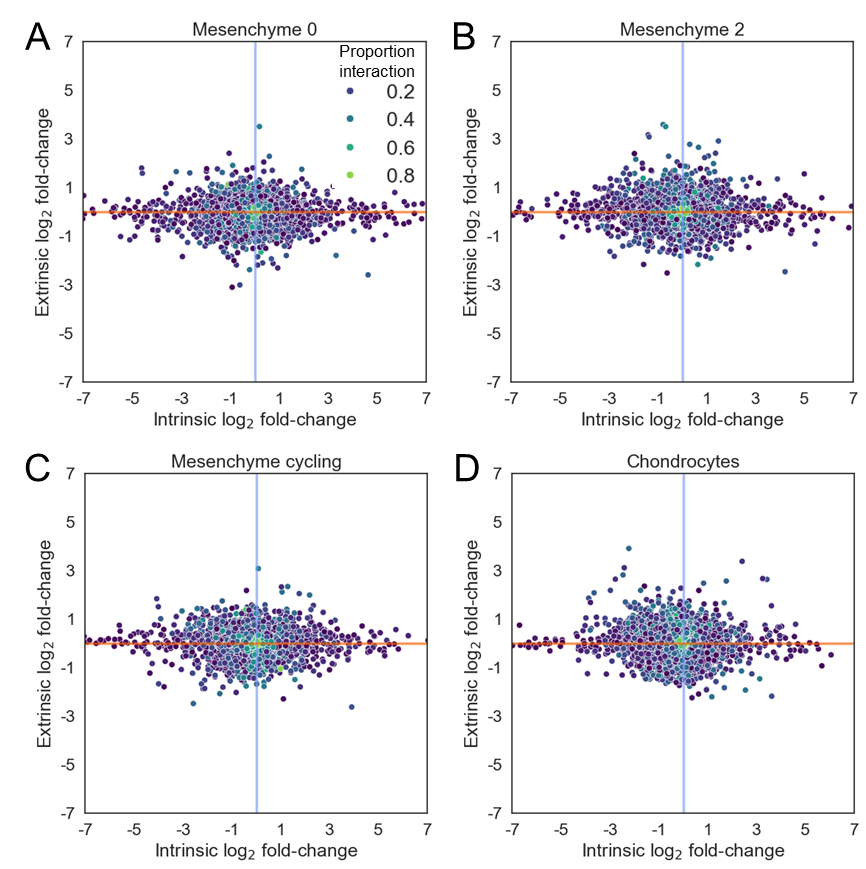

**Fig. S11: Intrinsic and extrinsic divergence of connective tissue cell types.** Each point is a gene. Intrinsic divergence is on the y-axis and extrinsic divergence is on the x-axis. Genes are colored by their proportion interaction. **A)** Plot for mesenchyme cluster 0. **B)** Plot for mesenchyme cluster 2. **C)** Plot for cycling mesenchymal cells. **D)** Plot for chondrocytes.

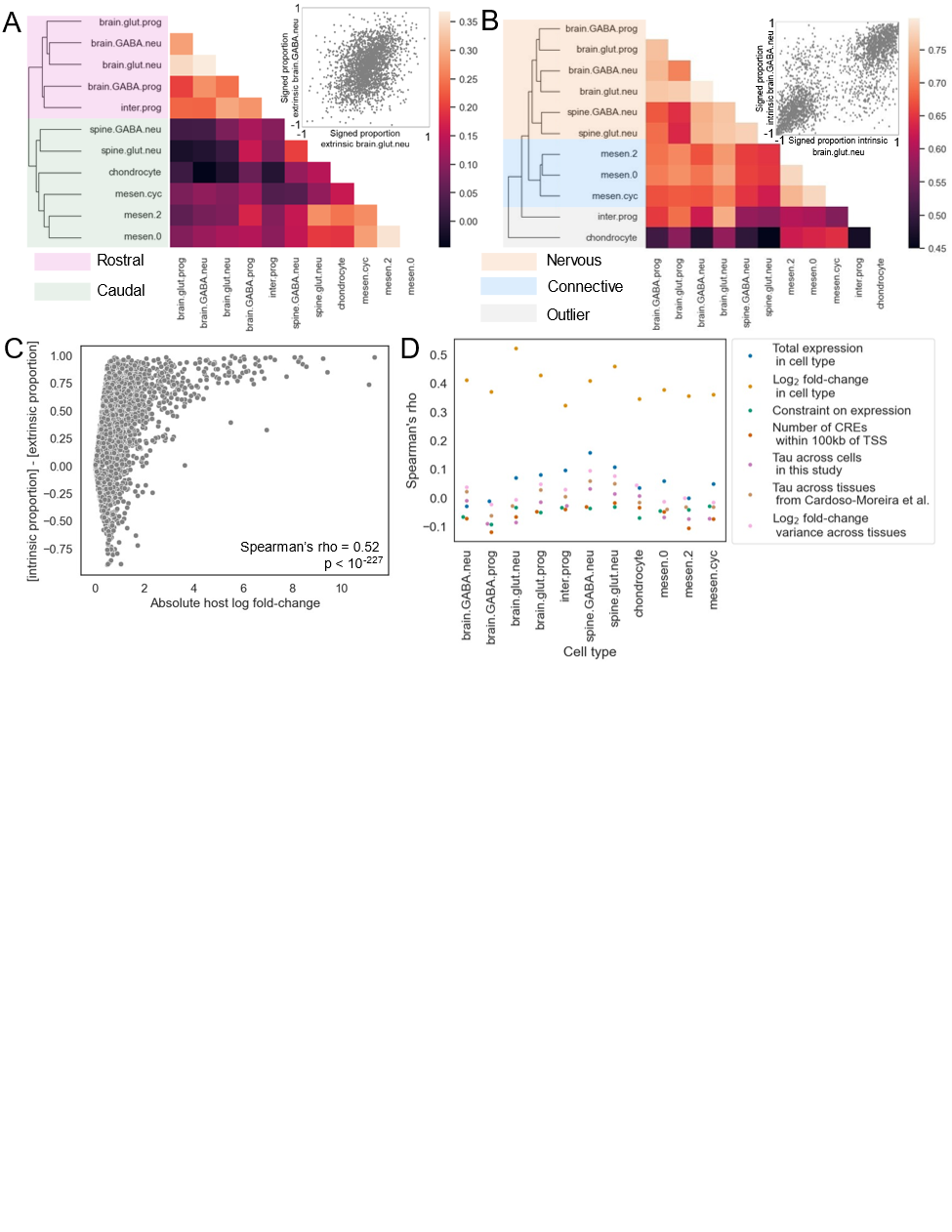

**Fig. S12: Correlates of intrinsic and extrinsic divergence across cell types.** A) Heatmap showing Spearman correlation of signed proportion extrinsic divergence between cell types. Hierarchical clustering was performed on the Spearman rho values using the Euclidean distance metric. Cell types are shaded by their anatomical location of origin, with spinal neurons being more caudal similar to connective tissue cells. For the scatter plot in the upper right, each dot is a gene, the x-axis is the forebrain glutamatergic neuron signed proportion extrinsic divergence, and the y-axis is the forebrain GABAergic neuron signed proportion extrinsic divergence. **B)** The same as in (A) but showing the signed proportion intrinsic divergence and shading cell types by whether they cluster with nervous system cell types, connective tissue cell types, or are outliers. **C)** Scatter plot showing the relationship between absolute log_2_ fold-change between mouse cells in a mouse-like environment and rat cells in a rat-like environment (x-axis) and the proportion extrinsic subtracted from the proportion intrinsic (referred to as [proportion intrinsic - proportion extrinsic], y-axis) for each gene in forebrain glutamatergic neurons. **D)** Spearman correlation coefficients for different per-gene variables and [proportion intrinsic - proportion extrinsic] across cell types.

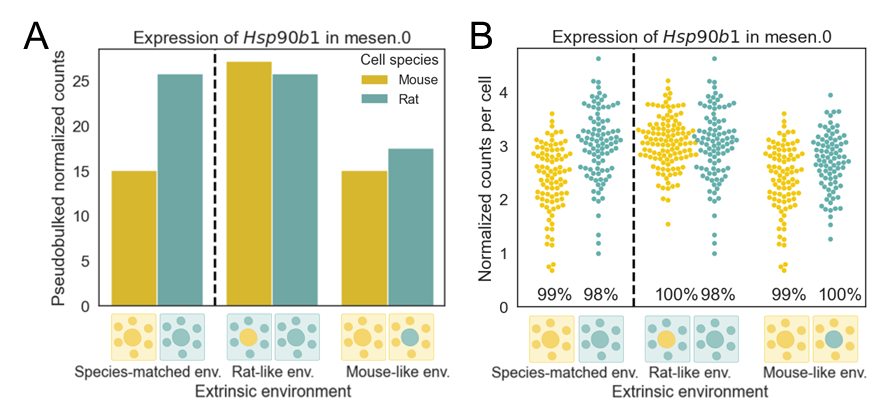

**Fig. S13: Expression of *Hsp90b1*. A)** Expression of *Hsp90b1*, a gene involved in the ER stress response, in mesenchymal cluster 0. **B)** Per-cell expression of *Hsp90b1* in in mesenchymal cluster 0. Each swarm of points shows the normalized counts for a gene in each cell with non-zero counts for that gene. The percentage near the bottom of the plot indicates the percentage of cells with non-zero counts for that gene.

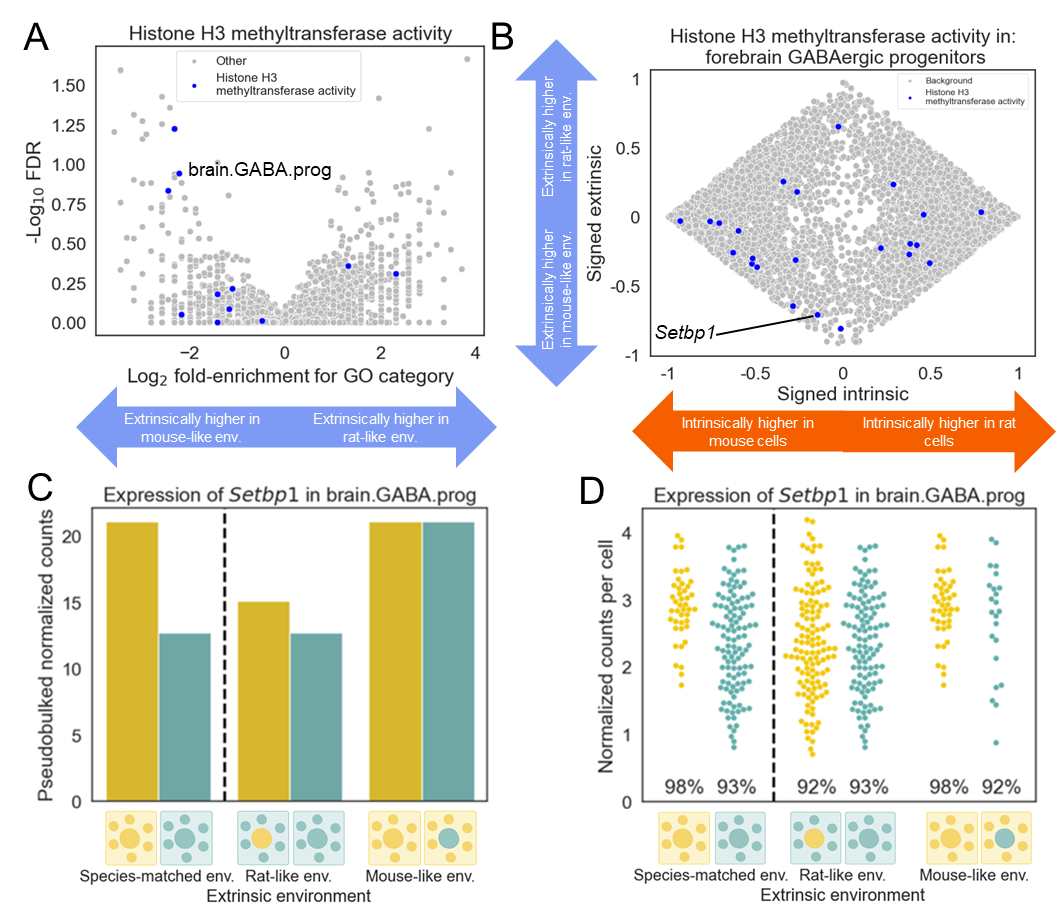

**Fig. S14: Extrinsic divergence of genes encoding histone methyltransferases in central nervous system cell types. A)** Enrichment of genes encoding histone H3 methyltransferase for signed extrinsic divergence across cell types. Each point is a GO biological process category in a cell type and the points corresponding to the histone H3 methyltransferase activity GO category are colored blue. The x-axis shows the log­_2_ fold-enrichment and the y-axis shows the -log_10_ false discovery rate. **B)** Scatterplot showing signed proportion intrinsic divergence (x-axis) and signed proportion extrinsic divergence (y-axis) for all genes passing our filtering criteria for forebrain GABAergic progenitors. Genes coding for histone H3 methyltransferases are shown in blue and all other genes are shown in grey. **C)** Expression of *Setbp1*, a gene involved in histone methylation, in forebrain GABAergic progenitors. **D)** Per-cell expression *Setbp1* in forebrain GABAergic progenitors. Each swarm of points shows the normalized counts for a gene in each cell with non-zero counts for that gene. The percentage near the bottom of the plot indicates the percentage of cells with non-zero counts for that gene.

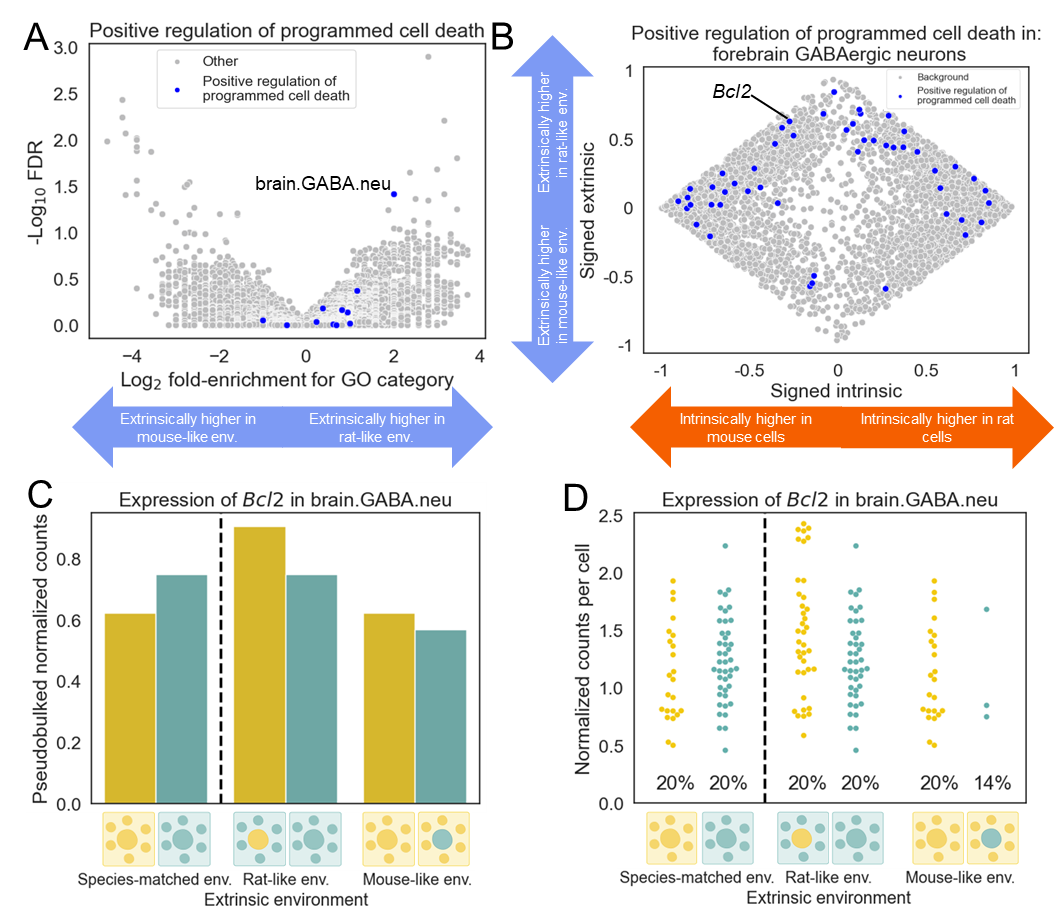

**Fig. S15: Extrinsic divergence of genes involved in the positive regulation of programmed cell death in forebrain GABAergic neurons. A)** Enrichment of genes involved in the positive regulation of programmed cell death for signed extrinsic divergence across cell types. Each point is a GO biological process category in a cell type and the points corresponding to the positive regulation of programmed cell death GO category are colored blue. The x-axis shows the log­_2_ fold-enrichment and the y-axis shows the -log_10_ false discovery rate. **B)** Scatterplot showing signed proportion intrinsic divergence (x-axis) and signed proportion extrinsic divergence (y-axis) for all genes passing our filtering criteria for forebrain GABAergic neurons. Genes involved in the positive regulation of programmed cell death are shown in blue and all other genes are shown in grey. **C)** Expression of *Bcl2*, a gene involved in the regulation of programmed cell death, in forebrain GABAergic neurons. **D)** Per-cell expression *Bcl2* in forebrain GABAergic neurons. Each swarm of points shows the normalized counts for a gene in each cell with non-zero counts for that gene. The percentage near the bottom of the plot indicates the percentage of cells with non-zero counts for that gene.

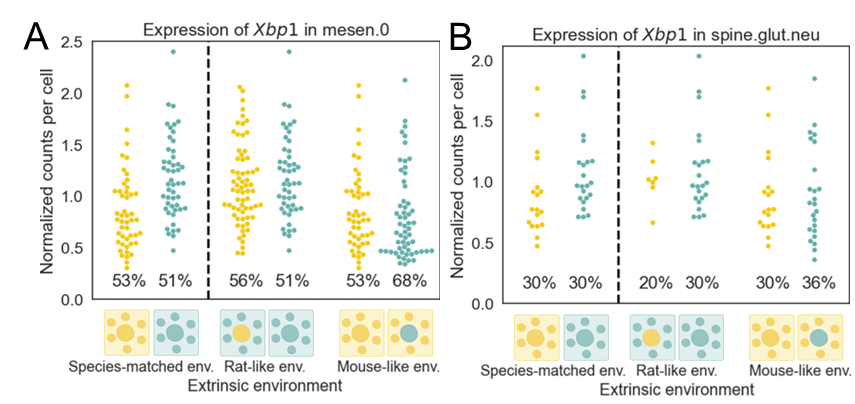

**Fig. S16: Per-cell expression of *Xbp1*.** Each swarm of points shows the normalized counts for a gene in each cell with non-zero counts for that gene. The percentage near the bottom of the plot indicates the percentage of cells with non-zero counts for that gene. **A)** Per-cell expression *Xbp1* in mesenchymal cluster 0. **B)** Per-cell expression of *Xbp1* in spinal glutamatergic neurons.

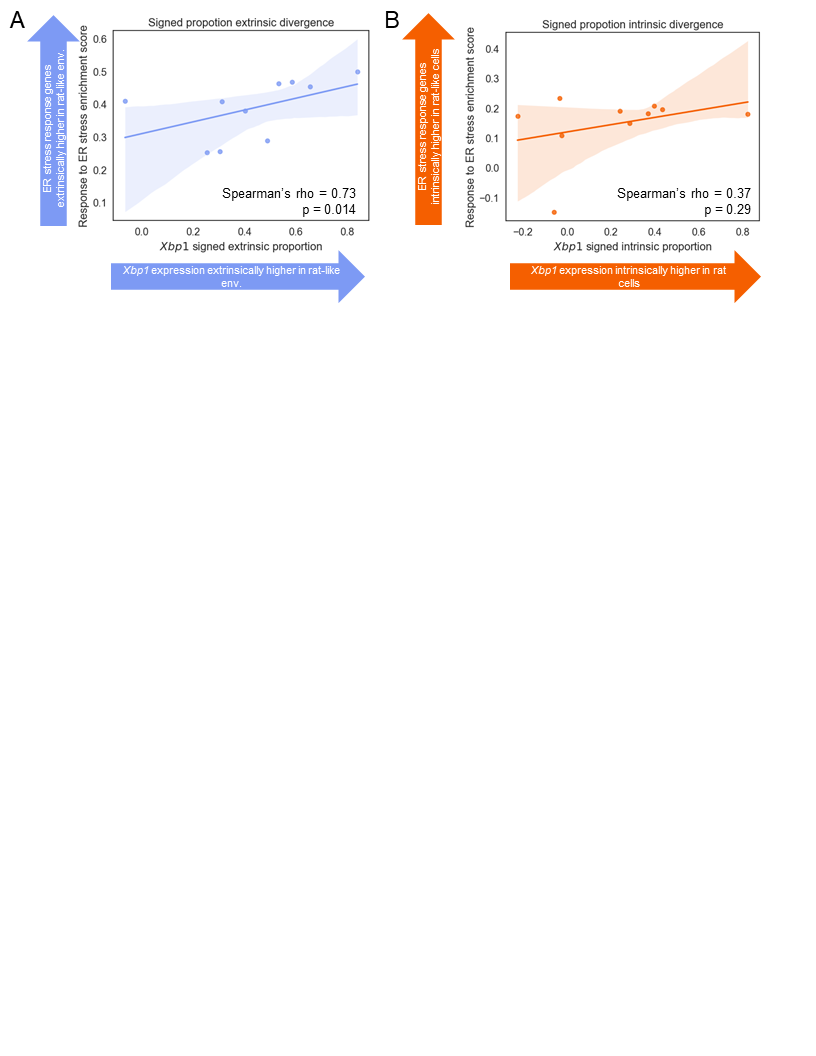

**Fig. S17: Relationship between divergence in *Xbp1* expression and divergence in ER stress response gene expression. A)** Plot showing the relationship between signed proportion extrinsic divergence for *Xbp1* (x-axis) and the GSEA preranked enrichment score for the Response to ER stress GO category and signed proportion extrinsic divergence (y-axis). Each point is a cell type and cell types with extrinsically driven increased expression of *Xbp1* in a rat-like environment have larger values on the x-axis. Cell types with extrinsically driven increased expression of ER stress response genes in a rat-like environment have larger values on the y-axis. The line and shaded region represent the best fit and 95% confidence interval of a linear model fit to the data. **B)** Plot showing the relationship between signed proportion intrinsic divergence for *Xbp1* (x-axis) and the GSEA preranked enrichment score for the Response to ER stress GO category and signed intrinsic proportion divergence (y-axis). Each point is a cell type and cell types with intrinsically driven increased expression of *Xbp1* in rat cells have larger values on the x-axis. Cell types with intrinsically driven increased expression of ER stress response genes in rat cells have larger values on the y-axis. The line and shaded region represent the best fit and 95% confidence interval of a linear model fit to the data.

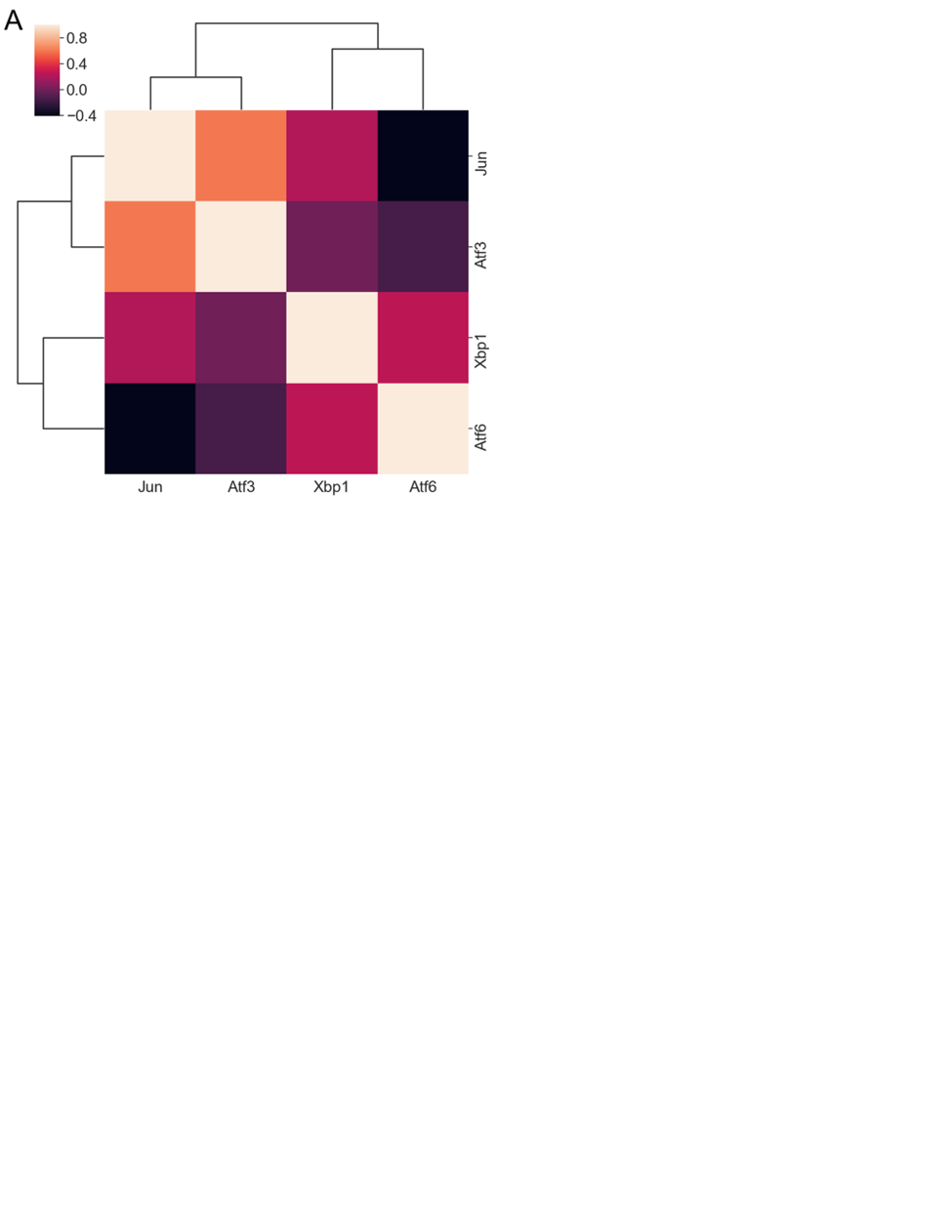

**Fig. S18: Correlation between ER stress response TF signed proportion extrinsic divergence. A)** Heatmap showing the correlation between the signed proportion extrinsic divergence of TFs associated with the ER stress response. The genes were hierarchically clustered using the Euclidean distance metric. Only cell types in which both genes passed our filtering criteria were used to compute the correlation (see Methods).

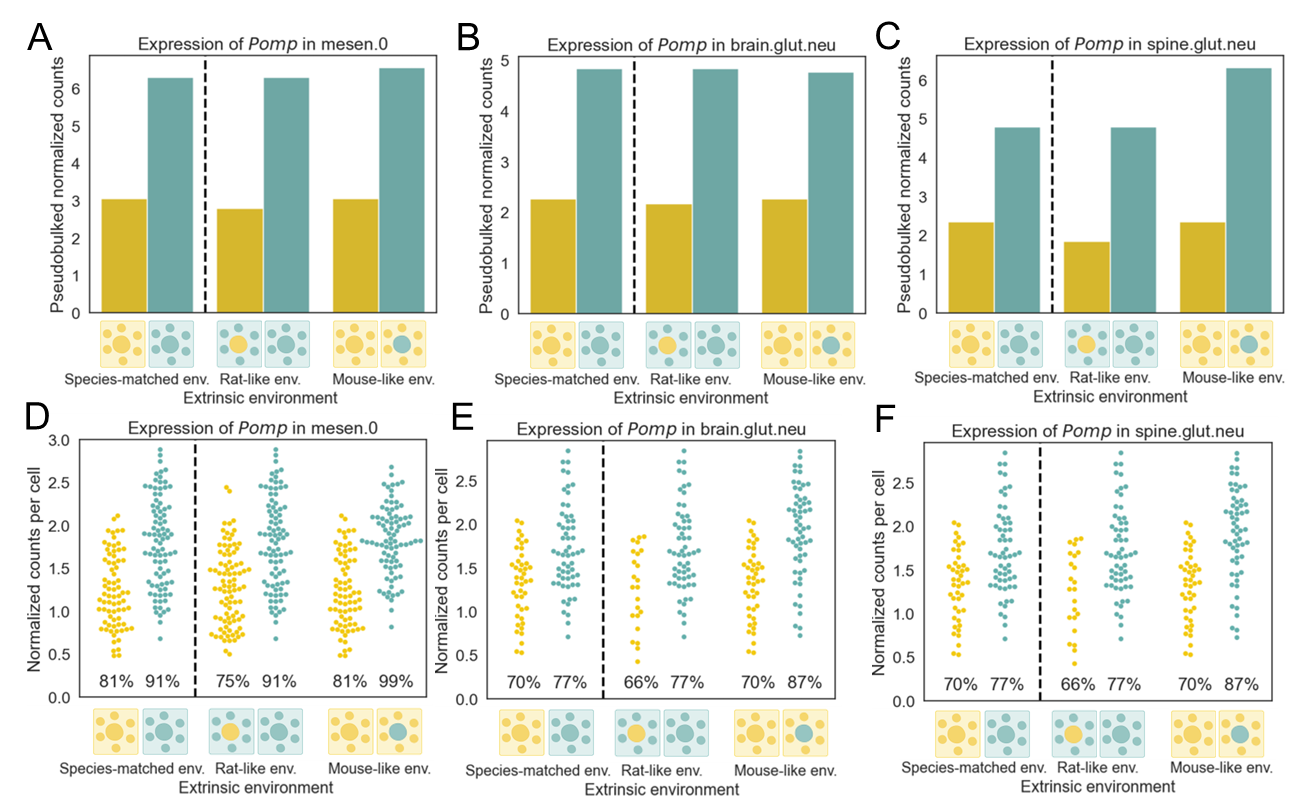

**Fig. S19: Expression of *Pomp* across cell types. A)** Expression of the *Nfe2l1* target gene *Pomp* in mesenchymal cluster 0 cells. Expression is intrinsically higher in rat cells. **B)** Expression of *Pomp* in forebrain glutamatergic neurons. Expression is intrinsically higher in rat cells. **C)** Expression of *Pomp* in spinal glutamatergic neurons. Expression is primarily intrinsically higher in rat cells. **D)** Per-cell expression *Pomp* in mesenchymal cluster 0. Each swarm of points shows the normalized counts for a gene in each cell with non-zero counts for that gene. The percentage near the bottom of the plot indicates the percentage of cells with non-zero counts for that gene. **E)** Per-cell expression of *Pomp* in forebrain glutamatergic neurons. **F)** Per-cell expression of *Pomp* in spinal glutamatergic neurons.

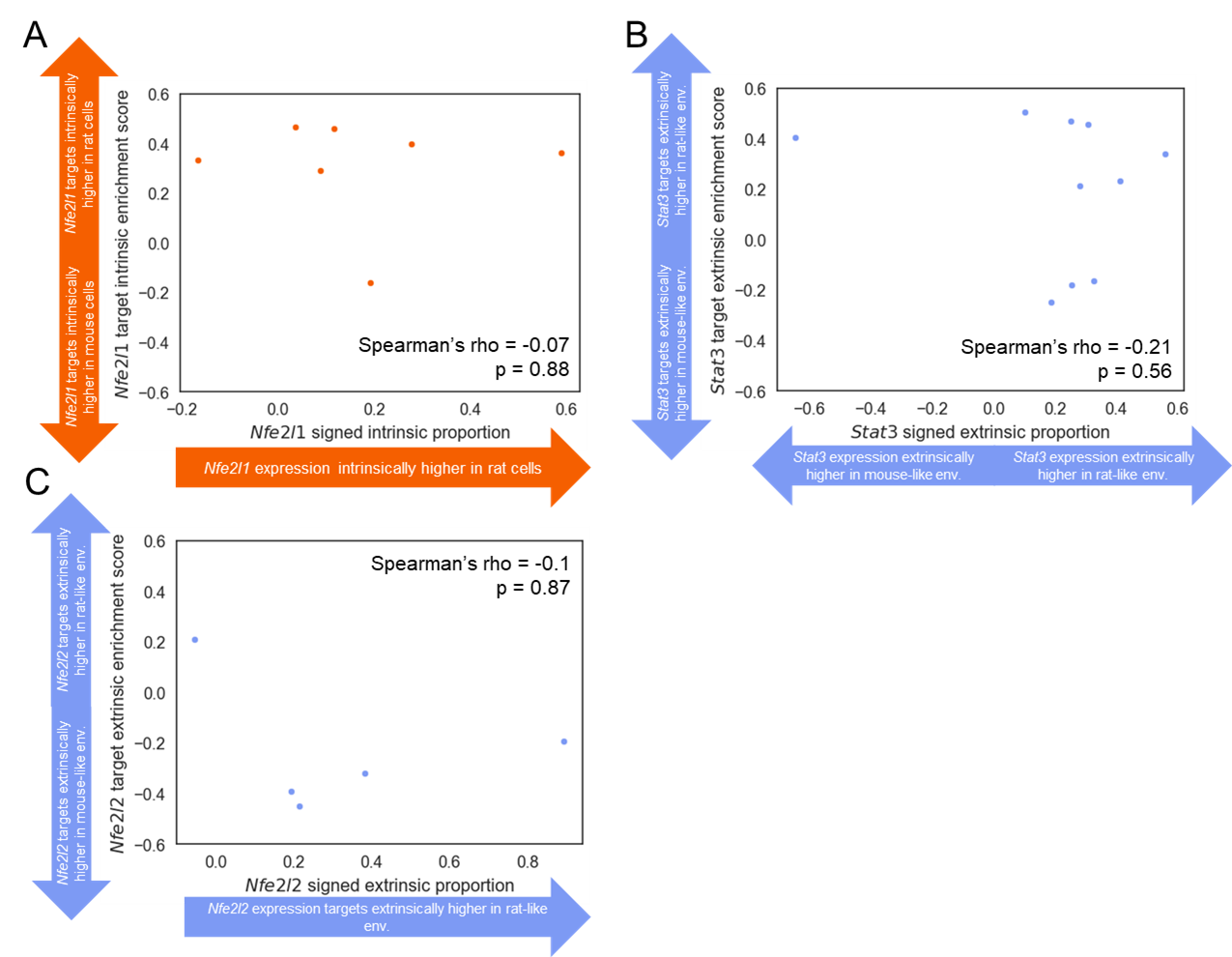
**Fig. S20: Analysis of TFs regulating proteasomal subunit expression and their target genes. A)** Scatter plot showing the relationship between signed intrinsic proportion divergence for *Nfe2l1* (x-axis) and the enrichment of its target genes for signed intrinsic proportion divergence (y-axis). Each point is a cell type and cell types with intrinsically driven increased expression of *Nfe2l1* in rat cells have larger values on the x-axis. Cell types with intrinsically driven increased expression of *Nfe2l1* target genes in rat cells have larger values on the y-axis. **B)** Correlation between signed extrinsic proportion divergence for *Stat3* (x-axis) and the enrichment of its target genes for signed extrinsic proportion divergence (y-axis). Each point is a cell type and cell types with extrinsically driven increased expression of *Stat3* in a rat-like environment have larger values on the x-axis and cell types with extrinsically driven increased expression of *Stat3* in a mouse-like environment have smaller values on the x-axis. Cell types with extrinsically driven increased expression of *Stat3* target genes in a rat-like environment have larger values on the y-axis whereas cell types with extrinsically driven increased expression of *Stat3* target genes in a mouse-like environment have smaller values on the y-axis. **C)** Correlation between signed extrinsic proportion divergence for *Nfe2l2* (x-axis) and the enrichment of its target genes for signed extrinsic proportion divergence (y-axis). Each point is a cell type and cell types with extrinsically driven increased expression of *Nfe2l2* in a rat-like environment have larger values on the x-axis and cell types with extrinsically driven increased expression of *Nfe2l2* in a mouse-like environment have smaller values on the x-axis. Cell types with extrinsically driven increased expression of *Nfe2l2* target genes in a rat-like environment have larger values on the y-axis whereas cell types with extrinsically driven increased expression of *Nfe2l2* target genes in a mouse-like environment have smaller values on the y-axis.

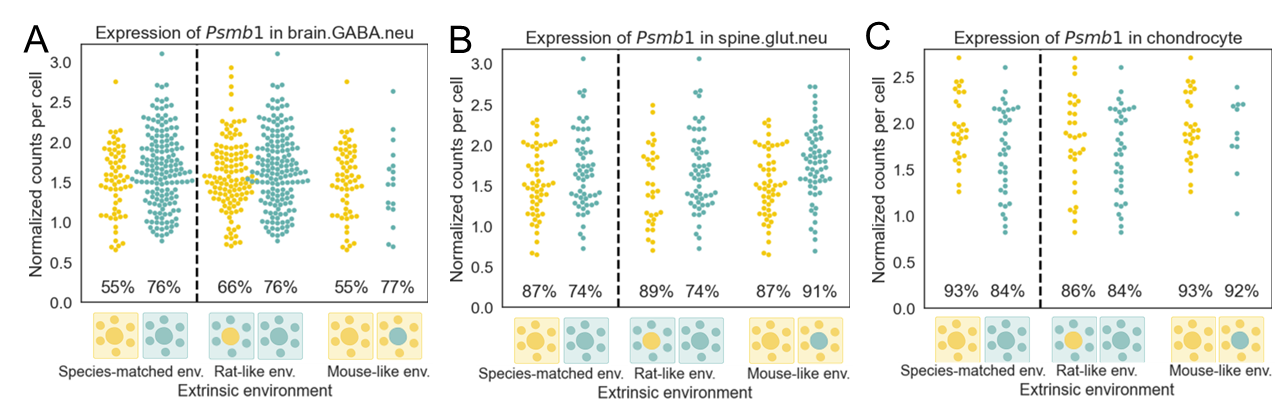

**Fig. S21: Per-cell expression of *Psmb1* across cell types.** Each swarm of points shows the normalized counts for a gene in each cell with non-zero counts for that gene. The percentage near the bottom of the plot indicates the percentage of cells with non-zero counts for that gene. **A)** Per-cell expression *Psmb1* in forebrain GABAergic neurons. **B)** Per-cell expression of *Psmb1* in spinal glutamatergic neurons. **C)** Per-cell expression of *Psmb1* in chondrocytes.

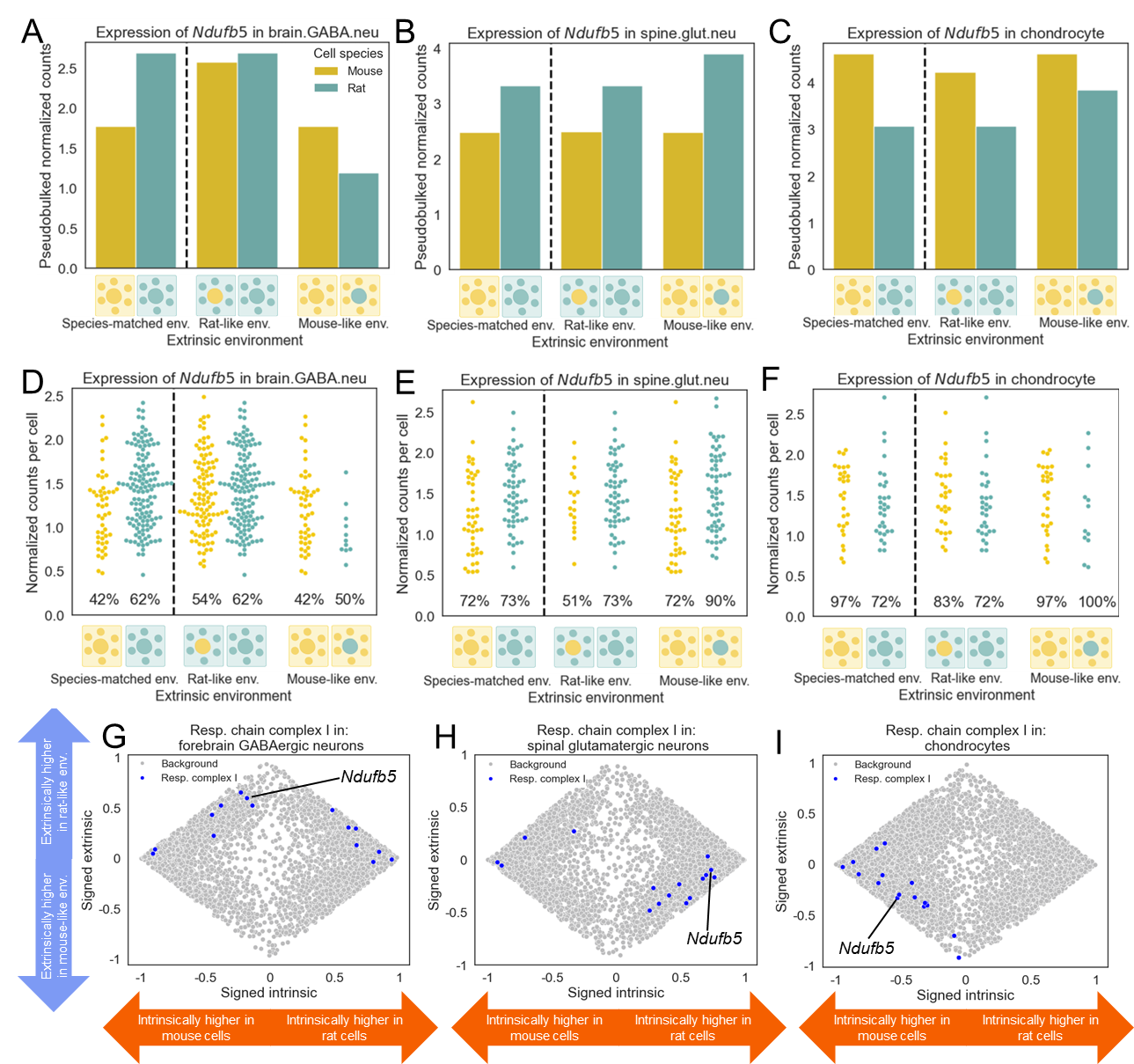

**Fig. S22: Cell-type specific intrinsic and extrinsic divergence of the expression of genes encoding mitochondrial respiratory chain complex I subunits across cell types. A)** Expression of *Ndufb5* in forebrain GABAergic neurons. **B)** Same as in (A) but for spinal glutamatergic neurons. **C)** Same as in (A) but for chondrocytes. **D)** Per-cell expression *Ndufb5* in forebrain GABAergic neurons. Each swarm of points shows the normalized counts for a gene in each cell with non-zero counts for that gene. The percentage near the bottom of the plot indicates the percentage of cells with non-zero counts for that gene. **E)** Per-cell expression of *Ndufb5* in spinal glutamatergic neurons. **F)** Per-cell expression of *Ndufb5* in chondrocytes. **G)** Scatterplot showing signed proportion intrinsic divergence (x-axis) and signed proportion extrinsic divergence (y-axis) for all genes passing our filtering criteria for forebrain GABAergic neurons. Genes coding for mitochondrial respiratory chain complex I subunits are shown in blue and all other genes are shown in grey. Expression of genes coding for mitochondrial respiratory chain complex I subunits is generally extrinsically higher in a rat-like environment. **H)** Same as in (G) but for spinal glutamatergic neurons. Expression of genes coding for mitochondrial respiratory chain complex I subunits is generally intrinsically higher in rat cells but extrinsically higher in a mouse-like environment. **I)** Same as in (G) but for chondrocytes. Expression of genes coding for mitochondrial respiratory chain complex I subunits is generally intrinsically higher in mouse cells and extrinsically higher in a mouse-like environment.

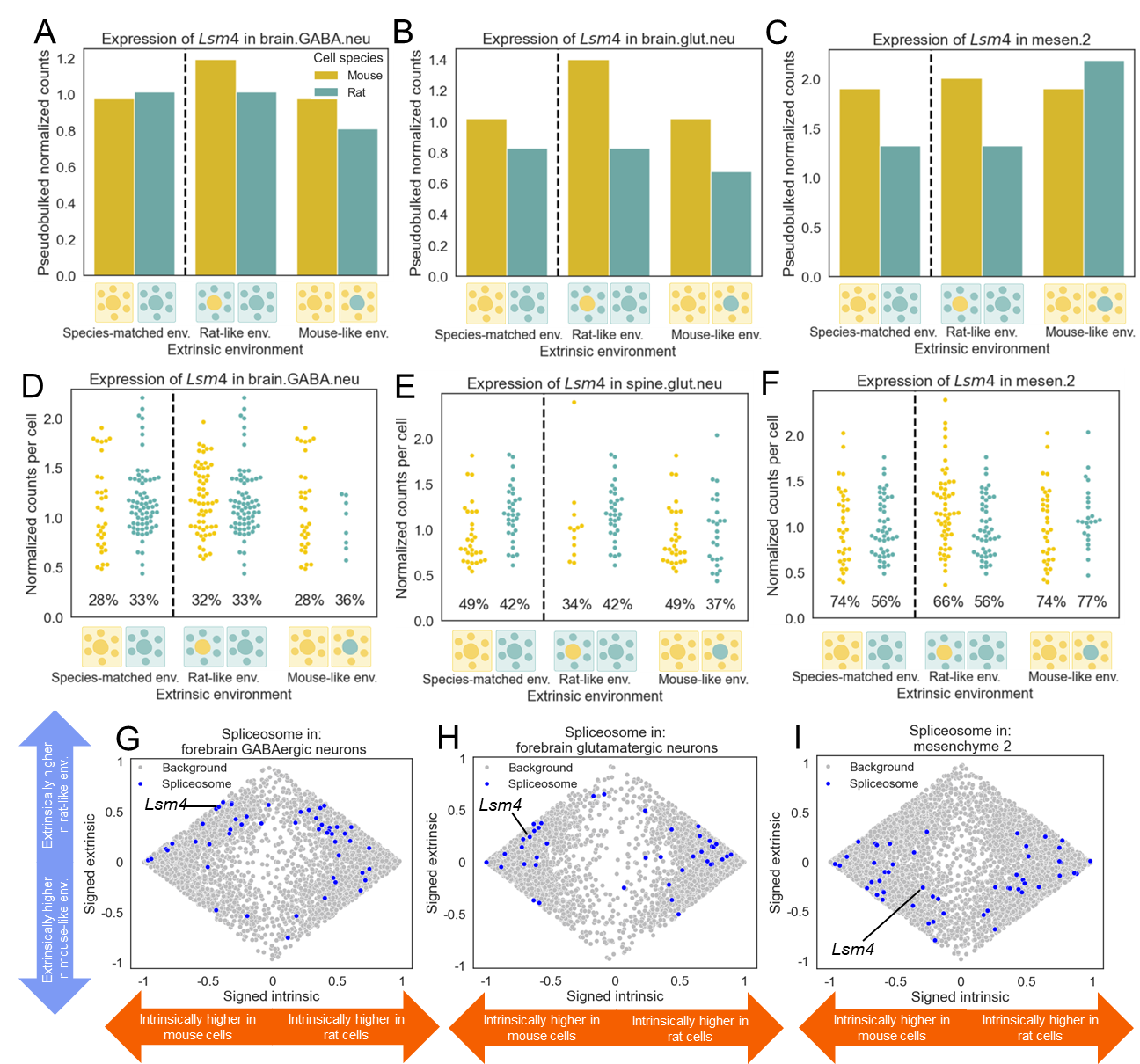

**Fig. S23: Cell-type specific intrinsic and extrinsic divergence of the expression of genes encoding spliceosomal subunits across cell types. A)** Expression of *Lsm4* in forebrain GABAergic neurons. **B)** Same as in (A) but for spinal glutamatergic neurons. **C)** Same as in (A) but for mesenchymal cluster 2 cells. **D)** Per-cell expression *Lsm4* in forebrain GABAergic neurons. Each swarm of points shows the normalized counts for a gene in each cell with non-zero counts for that gene. The percentage near the bottom of the plot indicates the percentage of cells with non-zero counts for that gene. **E)** Per-cell expression of *Lsm4* in spinal glutamatergic neurons. **F)** Per-cell expression of *Lsm4* in mesenchymal cluster 2 cells. **G)** Scatterplot showing signed proportion intrinsic divergence (x-axis) and signed proportion extrinsic divergence (y-axis) for all genes passing our filtering criteria for forebrain GABAergic neurons. Genes coding for spliceosomal subunits are shown in blue and all other genes are shown in grey. Expression of genes coding for spliceosomal subunits is generally extrinsically higher in a rat-like environment. **H)** Same as in (G) but for forebrain glutamatergic neurons. Expression of genes coding for spliceosomal subunits is generally extrinsically higher in a mouse-like environment and intrinsically higher in rat cells. **I)** Same as in (G) but for mesenchymal cluster 2 cells. Expression of genes coding for spliceosomal subunits is generally extrinsically higher in a mouse-like environment and intrinsically higher in mouse cells.

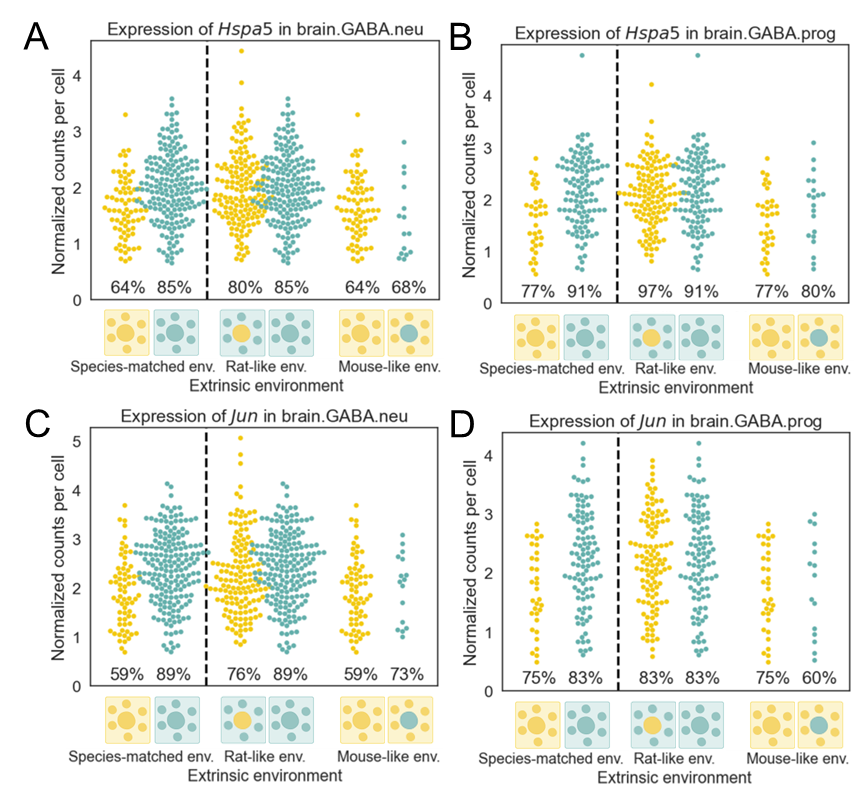

**Fig. S24: Per-cell expression of *Jun* and *Hspa5* in forebrain GABAergic neurons and progenitors.** Each swarm of points shows the normalized counts for a gene in each cell with non-zero counts for that gene. The percentage near the bottom of the plot indicates the percentage of cells with non-zero counts for that gene. **A)** Per-cell expression of *Hspa5* in forebrain GABAergic neurons. **B)** Per-cell expression of *Hspa5* in forebrain GABAergic progenitors. **C)** Per-cell expression of *Jun* in forebrain GABAergic neurons. **D)** Per-cell expression of *Jun* in forebrain GABAergic progenitors.

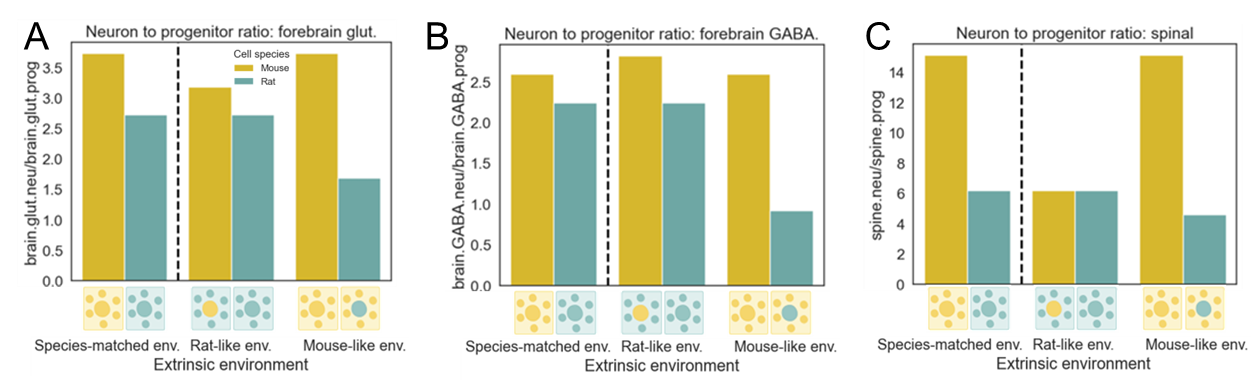

**Fig. S25: Neuron-to-progenitor ratio in different neurogenic niches.** **A)** Plot showing the neuron-to-progenitor ratio for forebrain glutamatergic neurogenesis across the four different species-environment combinations. **B)** Same as in (A) but for forebrain GABAergic neurogenesis. **C)** Same as in (A) but for combined GABAergic and glutamatergic spinal neurogenesis. The two lineages were combined as we were unable to distinguish GABAergic and glutamatergic progenitors.

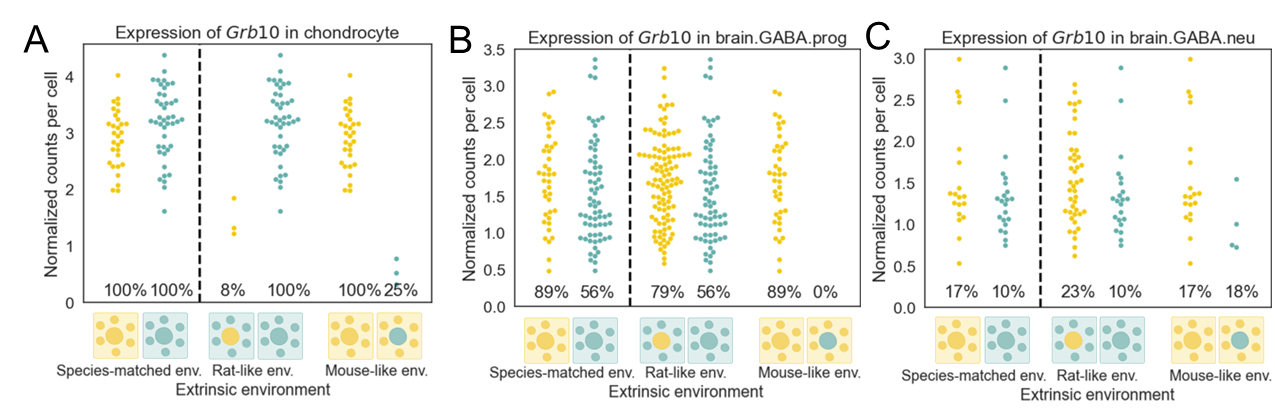

**Fig. S26: Per-cell expression of *Grb10* across cell types.** Each swarm of points shows the normalized counts for a gene in each cell with non-zero counts for that gene. The percentage near the bottom of the plot indicates the percentage of cells with non-zero counts for that gene. **A)** Per-cell expression of *Grb10* in chondrocytes. **B)** Per-cell expression of *Grb10* in forebrain GABAergic progenitors. **C)** Per-cell expression of *Grb10* in forebrain GABAergic neurons.

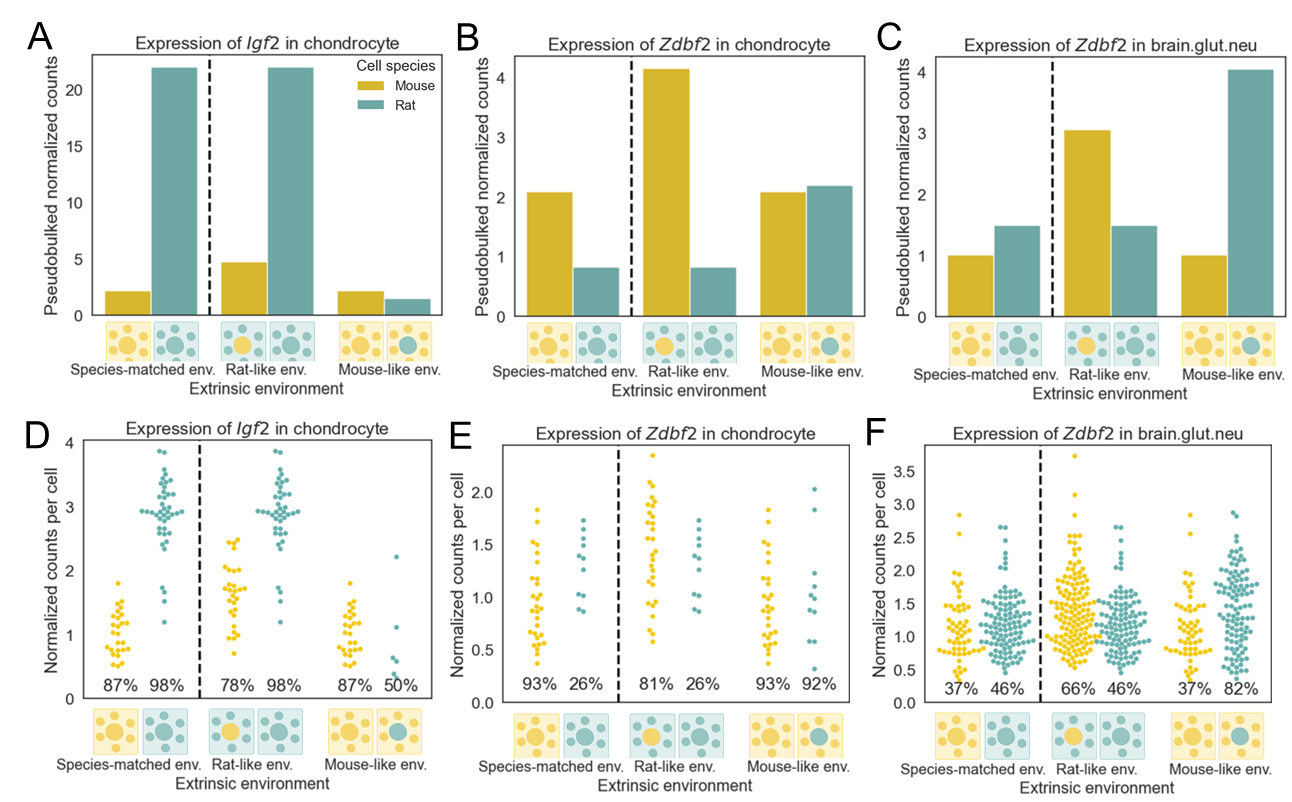

**Fig. S27: Expression of selected imprinted genes. A)** Expression of *Igf2* across the four species-environment combinations in chondrocytes. Expression is very high for rat cells in a rat-like environment but is very low for rat cells in a mouse-like environment. **B)** Expression of the imprinted gene *Zdbf2* across the four species-environment combinations in chondrocytes. Expression is higher in species-mismatched environments in cells from both species. **C)** Expression of the imprinted gene *Zdbf2* in forebrain glutamatergic neurons across the four species-environment combinations in chondrocytes. Expression is higher in species-mismatched environments in cells from both species. **D)** Per-cell expression *Igf2* in chondrocytes. Each swarm of points shows the normalized counts for a gene in each cell with non-zero counts for that gene. The percentage near the bottom of the plot indicates the percentage of cells with non-zero counts for that gene. **E)** Per-cell expression of *Zdbf2* in chondrocytes. **F)** Per-cell expression of *Zdbf2* in forebrain glutamatergic neurons.

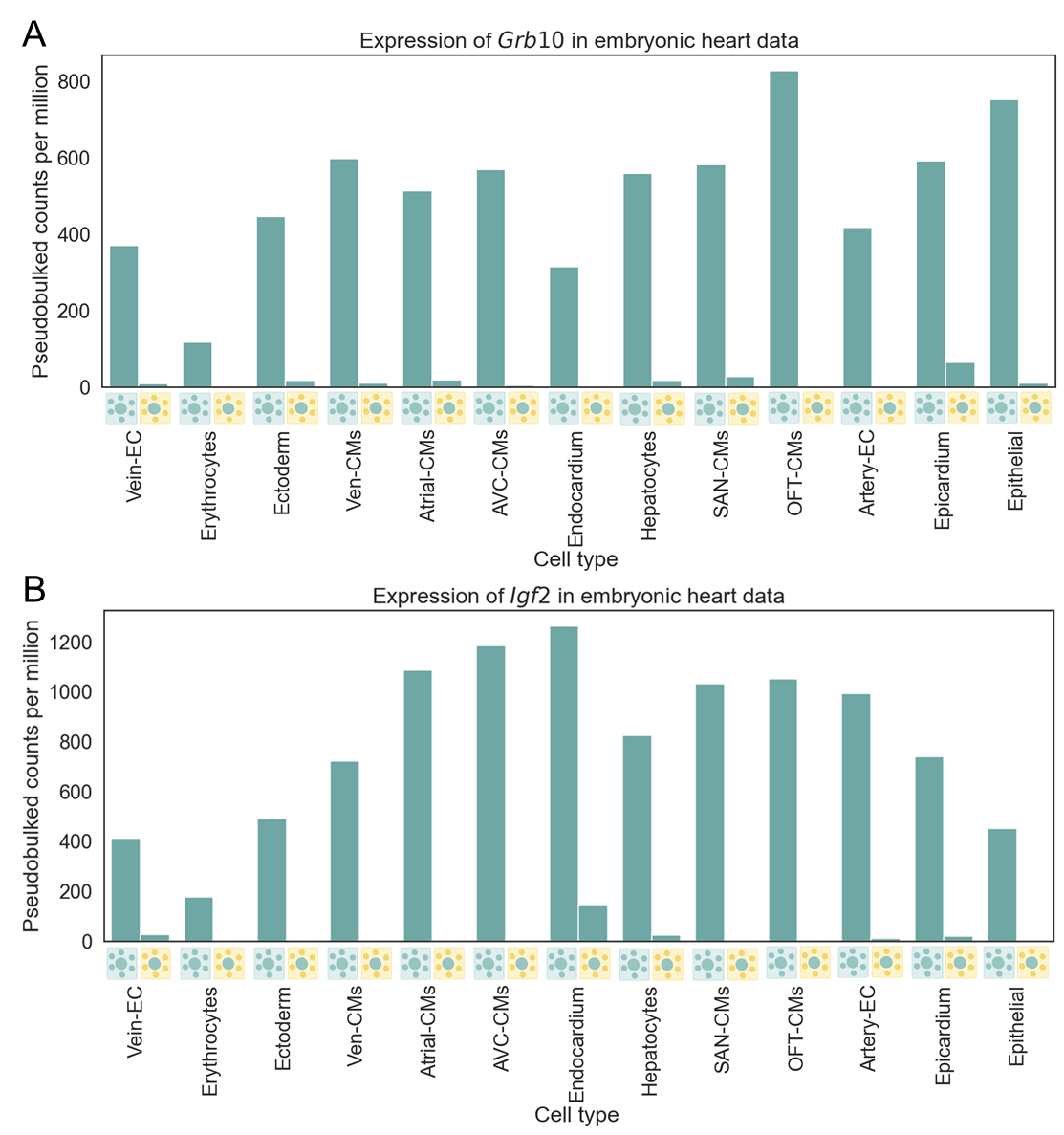

**Fig. S28: Expression of imprinted genes *Grb10* and *Igf2* across cell types from the heart dataset.** For each cell type, expression in rat cells in a rat-like environment is shown on the left and expression in rat cells in a mouse-like environment is shown on the right. **A)** Expression of *Grb10* across cell types. Expression is higher in a species-matched environment. **B)** Expression of *Igf2* across cell types. Expression is higher in a species-matched environment.

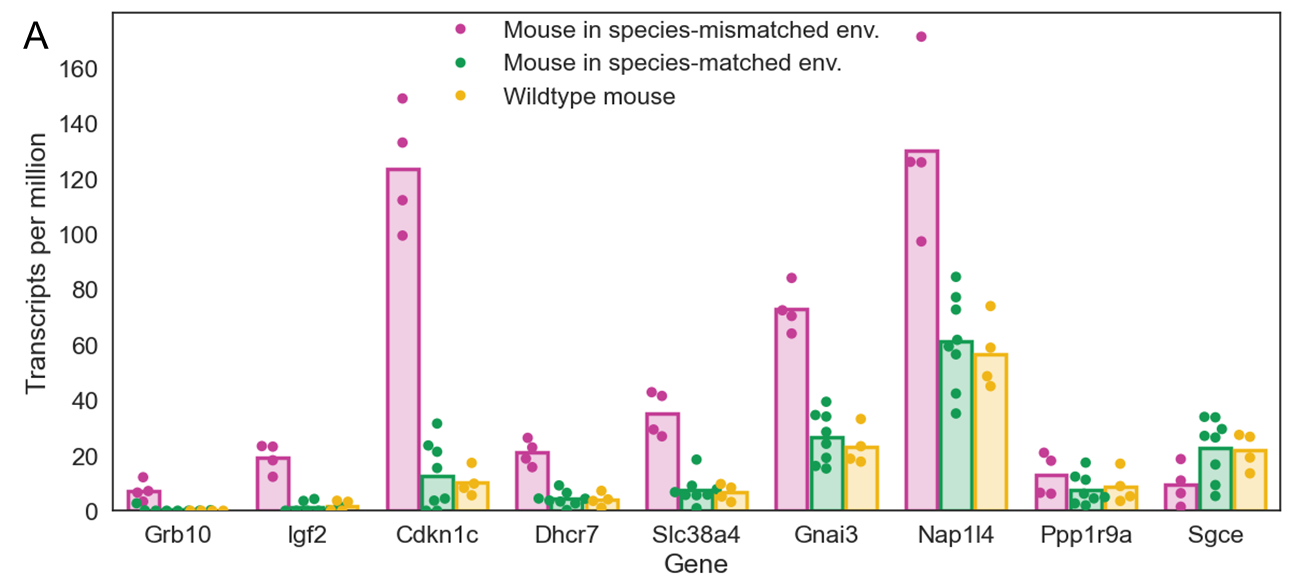

**Fig. S29: Disrupted expression of imprinted genes in adult mouse parathyroid cells in species-mismatched environments. A)** Expression levels of mis-expressed imprinted genes for samples from species-mismatched environments, species-matched environments, and wildtype mice. Expression is higher in species-mismatched environments, but very similar between donor mouse cells in species-matched environments and wildtype mice. All imprinted genes with absolute difference between expression for species-mismatched and species-matched environment samples greater than 1 were plotted.
